# GBP2 aggregates LPS and activates the caspase-4 inflammasome independent of the bacterial encapsulation factor GBP1

**DOI:** 10.1101/2022.10.05.511023

**Authors:** Mary S. Dickinson, Miriam Kutsch, Linda Sistemich, Dulcemaria Hernandez, Anthony S. Piro, David Needham, Cammie F. Lesser, Christian Herrmann, Jörn Coers

**Author notes:** corresponding author: Jörn Coers. authors contributed equally (Mary S. Dickinson, Miriam Kutsch). **Author Contributions:**MSD, MK, JC designed research; MSD, MK, LS performed research; DH, ASP, DN, CFL, CH contributed new reagents/ analytic tools; MSD, MK, LS, DN, CH, JC analyzed data; MSD, MK, JC wrote paper, MK, CFL, CH, JC obtained funding. **Competing Interest Statement:**The authors declare no competing interests.

## Abstract

Sensing and killing of intracellular bacterial pathogens are important features of cell-autonomous immunity. The cytokine gamma-interferon (IFNγ) enhances cell-autonomous immunity through upregulation of interferon stimulated genes such as guanylate binding proteins (GBPs). GBPs promote defense against Gram-negative cytosolic bacteria in part through the induction of an inflammatory cell death pathway called pyroptosis. To activate pyroptosis, GBPs facilitate caspase-4 sensing of the Gram-negative bacterial outer membrane component lipopolysaccharide (LPS). There are seven human GBP paralogs and it is unclear how each GBP contributes to LPS sensing and pyroptosis induction. GBP1 forms a multimeric microcapsule on the surface of cytosolic bacteria through direct interactions with LPS and recruits caspase-4 to bacteria, a process deemed essential for caspase-4 activation. In contrast to GBP1, closely related paralog GBP2 is unable to bind bacteria on its own but requires GBP1 for direct bacterial binding. Unexpectedly, we find that GBP2 overexpression can restore Gram-negative-induced pyroptosis in GBP1^KO^ cells, without GBP2 binding to the bacterial surface. A mutant of GBP1 that lacks the triple arginine motif required for microcapsule formation also rescues pyroptosis in GBP1^KO^ cells, showing that binding to bacteria is dispensable for GBPs to promote pyroptosis. Instead, we find that GBP2, like GBP1, directly binds and aggregates ‘free’ LPS through protein polymerization. This provides a novel mechanistic framework for non-canonical inflammasome activation where GBP1 or GBP2 assemble cytosol-contaminating LPS into a protein-LPS interface for caspase-4 activation as part of a coordinated host response to Gram-negative bacterial infections.

**Significance Statement:** Sensing Gram-negative bacterial lipopolysaccharide by human caspase-4 is critical for host defense to intracellular Gram-negative bacterial pathogens. Human guanylate binding proteins (GBPs) facilitate caspase-4 activation in response to Gram-negative infections by a poorly understood mechanism. The prevailing model suggests GBP1 binding to bacteria and consequential recruitment of caspase-4 to the bacterial surface are essential for triggering this host response. Here, we show GBP1 binding to bacteria is dispensable for caspase-4 activation and identify GBP2 as an additional lipopolysaccharide-binding protein that can functionally replace GBP1. We demonstrate that GBP1 and GBP2 share the ability to form lipopolysaccharide-protein complexes, which, we propose, allow caspase-4 activation. Our study provides a new mechanistic framework for cytosolic LPS sensing.

## Introduction

Cell-autonomous immune responses to intracellular pathogens are a powerful defense against infection (1). Within a single cell there are highly orchestrated pathways that sense and respond to invading microbes. During infection with Gram-negative bacteria, immune sensors detect the presence of lipopolysaccharide (LPS), an abundant component of the Gram-negative cell wall. Extracellular LPS is detected through TLR4, causing pro-inflammatory transcriptional changes within the cell, and cytosolic LPS is sensed through the caspase-4 non-canonical inflammasome, triggering inflammatory cell death through pyroptosis (2–8). Activation of caspase-4 not only occurs in professional immune sentinel cells but also in non-immune cells such as colonic epithelial cells, where caspase-4 plays an essential role in host defense against enteric bacterial pathogens (9–11).

Pyroptosis exerts host defense through the destruction of the replicative niche of intracellular bacteria. Additionally, pyroptotic death is proinflammatory and thereby promotes the recruitment and activation of professional immune cells to further restrict infection (12). Pyroptosis can be triggered by diverse stimuli which include cytosolic LPS. Cytosolic LPS binds and activates caspase-4, which then cleaves gasdermin D (GSDMD). The cleaved N-terminus of GSDMD forms a pore in the plasma membrane (13, 14), disrupting the ionic balance of the cell and leading to cell swelling and eventual rupture of the plasma membrane through a programmed cell lysis pathway (15). The GSDMD pore also allows secretion of pro-inflammatory alarmins and cytokines such as IL-18. Due to this inflammatory nature of pyroptosis, professional immune cells are recruited to and activated at the site of infection, where they provide critical immune effector functions. Recruited immune cells release additional cytokines to enhance local responses to pathogens. For example, innate and adaptive lymphocytes activated during a type 1 immune response secrete the potent antibacterial cytokine gamma-interferon (IFNγ). IFNγ induces expression of hundreds of proteins encoded by interferon stimulated genes, including the dynamin-related guanylate binding proteins (GBPs). In both mouse and human cells, GBPs have been shown to promote pyroptosis in response to LPS introduced into the cytosol, either through LPS transfection, uptake of outer membrane vesicles containing LPS, or during Gram-negative bacterial infection (16–22).

Humans have seven GBPs, of which GBP1, GBP2, GBP3, GBP4, and GBP5 are widely expressed in many cell types, and are some of the most highly upregulated genes in response to IFNγ (23). GBP1 is the most well characterized paralog and can target Gram-negative bacteria in the cytosol due to GBP1’s ability to bind LPS (24–26). Following initial docking on the bacterial surface, GBP1 intercalates into the bacterial outer membrane and forms a stable coat called the GBP1 microcapsule (24). GBP1 recruits GBP2, GBP3, and GBP4 to the bacterial surface through unknown mechanisms (27–29). It was proposed that this protein coat consisting of multiple GBP paralogs forms a signaling platform on the surface of bacteria, allowing recruitment and activation of caspase-4 and consequential pyroptosis in human epithelial cells (25, 26). Whether and how each GBP paralog contributes to pyroptosis or restriction of bacterial growth during Gram-negative infection of epithelial cells has not been clearly defined. Similarly, it has remained unclear how the model of a GBP signaling platform on the surface of bacteria relates to GBP-mediated caspase-4 activation following outer membrane vesicle uptake or LPS transfection.

GBP2, the protein with the highest degree of homology to GBP1, is unable to encapsulate bacteria in cells lacking its interaction partner and bacterial encapsulation factor GBP1. Although GBP2 cannot bind to bacteria in the absence of GBP1, we show here that GBP2 overexpression restores pyroptotic cell death in response to Gram-negative infections in GBP1 knockout cells. We demonstrate that GBP2 binds to LPS and forms LPS aggregates, thus sharing critical biochemical properties with GBP1. A GBP1 mutant lacking its unique C-terminal polybasic motif essential for GBP1 microcapsule formation maintains the ability to aggregate LPS and to activate pyroptosis in response to Gram-negative infections or LPS transfections. Our studies therefore demonstrate that GBP1 or GBP2 binding to the bacterial surface is dispensable for caspase-4 activation in response to infections with cytosolic Gram-negative bacteria and instead support a model in which GBP1 and GBP2 facilitate pyroptosis independent of each other through the formation of LPS aggregates.

## Results

### GBP1 targeting of *S. flexneri* does not correlate with pyroptosis

The prevailing model of caspase-4 activation during infections with cytosolic Gram-negative bacteria postulates that GBP1 binding to the bacterial outer membrane recruits GBP2, 3, and 4, and that this complex assembled on the bacterial surface forms an essential platform for caspase-4 activation (25, 26). Following caspase-4 activation, cells undergo pyroptosis and invading bacteria are killed. While GBP1, 2, 3, and 4 are all present on the bacterial surface, it is unclear how each GBP contributes to caspase-4 activation. *Shigella flexneri* is a useful model pathogen for studying GBP regulation of the caspase-4 inflammasome, because it encodes specific effectors that modulate this process; OspC3 blocks caspase-4 activation and IpaH9.8 reduces GBP bacterial targeting through degradation of GBP1, 2, and 4 (10, 27, 28, 30, 31). *S. flexneri* Δ*ospC3* induces pyroptosis that is dependent on caspase-4. However, it is unclear whether IpaH9.8 affects cell death by reducing the targeting of GBPs to the *S. flexneri* surface (10). We previously showed that there is significantly less binding of GBP1 to *S. flexneri* in A549 cells compared to HeLa cells, making these two cell lines useful for testing how different levels of GBP targeting to the *S. flexneri* surface affect cell death or bacterial growth restriction (27). To investigate the role of GBP binding in activation of pyroptosis or bacterial killing, we infected A549 and HeLa cells with four *S. flexneri* strains: wildtype, Δ*ipaH9.8*, Δ*ospC3* and Δ*ospC3*Δ*ipaH9.8*. Since GBP1 binding to *S. flexneri* is required for subsequent recruitment of other GBP family members, we used GBP1 targeting as a proxy for all GBP recruitment (27, 28). Using immunofluorescence microscopy, we again saw that GBP1 targeting of bacteria in A549 cells was significantly lower than in HeLa cells, with 3.9% of *S. flexneri* Δ*ipaH9.8* targeted by GBP1 in A549 cells compared to 32.9% of *S. flexneri* Δ*ipaH9.8* targeted in HeLa cells at 1.5 hours post infection (Fig. 1A). This may be due to lower levels of GBP expression in A549 cells (Fig. 1B). Strains lacking OspC3 had slightly lower levels of targeting than isogenic controls, likely due to cell death or bacterial killing, but knockout (KO) of IpaH9.8 in *S. flexneri* Δ*ospC3* still substantially increased the levels of GBP1 targeting (Fig. 1A).

**Figure 1.**
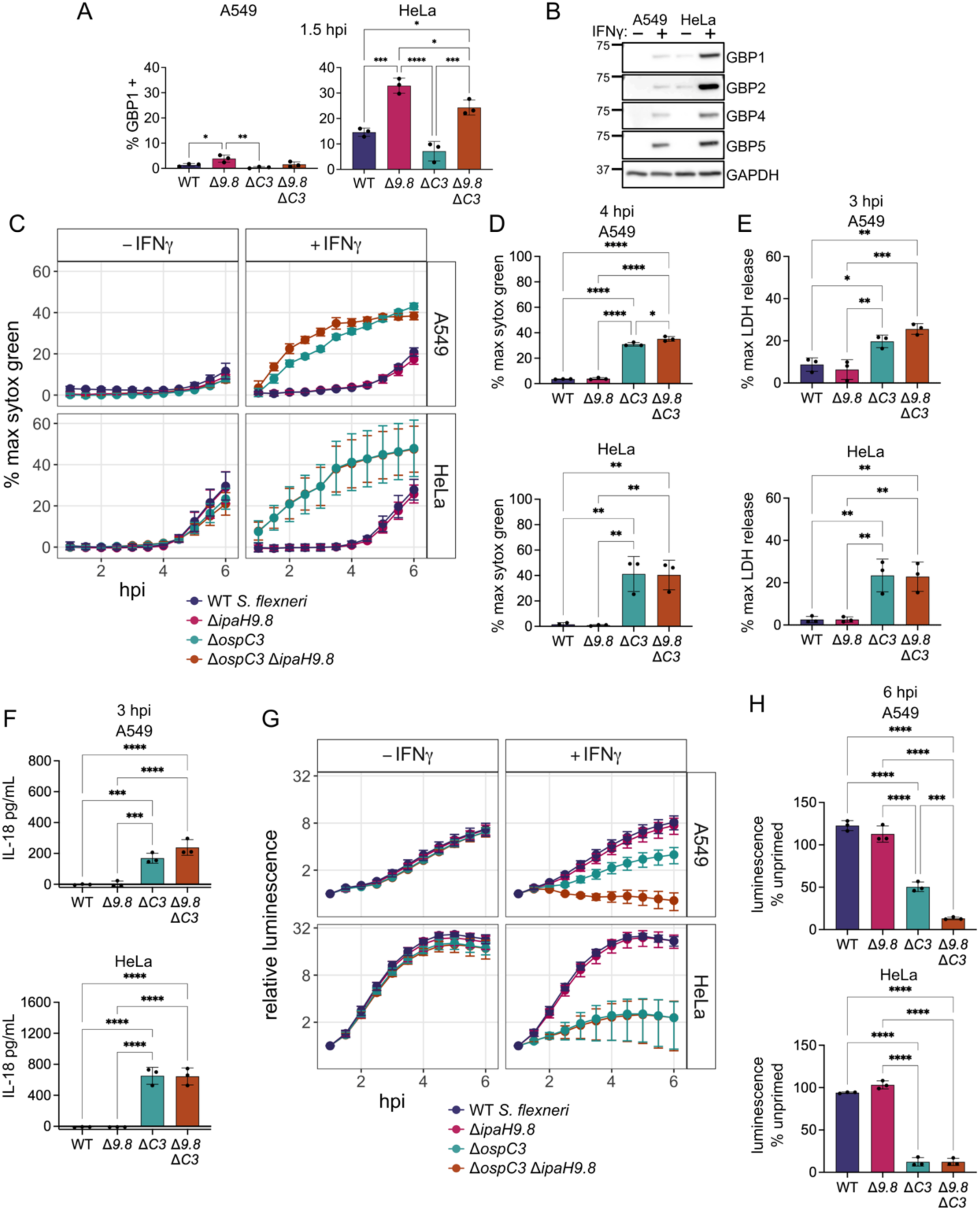
GBP1 targeting of *S. flexneri* is reduced in A549 cells compared to HeLa cells, but pyroptosis levels are similar. (A) A549 and HeLa cells were infected with the indicated strains, fixed at 1.5 hours post infection and immunostained for GBP1. *S. flexneri* with GBP1 surrounding >50% of the bacterial surface were counted as GBP1 positive. (B) A549 and HeLa cells were grown with or without 100 U/ml IFNγ overnight, then lysed and used for western blot. Membrane was probed with antibodies against the indicated GBPs and GAPDH. (C-H) A549 and HeLa cells unprimed or primed with 100 U/ml IFNγ overnight were infected with the indicated *S. flexneri* strains expressing a bioluminescent reporter plasmid. Cell death was measured over time using sytox green fluorescence (C). 4 hour timepoint of IFNγ primed cells was used for statistical analysis (D). Supernatant from IFNγ primed cells infected with *S. flexneri* was removed at 3 hpi and LDH levels (E) or IL-18 secretion (F) were measured. Bacterial luminescence was measured on a plate reader over time (G). Luminescence measurements from the 6 hour timepoint were used to calculate the growth of each strain in primed cells relative to unprimed cells (H). Graphs are averages from three independent experiments and are represented by mean ± SD. One-way ANOVA with Tukey’s multiple comparisons test was used, all statistically significant comparisons are shown. * = P < 0.05, ** = P < 0.01, *** = P < 0.001, **** = P < 0.0001.

If GBP binding to the bacterial surface is required for cell death, we hypothesized that A549 cells would exhibit lower levels of pyroptosis during *S. flexneri* infection, and that bacterial killing would be similarly reduced. We also hypothesized that *S. flexneri* Δ*ospC3*Δ*ipaH9.8* would induce higher levels of cell death than *S. flexneri* Δ*ospC3*. To study pyroptosis and bacterial killing concurrently over time, we infected cells with *S. flexneri* containing a bioluminescent reporter plasmid to monitor bacterial growth. Pyroptosis was measured using the DNA dye sytox green, which fluoresces in cells with compromised plasma membrane integrity, such as pyroptotic cells with GSDMD pores (32). When we infected A549 and HeLa cells there was significant IFNγ-dependent pyroptosis triggered by both *S. flexneri* Δ*ospC3* and *S. flexneri* Δ*ospC3*Δ*ipaH9.8*, while there was no detectable cell death for wildtype or *S. flexneri* Δ*ipaH9.8* (Fig 1C-D). We saw similar results using lactate dehydrogenase (LDH) release as a measure of the cellular rupture that occurs during pyroptosis (Fig. 1E). Interestingly, in IFNγ primed cells there was a similar amount of cell death in both cell lines, despite HeLa cells having significantly more GBP1 targeting of *S. flexneri*. There was also very little difference in the amount of cell death induced by infection with *S. flexneri* Δ*ospC3* or *S. flexneri* Δ*ospC3*Δ*ipaH9.8*, suggesting that IpaH9.8 plays a limited role in reducing cell death through GBP degradation. As an independent measure of inflammasome activation, we measured IL-18 secretion. IFNγ-dependent IL-18 processing and release only occurred after infection with *S. flexneri* Δ*ospC3* or *S. flexneri* Δ*ospC3*Δ*ipaH9.8*, and there was no significant difference in the amount of IL-18 secretion induced by these two strains (Fig. 1F). For bacterial restriction, there was no difference in growth of the different strains in unprimed cells, however IFNγ priming elicited significant growth restriction of both *S. flexneri* Δ*ospC3* and *S. flexneri* Δ*ospC3*Δ*ipaH9.8* (Fig. 1G, H). While in HeLa cells there was no significant difference in restriction between *S. flexneri* Δ*ospC3* and *S. flexneri* Δ*ospC3*Δ*ipaH9.8*, in A549 cells *S. flexneri* Δ*ospC3* was restricted significantly less than *S. flexneri* Δ*ospC3*Δ*ipaH9.8* (Fig. 1H). The differences between HeLa and A549 cells regarding the effect of IpaH9.8 on bacterial burden are not readily explained and may reflect cell line-specific variation in any activity modulated by IpaH9.8, which not only targets GBPs but also NEMO, a protein that regulates NF-κB-dependent signaling (33). Because *S. flexneri* Δ*ospC3* and *S. flexneri* Δ*ospC3*Δ*ipaH9.8* displayed comparable phenotypes across all assays in both cell lines, we focused our studies on the comparison of wildtype and *S. flexneri* Δ*ospC3*Δ*ipaH9.8* from here on forward.

### Endogenous GBP1 is the only GBP required for cell death or bacterial killing

Human GBP1, GBP2, GBP3, and GBP4 have been suggested to form a complex on the bacterial membrane that is essential for pyroptosis and bacterial killing. However, in epithelial cells only knockout cells of GBP1 have been tested, whereas other GBPs were targeted for reduced expression using siRNAs (25, 26). To avoid possible off-target effects prevalent with siRNA technology and to test the role of each GBP independently, we generated CRISPR knockout lines of GBP1, 2, 3, 4, or 5 in A549 and HeLa cells (Fig. S1A-C). In the A549 knockout cells, only knockout of GBP1 led to a significant reduction in cell death. GBP2, 3, 4, and 5^KO^ cells had similar levels of cell death to wildtype cells in response to infection with *S. flexneri* Δ*ospC3*Δ*ipaH9.8* (Fig. 2A). IL-18 secretion was completely lost in GBP1^KO^ cells, while secretion from GBP2-5^KO^ knockout cells was comparable to levels observed with wildtype cells (Fig. 2B). For bacterial restriction there were similar results; *S. flexneri* Δ*ospC3*Δ*ipaH9.8* growth was only restored in the GBP1^KO^ cells, while there was still IFNγ-dependent growth restriction in GBP2-5^KO^ cells (Fig. 2C, D). In HeLa cells we saw identical results to A549 cells, where only GBP1 was required for pyroptosis, IL-18 secretion, or restriction of bacterial growth (Fig. 2E-H). Since we observed very similar results between HeLa and A549 cells for all assays, and GBP1 is the only GBP required in both A549 and HeLa cells, we focused on a single cell line, A549 GBP1^KO^ cells, from here on to investigate why GBP1, but not the other GBPs, is required for pyroptosis during *S. flexneri* infection.

**Figure 2.**
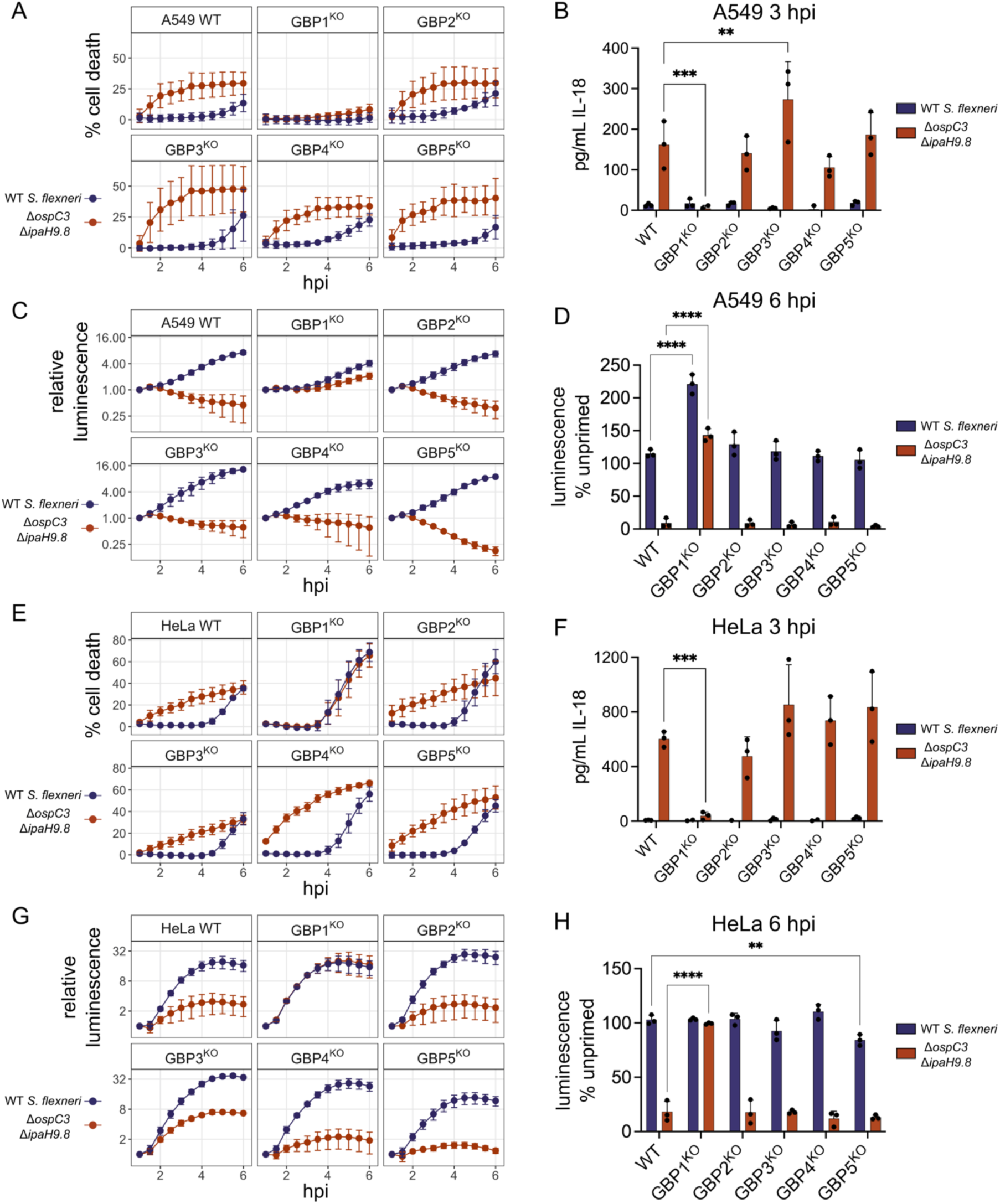
Both A549 and HeLa cells require endogenous GBP1 for pyroptosis and restriction of *S. flexneri* growth. A549 cells (A-D) or HeLa cells (E-H) were primed overnight with 100 U/ml IFNγ, then infected with wildtype *S. flexneri* or *S. flexneri* Δ*ipaH9.8*Δ*ospC3* expressing a bioluminescent reporter plasmid. Cell death was measured using sytox green (A and E). Supernatant was taken at 3 hpi to measure IL-18 secretion (B and F). Bacterial growth was monitored using a luminescence over time (C and G). Luminescence measurements from the 6 hour timepoint were used to calculate the growth of each strain in primed cells relative to unprimed cells (D and H). Data are averages from three independent experiments and are represented by mean ± SD. Two-way ANOVA with Dunnett’s multiple comparisons test was used, for each bacterial strain values for each knockout cell line were compared to the wildtype cells. All statistically significant comparisons are shown. * = P < 0.05, ** = P < 0.01, *** = P < 0.001, **** = P < 0.0001.

### Overexpression of GBP2 promotes pyroptosis in the absence of GBP1

Although endogenous GBP2, 3, 4, and 5 are not essential for pyroptosis or bacterial restriction, they share many structural similarities with GBP1 and could have some functional redundancy with GBP1. Individual GBP paralogs differ from each other in their respective expression levels and it is therefore unclear whether GBP1 is the only GBP capable of promoting pyroptosis, or alternatively whether endogenous GBP1 is the only GBP expressed at high enough levels to promote pyroptosis in the cells we tested. We hypothesized that overexpression of each GBP individually would reveal which of the paralogs are able to promote pyroptosis or restrict *S. flexneri* growth. We overexpressed GBP1, 2, 3, 4, and 5 in GBP1^KO^ cells and measured bacterial binding, cell death, and bacterial restriction during *S. flexneri* infection. We first confirmed previous studies (24–27, 29) showing overexpressed GBP2, 3, 4, and 5 are not recruited to bacteria in the absence of GBP1 (Fig. S2A-C). In unprimed cells there was minimal cell death with any construct, although GBP1 overexpression did promote some cell death (Fig. 3A and 3B). In IFNγ primed cells, overexpression of either GBP1 or GBP2 rescued cell death in GBP1^KO^ cells, while overexpression of GBP3, 4, and 5 did not (Fig. 3C and 3D). For bacterial growth, there was minimal restriction in unprimed cells, with GBP1 overexpression reducing the *S. flexneri* Δ*ospC3*Δ*ipaH9.8* bioluminescence signal by approximately half (Fig. 3E). In IFNγ primed cells both GBP1 and GBP2 rescued *S. flexneri* restriction in GBP1^KO^ cells (Fig. 3F and 3G). It is unclear whether the requirement for IFNγ is due to the upregulation of GBP3, GBP4, and GBP5, or if there are other ISGs that are required for efficient execution of pyroptosis. In primed cells there was no significant difference between the efficiency of GBP1 or GBP2 at executing pyroptosis or restricting bacterial growth. This shows that GBP1 is not unique, and GBP2 can also promote pyroptosis and bacterial restriction.

**Figure 3.**
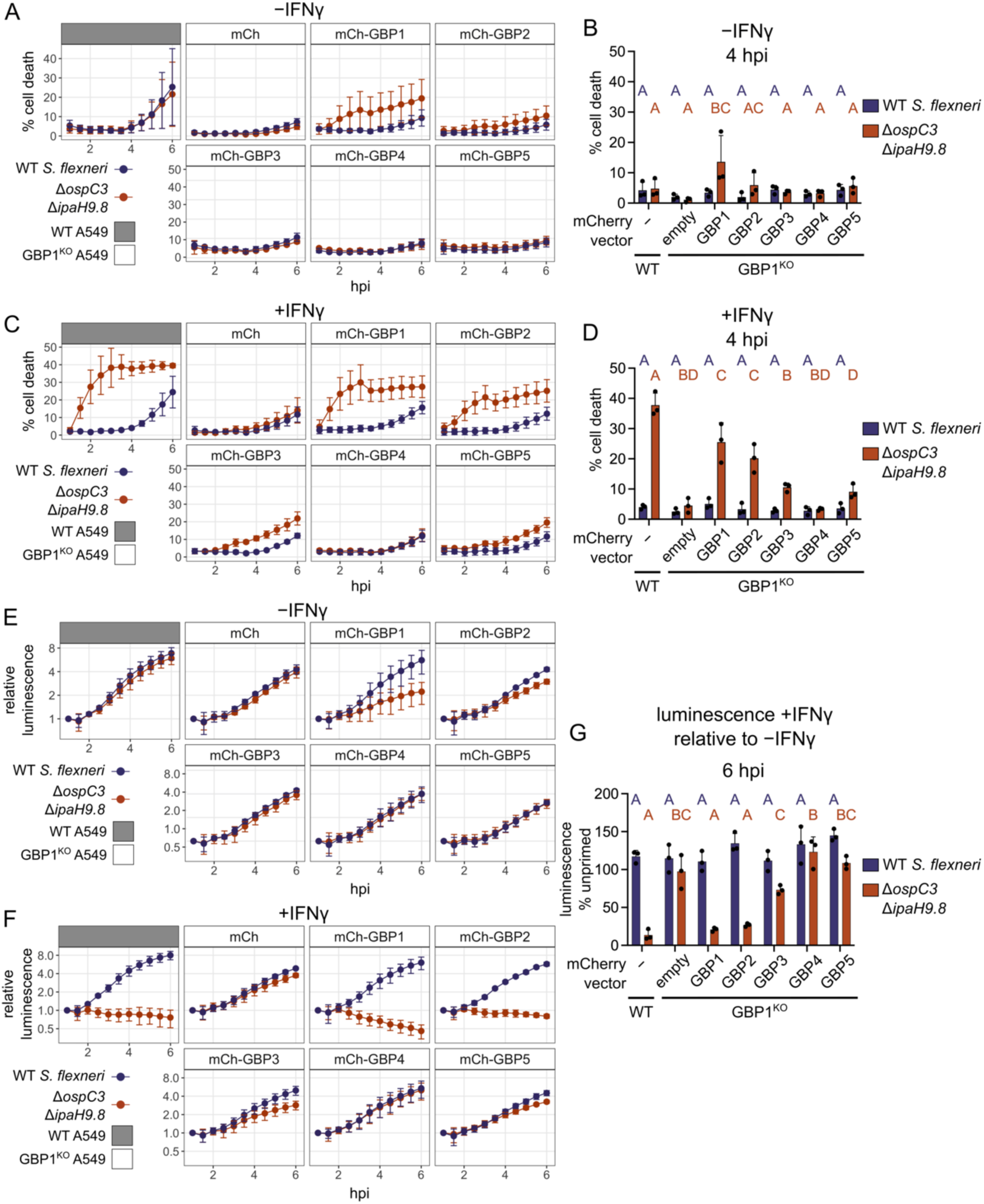
Overexpression of GBP1 or GBP2 can rescue pyroptosis and bacterial restriction in GBP1^KO^ cells. A549 cells were stably transduced to express mCherry or mCherry-GBPs. Cells were then infected with bioluminescent *S. flexneri* and cell death was measured over time using sytox green fluorescence in unprimed cells (A) or cells primed with 100 U/ml IFNγ overnight before infection (C). The sytox green signal at 4 hours was used to determine statistical significance (B and D). Bacterial luminescence was measured over time (E and F). Luminescence measurements from the 6 hour timepoint were used to calculate the growth of each strain in primed cells relative to unprimed cells (G). Data are averages from three independent experiments and are represented by mean ± SD. Two-way ANOVA with Tukey’s multiple comparisons test was used. Statistical comparisons are shown by letters, with bars sharing no matching letters being significantly different. Purple letters correspond to statistical comparisons for wildtype *S. flexneri*, and orange letters correspond to *S. flexneri* Δ*ipaH9.8*Δ*ospC3*.

### Binding of GBP2 to the bacterial surface requires the formation of mixed polymers with GBP1

Our finding that GBP2 overexpression can restore host cell pyroptosis in response to *S. flexneri* invasion of GBP1-deficient epithelial cells led us to hypothesize that GBP2 shares specific biochemical characteristics with GBP1 that are required for pyroptosis induction. Previous work showed that following GTP hydrolysis, GBP1 exits its closed monomeric state, adopts a dimeric outstretched state, and forms dimers (Fig. S3A) (34). GBP1 dimers can assemble into large polymers holding over 1000 molecules (35, 36). These GBP1 polymers attach to the bacterial surface and transition into a GBP1 microcapsule encasing the entire bacteria (24). Because GBP1 polymerization and microcapsule formation are dependent on the post-translational attachment of farnesyl lipid moieties to GBP1 molecules, we asked whether the other two lipidated members of the GBP family, geranylgeranylated GBP2 and GBP5 (37) (Fig 4A), could similarly form polymers and attach directly to bacteria. Although GBP2 and other GBP paralogs are unable to dock on cytosolic bacteria in GBP1-deficient epithelial cells (Fig. S2A-C), we hypothesized that lipidated GBP2 or GBP5 may bind bacteria transiently or at low levels that are difficult to detect in cells but could be detectable in a highly manipulable and more sensitive *in vitro* GBP-bacteria binding assay. We recently developed such an *in vitro* binding assay for GBP1 (24) and here leveraged an analogous system to test the ability of GBP2 and GBP5 to bind to bacteria. For these experiments, we added recombinant Alexa Fluor 488 (488)-labeled GBP1, GBP2, or GBP5 supplemented with GTP to broth cultured RFP-expressing *S. flexneri* and observed that only GBP1 but not GBP2 or GBP5 formed a microcapsule surrounding *S. flexneri in vitro* (Fig. 4B, Movie 1). These data further supported the concept that GBP1 but not GBP2 or GBP5 can directly and autonomously bind bacteria.

**Figure 4.**
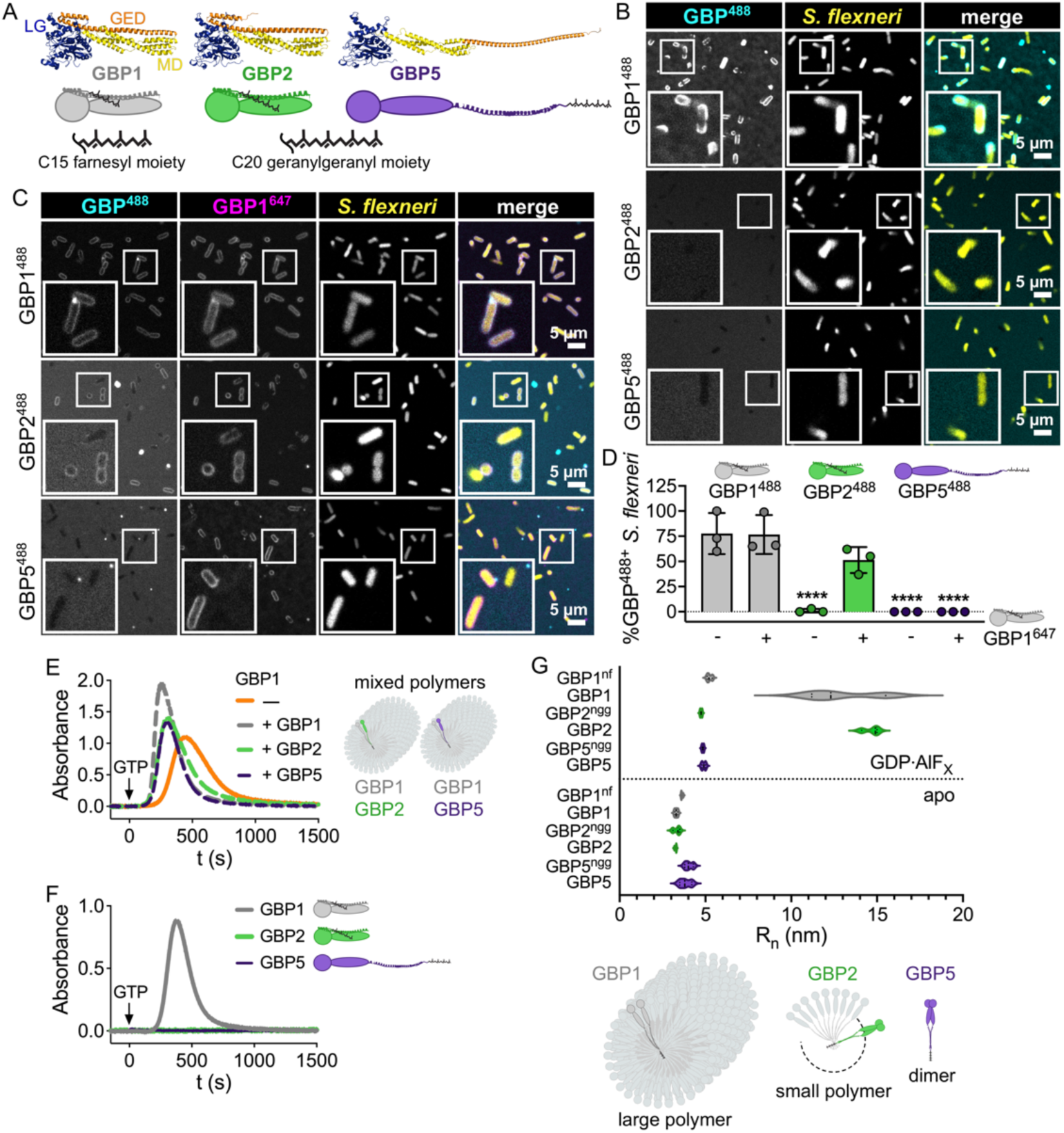
GBP2 fails to bind bacteria on its own but forms small polymers. (A) Structures and schematics for GBP1, GBP2, and GBP5. All GBP paralogs consist of a large GTPase domain (LG), a middle domain (MD), and a GTPase effector domain (GED), whereby C-terminal isoprenyl moieties vary with GBP1 becoming post transcriptionally farnesylated, and GBP2 and GBP5 geranylgeranylated. (B-D) Recombinant Alexa Fluor 488-labeled GBP1, GBP2, and GBP5 alone (B) or mixed with Alexa Fluor 647-labeled GBP1 (C) were supplemented with GTP and added to formaldehyde-fixed RFP-expressing *S. flexneri*. Confocal microscopy time-lapses were recorded, and targeted bacteria were quantified after 1 h (D). (E, F) GBP polymerization was monitored in UV-absorbance-based light scattering experiments. Equal molar ratios of recombinant GBP1, GBP2, or GBP5 were mixed with GBP1, and polymerization was induced with GTP (E). GTP-induced polymerization of single GBPs was monitored over time (F). (G) Number-weighted mean radius (R_n_) of nucleotide-free (apo) or GDP·AlF_X_-bound nonisoprenylated (non-farnesylated - nf, non-geranylgeranylated - ngg) and isoprenylated GBPs were determined in DLS experiments. (A) PDB entry 1F5N and AlphaFold models AF-P32456, AF-Q96PP8. (B, C) Representative images of three independent experiments. (D, G) Graphs are averages from three independent experiments and are represented by mean ± SD. (D) One-way ANOVA with Dunnett’s multiple comparisons test comparing to GBP1^488+^ *S. flexneri* was used, all statistically significant comparisons are shown. **** = P < 0.0001. (E, F) Representative graphs from two independent experiments.

We next investigated the mechanism by which GBP1 facilitates GBP2 binding to cytosolic bacteria in infected cells, which was not previously determined (25–28). We found that upon addition of GTP, Alexa Fluor 647 (647) labeled GBP1 formed mixed polymers with 488-labeled GBP1, GBP2, and GBP5, which associated with the bacterial surface (Fig. S3B). Over time, the surface-attached GBP1-GBP1 and GBP1-GBP2 polymers formed a microcapsule surrounding the bacterial cell, whereas GBP5 failed to be incorporated in the GBP1 microcapsule (Fig. 4C-D, Movie 2). Incorporation of GBP2 and GBP5 into GBP1 polymers was confirmed in UV-absorption-based light scattering experiments, where we observed that the addition of equimolar amounts of GBP1, GBP2, or GBP5 to GBP1 accelerated the initiation of polymerization and increased the absorbance signal (Fig. 4E). Together these experiments showed that although GBP1 recruits both GBP2 and GBP5 to the bacterial surface as mixed polymers, only the GBP1-GBP2 mixed polymer transforms into a microcapsule. This explains the reported observation that GBP5 is not part of the GBP coat surrounding cytosolic bacteria in infected cells (25–28). Our present observations further implied that GBP2 only interacts with the bacterial outer membrane when stabilized in its active conformation by GBP1 through mixed polymer formation. Because GBP2-driven pyroptosis did not require GBP1 (Fig. 3), we can conclude that mixed GBP1-GBP2 polymer formation and the resulting GBP1-dependent binding of GBP2 to the bacterial surface are dispensable for GBP2-dependent cell death.

### GBP2 self-assembles into small polymers independent of GBP1

We previously reported that reversible GBP1 polymerization precedes encapsulation of Gram-negative bacteria. We also showed that GBP1 polymerization promotes clustering of LPS (24). To determine whether, similar to farnesylated GBP1, geranylgeranylated GBP2 and GBP5 self-assemble into polymers, we monitored GTP-induced polymerization of GBPs using the UV-absorption based turbidity assay in which light scattering by large protein polymers leads to an increase in absorbance (35, 36). We observed that only GBP1 but not GBP2 or GBP5 formed large polymers during GTP hydrolysis which dissociated with GTP depletion (Fig. 4F). To stabilize smaller, short-lived GBP complexes, we utilized the GTP transition state analog GDP·AlF_X_ in dynamic light scattering (DLS) experiments. We determined the number-weighted mean radius (R_n_) as a measure of size for nonisoprenylated and isoprenylated GBPs in their nucleotide-free, inactive (apo), and GDP·AlF_X_-bound, activated states (Fig. 4G, S2F). As expected, all GBPs were monomeric in their nucleotide-free resting state with nonisoprenylated and isoprenylated GBP5 appearing slightly larger in size due to their outstretched conformations (Fig. 4A). In the presence of GDP·AlF_X_ the R_n_ of nonisoprenylated GBP1, GBP2, and GBP5 increased to approximately 5 nm indicating that dimers formed by different GBP paralogs have a uniform size (Figs. 4G and S2F). While GDP·AlF_X_-bound geranylgeranylated GBP5 remained dimeric, farnesylated GBP1 and geranylgeranylated GBP2 self-assembled further into complexes with R_n_ of approximately 13-15 nm, corresponding to small polymers. Together these light scattering experiments demonstrate that, although GBP1 is unique in forming large polymers, GBP2, like GBP1, can self-assemble into small polymers. Since GBP2 can form polymers on its own, but not bind bacteria, this suggested that polymerization rather than bacterial binding is an important characteristic that allows GBP2 to promote pyroptosis during infection.

### GBP1 binding to bacterial surface is dispensable for cell death and bacterial killing

Because GBP1 binding to the bacterial surface precedes cell death during *Salmonella enterica* Typhimurium or *S. flexneri* infection, it has been suggested that GBP1 binding to bacteria is required for caspase-4 activation (25, 26). We showed that GBP2 promotes pyroptosis without binding to bacteria and therefore hypothesized that a GBP1 mutant that can bind free LPS and polymerize, but not bind the bacterial outer membrane would also be able to promote pyroptosis. To test this hypothesis, we monitored pyroptosis or bacterial killing of infected cells that overexpress wildtype and mutant GBP1 variants in GBP1^KO^ cells. As expected, wildtype GBP1 completely restored pyroptosis and bacterial killing in GBP1^KO^ cells, whereas there was no rescue with the empty mCherry vector (Fig. 5A,B). We previously showed that a polybasic motif of three arginines (3R) at the C-terminus of GBP1 is important for stable binding of GBP1 to the bacterial surface (24, 27, 29) Importantly, like wildtype GBP1, GBP1^3R^ is still able to polymerize, as well as bind and cluster free LPS (24). In GBP1^KO^ cells, overexpression of GBP1^3R^ restored cell death and bacterial killing to similar levels as wildtype GBP1 (Fig. 5A-D). This shows that GBP1 binding to the *S. flexneri* surface is dispensable for both pyroptosis and restriction of bacterial growth. GBP1^R48A^, which lacks GTPase activity and cannot polymerize, was unable to promote cell death or bacterial killing (Fig. 5A-D). Similarly, GBP1^C589A^, which lacks the farnesyl tail and is unable to form polymers, cluster free LPS, or bind the bacterial surface, did not restore cell death or bacterial restriction.

**Figure 5.**
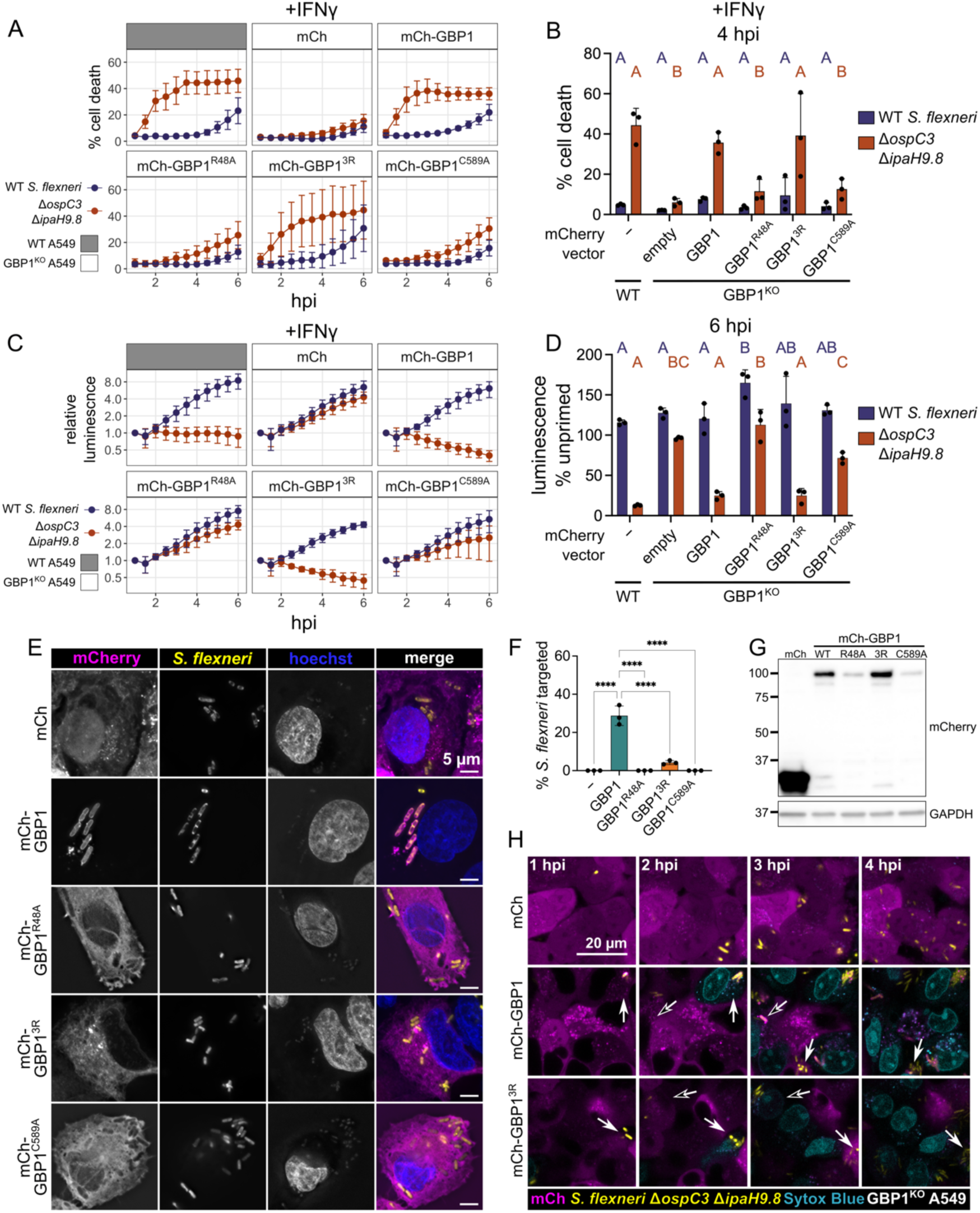
GBP1-dependent pyroptosis and restriction of *S. flexneri* growth is independent of GBP1 binding to bacteria. (A-D) Wildtype A549 or GBP1^KO^ cells expressing mCherry or mCherry-GBP1 mutants were unprimed or primed with 100 U/ml IFNγ overnight, where indicated. Cells were infected with *S. flexneri* expressing a bioluminescent reporter and cell death was measured over time using sytox green fluorescence (A). The sytox green signal at 4 hours was used to determine statistical significance (B). Bacterial luminescence was measured over time (C). Luminescence measurements from the 6 hour timepoint were used to calculate the growth of each strain in primed cells relative to unprimed cells (D). (E and F) Wildtype A549 or GBP1^KO^ cells expressing mCherry or mCherry-GBP1 mutants were primed overnight with IFNγ, then infected with GFP expressing *S. flexneri* Δ*ipaH9.8* and fixed at 2 hours post infection. (E) Coverslips were imaged at 100x magnification using widefield microscopy. Images were deconvolved and z-projection is shown, scale bar is 5 μm. (F) Coverslips were imaged at 63x magnification, with images taken from five independent fields. Targeting of *S. flexneri* by each overexpressed protein was quantified using ImageJ. *S. flexneri* with indicated protein around at least 50% of the bacterial membrane were counted as targeted. (G) Expression levels of indicated overexpressed proteins determined by western blot. (H) Frames from timelapse microscopy at indicated time points for GBP1^KO^ cells expressing mCherry, mCherry-GBP1, or mCherry-GBP1^3R^ infected with GFP expressing *S. flexneri* Δ*ospC3*Δ*ipaH9.8*. Dying cells are shown in blue (sytox blue). All graphs show averages from three independent experiments and are represented by mean ± SD. (B and D) Significance determined using two-way ANOVA with Tukey’s multiple comparisons test. Statistical comparisons are shown by letters, with bars sharing no matching letters being significantly different. Purple letters correspond to statistical comparisons for wildtype *S. flexneri*, and orange letters correspond to *S. flexneri* Δ*ipaH9.8*Δ*ospC3*. (F) One-way ANOVA with Tukey’s multiple comparisons test was used. All significant comparisons are shown. * = P < 0.05, ** = P < 0.01, *** = P < 0.001, **** = P < 0.0001.

We next confirmed that GBP1, but not GBP1^3R^, binds to the surface of *S. flexneri* Δ*ipaH9.8* (Fig. 5E, 5F, 5G) within infected cells. To check whether there was transient binding of GBP1^3R^ immediately preceding cell death we used timelapse microscopy to view cells expressing mCherry-GBP1 or GBP1^3R^ and infected with GFP expressing *S. flexneri* Δ*ospC3*Δ*ipaH9.8*. Using the cell-impermeant dye sytox blue to label cells undergoing pyroptosis, we confirmed prior studies showing GBP1 targeting of *S. flexneri* Δ*ospC3*Δ*ipaH9.8* precedes cell death (Fig. 5H, Movie 3). Importantly, in cells expressing mCherry-GBP1^3R^ we observed pyroptosis in cells without appreciable binding of GBP1^3R^ to the bacterial surface (Fig. 5H, Movie 3). This supports a new model where GBP1 or GBP2 can promote caspase-4 activation without binding the bacterial surface. Demonstrating that this result was not specific to *S. flexneri*, we found that GBP1, GBP1^3R^ and GBP2 were also able to rescue pyroptosis in GBP1^KO^ cells during *S.* Typhimurium infection (Fig. S3A and S3B). Because GBP2 and GBP1^3R^ cannot bind *S.* Typhimurium in the absence of wildtype GBP1 (29), we conclude that GBP binding to the surface of *S. flexneri* or *S.* Typhimurium is dispensable for GBP-dependent pyroptosis.

### GBP1, GBP1^3R^, or GBP2 can mediate cell death in response to LPS transfection

Since we observed rescue of pyroptosis by both GBP1^3R^ and GBP2 during *S. flexneri* and *S.* Typhimurium infection, we wondered whether these proteins could mediate pyroptosis in response to direct delivery of LPS to the cytosol. First, we tested two methods of LPS delivery to the cytosol: electroporation and transfection. For electroporation, although there was lower cell death in GBP1^KO^ cells at early timepoints with lower concentrations of LPS, GBP1 was largely dispensable for pyroptosis (Fig. S4A and S4B). This suggests that electroporation is not a good model for GBP1-dependent pyroptosis. Conversely, there was a significant reduction in pyroptosis in response to LPS delivered through transfection (Fig. 6A). To test if GBP1^3R^ or GBP2 could promote pyroptosis in response to cytosolic free LPS, we overexpressed GBP1, GBP1^3R^, or GBP2 in GBP1^KO^ cells and measured cell death following LPS transfection. With two concentrations of LPS, expression of GBP1, GBP1^3R^, or GBP2 promoted similar levels of cell death in GBP1^KO^ cells (Fig. 6B). There was also no significant difference in IL-18 secretion between cells expressing GBP1, GBP1^3R^ or GBP2 (Fig. 6C).

**Figure 6.**
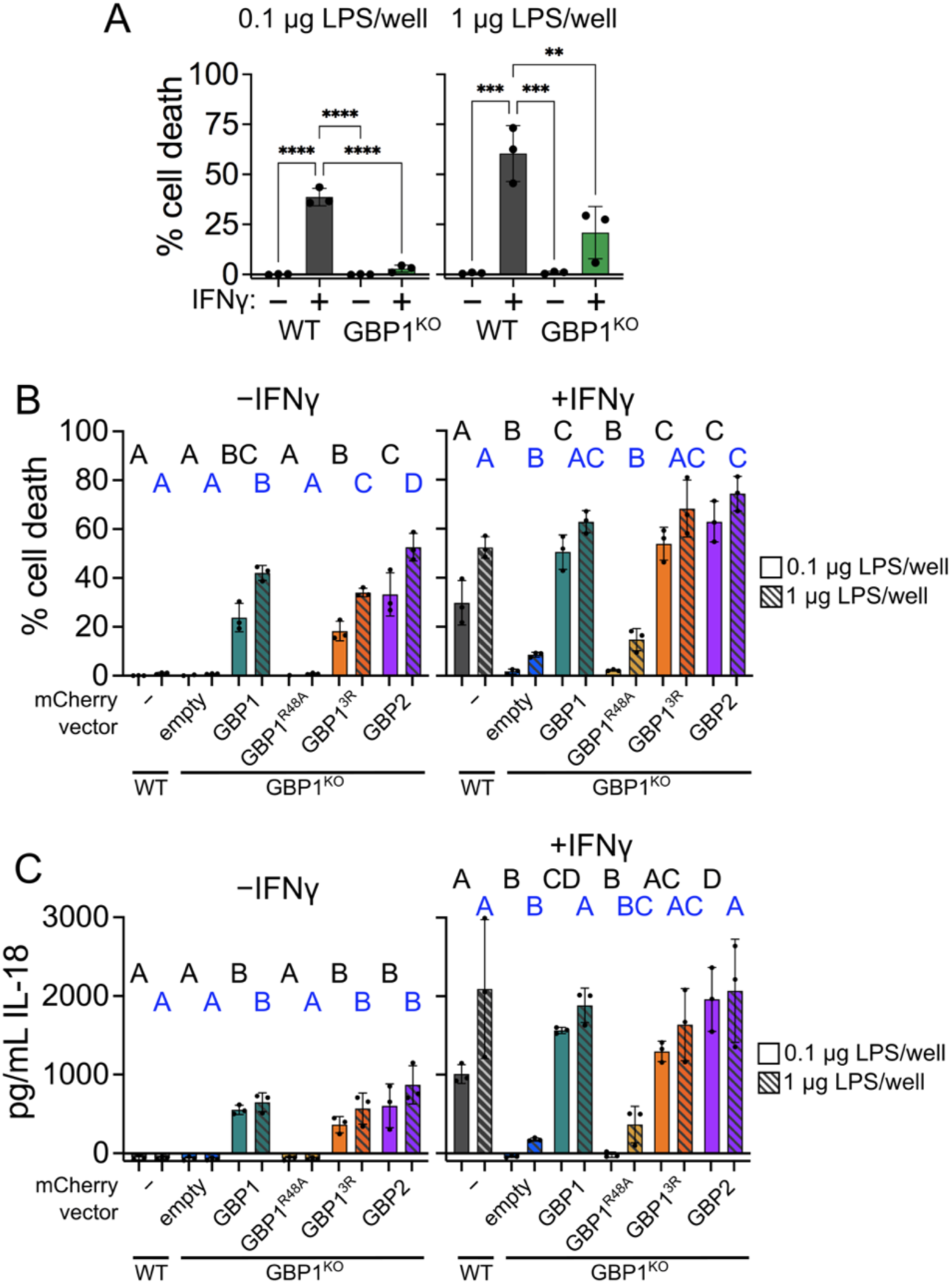
Ectopically expressed GBP1, GBP1^3R^, and GBP2 can rescue pyroptosis in GBP1^KO^ cells transfected with LPS. (A) Wildtype and GBP1^KO^ A549 cells in 96 well plates were unprimed or primed with 100 U/ml IFNγ overnight, then transfected with 0.1 μg or 1 μg per well *E. coli* O55:B5 LPS. Cell death was measured using sytox green fluorescence at 6 hours post transfection. (B and C) Wildtype A549 or GBP1^KO^ A549 cells overexpressing mCherry or the indicated mCherry-GBPs in 96 well plates were unprimed or primed with 100 U/ml IFNγ overnight, then transfected with 0.1 μg (solid bars) or 1 μg (striped bars) *E. coli* O55:B5 LPS per well. Cell death was measured using sytox green fluorescence at 6 hours post transfection (B). IL-18 secretion was measured in supernatants taken at 6 hours post transfection (C). All graphs show averages from three independent experiments and are represented by mean ± SD. (A) One-way ANOVA with Tukey’s multiple comparisons test was used. All significant comparisons are shown. * = P < 0.05, ** = P < 0.01, *** = P < 0.001, **** = P < 0.0001. (B and C) Significance determined using two-way ANOVA with Tukey’s multiple comparisons test. Statistical comparisons are shown by letters, with bars sharing no matching letters being significantly different. Black letters correspond to statistical comparisons for 0.1 μg per well LPS, and blue letters correspond to 1 μg per well LPS.

Interestingly, even in the absence of IFNγ priming there was significant cell death and IL-18 secretion in cells overexpressing GBP1, GBP1^3R^ or GBP2, indicating that expression of additional GBPs or other interferon stimulated genes is not required for cell death in response to cytosolic LPS (Fig. 6B and 6C). It was previously suggested that even with concurrent GBP1, 3, and 4 overexpression in unprimed cells there was significantly lower levels of cell death following LPS transfection compared to IFNγ primed cells (25). Our results suggest that overexpression of GBP1 or GBP2 alone is sufficient to promote pyroptosis, although addition of IFNγ does enhance the response.

### GBP2 acts as a surfactant and aggregates LPS

Because GBP2 induces pyroptosis in response to ‘free’ LPS in the cytosol, we hypothesized that it can directly recognize LPS similar to GBP1. We reported previously that GBP1 is an LPS-binding surfactant which clusters soluble LPS into larger LPS aggregates independent of its 3R motif (24). To test whether GBP2, like GBP1, can induce LPS clustering we mixed fluorescently labeled LPS with recombinant GBPs and GTP and assessed the LPS particle number and size with fluorescence microscopy (Fig. 7A, Fig S6A). We found that GBP2, like GBP1 and GBP1^3R^, increased the LPS aggregate area during GTP hydrolysis, although fewer aggregates were formed by GBP2. Nonisoprenylated GBPs and geranylgeranylated GBP5 on the other hand failed to induce clustering of LPS to larger aggregates. We further observed that GDP·AlF_X_-induced GBP1 and GBP2 polymers shifted to larger and smaller R_n_ when supplemented with LPS (Fig. S6B) which suggest changes in polymer size and/or polymer conformation due to LPS incorporation. However, these DLS experiments were inconclusive, because sizes of LPS aggregates comprised a broad range and could not be definitively distinguished from GBP monomers and complexes. We therefore employed native polyacrylamide gel electrophoresis (NPAGE) to separate GBP, LPS, and GBP-LPS aggregates not only by size but also by charge, then stained successively for LPS and protein to detect changes in LPS and GBP mobility. To stabilize either the resting state or the active state of the respective GBP in these experiments, we supplemented protein-LPS mixtures with GDP or GDP·AlF_X_. The migration of bovine serum albumin (BSA, negative control) and GBP5 was unchanged in the presence of LPS. However, GBP1 and GBP2 showed shifts in mobility with increasing LPS concentrations (Fig. 7C). This confirms that GBP1 and GBP2 interact with LPS directly and form mixed aggregates. To estimate the affinities of GBP-LPS complexes, we titrated GBP1, GBP1^3R^, and GBP2 with LPS in the presence of GDP·AlF_X_ and analyzed the gray values of the NPAGE gels after protein staining. We then plotted the percentage of protein signal with altered mobility against the LPS concentration (Fig. 7C, S6C). GBP1^3R^ required roughly 10 times and GBP2 roughly 100 times higher LPS concentrations than GBP1 to generate similar shifts in mobility, suggesting that GBP2 binds LPS and forms LPS-protein aggregates with lower affinity than GBP1. Together, these data identify GBP2 as an LPS clustering surfactant, a property that GBP2 shares with GBP1. We propose that their LPS clustering activities account for the role GBP1 and GBP2 play in caspase-4 activation during Gram-negative bacterial infections.

**Figure 7.**
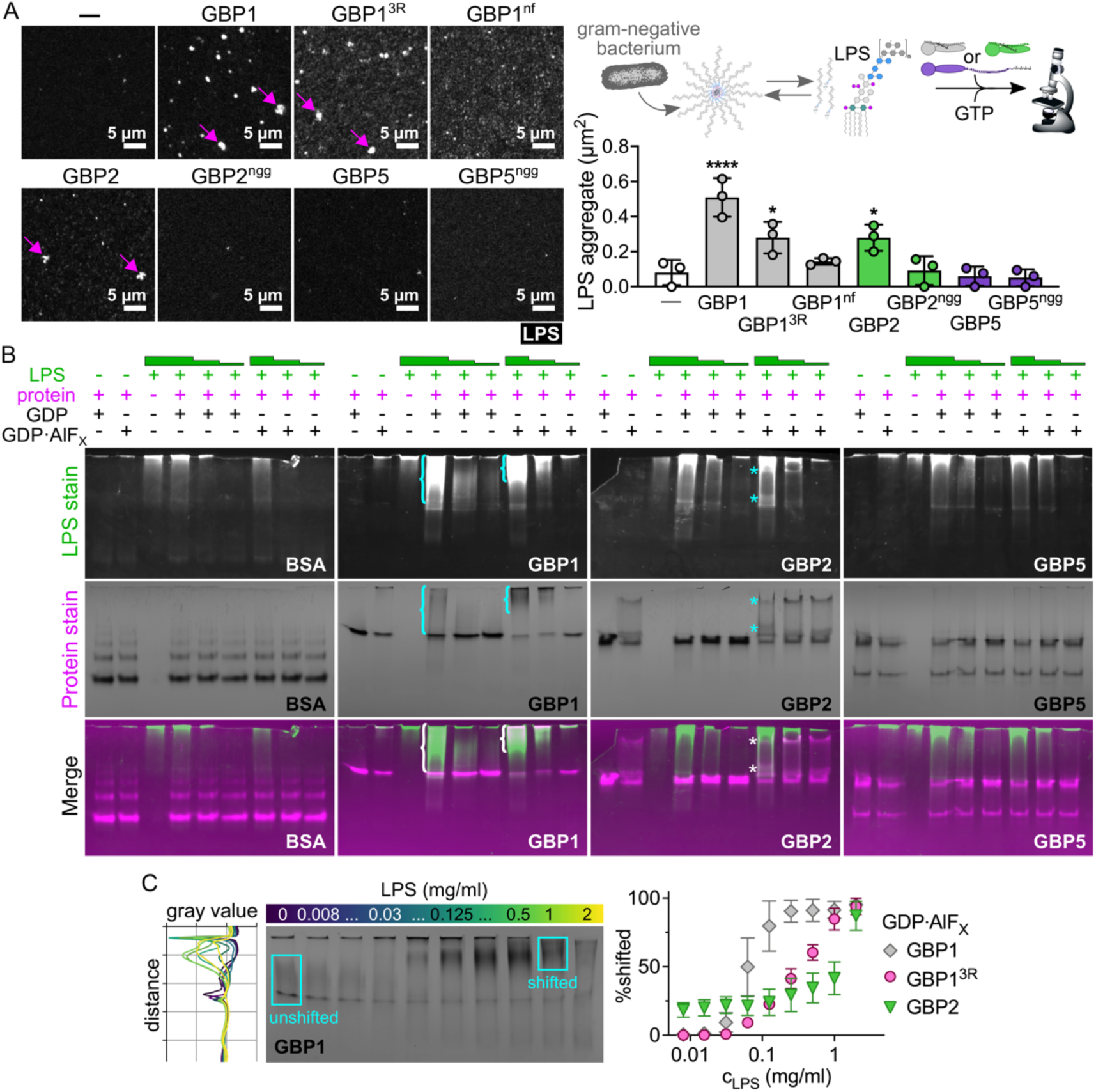
GBP2 binds and aggregates LPS. (A) Recombinant isoprenylated and nonisoprenylated GBPs were supplemented with GTP and added to Alexa Fluor 568-labeled *E. coli* O55:B5 LPS. LPS particles were analyzed from different fields of views taken after 20 min with Fiji, and the average aggregate area was plotted. (B, C) Following mixing of recombinant proteins with *E. coli* O55:B5 LPS in the presence of different nucleotides, protein-LPS complexes were resolved with NPAGE, and gels were stained successively for LPS and proteins. (B) NPAGEs of BSA, GBP1, GBP2, and GBP5 supplemented with GDP or GDP·AlF_X_ in the absence or presence of LPS (final concentrations 1 mg/ml, 0.1 mg/l, or 0.01 mg/ml). (C) NPAGE of GBP1 titrated with LPS (final concentrations 2 mg/ml to 0.008 mg/ml) supplemented with GDP·AlF_X_. Grey values for each gel lane were plotted with Fiji and areas were defined representing unshifted and shifted protein fractions. Percentage of shifted protein was plotted against LPS concentrations. All graphs show averages from three independent experiments and are represented by mean ± SD. (A) One-way ANOVA with Dunnett’s multiple comparisons test comparing to control (no GBP addition) was used. All significant comparisons are shown. * = P < 0.05, **** = P < 0.0001. (B, C) Representative NPAGEs from two (B) or three (C) independent experiments are shown.

## Discussion

The cytokine IFNγ enhances non-canonical inflammasome activation in response to cytosolic LPS through upregulation of the GBP protein family. GBPs promote pyroptosis in different cell types and species, however, the exact role of each human GBP paralog and the mechanism by which GBPs promote pyroptosis has not been clear (38–42). It was previously thought that GBP1 binding to the surface of Gram-negative bacteria, and subsequent recruitment of GBP2, GBP3, and GBP4, were essential steps in initiating LPS-triggered and caspase-4-dependent pyroptosis during infection. While GBP2 can only bind Gram-negative bacteria when GBP1 is present, in this study we showed GBP2 promotes pyroptosis in the absence of GBP1, indicating that bacterial targeting is dispensable for GBP-dependent pyroptosis. We confirmed this by demonstrating that a mutant of GBP1 that binds free LPS but does not form a bacteria-encasing microcapsule, is able to induce pyroptosis during infection. We also found that, like GBP1, GBP2 directly binds and aggregates LPS, and we propose that this LPS clustering by GBPs facilitates caspase-4 activation and pyroptosis.

A major question remaining is why GBP1 and GBP2 have seemingly redundant roles in response to cytosolic LPS. One potential explanation is that GBP1 and GBP2 have unique specificities for different LPS species, allowing detection of a broader range of bacteria. Gram-negative bacteria contain a diverse repertoire of LPS structures, with many variants existing in the O-antigen, core oligosaccharide, and lipid A portions of LPS molecules (43). LPS variants have different abilities to activate sensors TLR4 or caspase-4 (43, 44). Supporting the hypothesis that GBP2 could preferentially drive inflammasome responses to a subset of LPS structures, it was shown that GBP2 was required for efficient activation of caspase-4 in response to tetra-acylated LPS from *Francisella novicida*, while it was dispensable for pyroptosis in response to hexa-acylated *E. coli* LPS (20).

The functions of GBP1 and GBP2 may also be defined by the route of entry of LPS into the cytosol, similar to the differential role for human caspase-4 and caspase-5 in LPS sensing. Caspase-4 and caspase-5 are structurally similar, and both are able to directly bind LPS and induce pyroptosis (4), yet in many assays of LPS-induced pyroptosis, caspase-4 is essential and caspase-5 is dispensable (9, 11, 26). Recently it was shown that caspase-5, but not caspase-4, responds to LPS present in OMVs trafficked through endosomes of intestinal epithelial cells (45). This suggests that the cell may have different ways of responding depending on the delivery route by which LPS reaches the cytosol.

Another reason for GBP1 and GBP2 redundancy may be to allow either cell type-specific or tissue-specific responses to Gram-negative infection, as these GBPs are each expressed at different levels in cell lines and tissues (46, 47). We found that GBP2 has a lower affinity for LPS, so it may be preferentially used to detect cytosolic LPS in tissues where a less robust response is warranted, for example, in the gut where there is abundant LPS released by commensal bacteria. Further studies investigating the differences between GBP1 and GBP2 function may provide an answer for the importance of each protein.

Our study highlights the ability of GBPs to promote immune defense even in the absence of direct binding to pathogens. While many studies have investigated the targeting-dependent functions of GBPs in immunity, there are now several examples of GBPs restricting pathogen growth without translocation to pathogens or pathogen-containing vacuoles. GBPs restrict growth of protozoan parasites *Leishmania donovani* and *Toxoplasma gondii* without detectable binding to the parasitophorous vacuoles (48–50). GBPs also promote pyroptosis in response to avirulent *E. coli* in macrophages, without significant GBP targeting of these bacteria in cells (18). We show here that GBP1 and GBP2 promote pyroptosis without binding the surface of bacteria, likely through aggregation of LPS released during infection. This could be a more universal strategy for GBPs to act as pattern recognition receptors, where pathogen associated molecular patterns (PAMPs), such as LPS but also others, are released into the host cytosol and are detected by GBPs. PAMPs bound by GBPs could then facilitate activation of other defense signaling pathways. Future studies could test this hypothesis and determine which additional PAMPs are detected by GBPs, and if these are similarly being released from pathogen-containing vacuoles and directly bound by GBPs. For pathogens where GBP targeting is observed, it will also be informative for future studies to use GBP mutants to separate targeting-dependent and independent functions.

Our study identifies a new LPS sensor, GBP2, that aggregates LPS and allows caspase-4 activation. While GBPs were known to be important for activation of the non-canonical inflammasome in response to ‘free’ LPS, LPS contained in outer membrane vesicles, or in response to Gram-negative bacterial infection, it was unclear how GBPs function to detect these different forms of LPS. During infection with cytosolic Gram-negative bacteria, one model proposes that GBP binding to the surface of invading bacteria provides a necessary platform for caspase-4 activation. The recent finding that GBP1 directly binds and clusters ‘free’ LPS provides another model for caspase-4 activation, where GBP1 acts as a surfactant to aggregate LPS into larger structures that are a preferred substrate for caspase-4 (24, 51). Since Gram-negative bacteria constitutively shed ‘free’ LPS (52), aggregation of LPS in the cytosol explains both responses to invading bacteria as well as ‘free’ LPS. Our current study supports a model where GBP1 or GBP2 promote caspase-4 activation through aggregation of LPS released into the cytosol during infection.

## Materials and Methods

### Cell lines and cell culture

HeLa and A549 cells (ATCC) were grown in DMEM (Gibco 11995-065) supplemented with 9% heat inactivated FBS (Omega Scientific) and non-essential amino acids (Gibco 11140-050). Cells were grown at 37°C in 5% CO_2_. Cell lines were routinely tested for mycoplasma contamination. Cell lines were authenticated using GenePrint 10 (Promega) performed by the Duke University DNA Analysis Facility.

### Knockout cells

For A549 GBP1^KO^, GBP2^KO^ and GBP4^KO^ cells, single guide RNAs (sgRNAs) sequences to human GBPs were selected using the optimized CRISPR design site <u>crispr.mit.edu</u>. The GBP1 sgRNA sequence was CATTACACAGCCTATGGTGG and the GBP2 sgRNA sequence was CTAGTTCTGCTCGACACTGA. For GBP4 the sgRNA sequences were ATTGTAGGGCTATACCGCACAGG and TATCTCATGAATCGTCTTGCAGG. sgRNAs were cloned into PX459 containing Puromycin resistance ((53); pSpCas9(BB)-2APuro (PX459) was a gift from Feng Zhang (Addgene plasmid #48139) or PX458 containing an eGFP cassette ((53); pSpCas9(BB)-2A-GFP (PX458) was a gift from Feng Zhang (Addgene plasmid #48138) following the Zhang lab, Addgene CRISPR Genome Engineering Toolbox (www.addgene.org/crispr/zhang/). A549 cells were transfected with PX459 or PX458 plasmids containing guide RNAs using FuGENE HD transfection reagent (Promega E2311) following manufacturers guidelines. 48 hours post transfection, PX459 transfected cells were put under Puromycin (Generon 1860-25) selection at 1-2 μg/ml for 48 hours and PX458 transfected cells were sorted one cell per well into a 48 well plate using a FACS Aria Fusion Cell Sorter.

A549 GBP3^KO^, GBP5^KO^, and HeLa GBP2^KO^, GBP3^KO^, GBP4^KO^, and GBP5^KO^ cells were generated by the Duke Functional Genomics core as previously described (54). sgRNAs were designed using CHOPCHOP (55) and Cas-OFFinder (56), then cloned into PX459 V2 (Addgene #62988; (53)). For A549 GBP3^KO^ cells the sgRNA sequences were TCGATCTGCCCATTCACCGC and AGAACTTCCGGATACAGAGT. For A549 GBP4^KO^ cells the sgRNA sequences were ATTGTAGGGCTATACCGCACAGG and TATCTCATGAATCGTCTTGCAGG. For A549 GBP5^KO^ cells the sgRNA sequences were GCTCATTAAAGTTCTCGATG and GCAAAGTAACATCCTAGACA. For HeLa GBP2^KO^ cells the sgRNA sequences were AGAGCTGACAGATCGAATCA and TCGTCTACAGAATTGTTACC. For HeLa GBP3^KO^ cells the sgRNA sequence was CCTCATTGAGAACACTAATG. For HeLa GBP4^KO^ cells the sgRNA sequences were CGTCTTGCAGGAAAGCGCAA and ATTGTAGGGCTATACCGCAC. For HeLa GBP5^KO^ cells the sgRNA sequences were GCTCATTAAAGTTCTCGATG and GCAAAGTAACATCCTAGACA. A549 cells were transfected with sgRNAs using Lipofectamine 3000 (ThermoFisher Scientific) according to manufacturer’s instructions. 24 hours after transfection, cells were selected with 2 mg/mL puromycin (Sigma) for three days. Cells were diluted to isolate single cells, which were expanded to generate clonal cell lines. Knockouts were validated by western blot or sanger sequencing.

To generate CRISPR Cas-9 mediated knockout of *GBP1* in HeLa cells, we chose an sgRNA sequence targeting exon 2 that would minimize off-target editing of the closely related *GBP3* gene, CATTACACAGCCTATGGTGG. The sgRNA was ordered as modified synthetic sgRNA from Synthego and delivered to HeLa cells by electroporating 200,000 cells with 4 ug TrueCut Cas9 protein (ThermoFisher Scientific) complexed with 180 pmol *GBP1* sgRNA using a Neon system (ThermoFisher Scientific) with the following settings: 1005 V, 35 ms, 2 pulses. Following electroporation, cells were recovered in 6 well dishes for three days before harvesting to check KO efficiency. Cells were diluted to isolate single cells, then expanded to generate a clonal knockout line.

### Bacterial strains

*Shigella flexneri* strain 2457T was used. *S. flexneri* Δ*ipaH9.8*, *S. flexneri* Δ*ospC3*, and *S. flexneri* Δ*ipaH9.8*Δ*ospC3* knockouts were made as previously described (24, 27, 57). *S. flexneri* was cultured on tryptic soy broth (TSB, Millipore Sigma) agar plates containing 0.01% Congo Red (Millipore Sigma), then grown in liquid culture in TSB. *Salmonella enterica* Typhimurium strain 14028s was used throughout. *S.* Typhimurium was cultured on lysogeny broth (LB), Miller formulation agar plates, or in liquid LB.

### Infections

For *S. flexneri* infections, *S. flexneri* was streaked on an agar plate, then single colonies were picked and grown overnight at 37°C shaking, in 3 ml TSB. The next day 250 μl overnight culture was diluted into 5 ml TSB and grown for approximately 1 hour 45 minutes, until OD600 was between 0.7 and 1. 1 ml bacterial culture was pelleted and resuspended in 1 ml infection media (phenol red free DMEM (Gibco 31053-028), supplemented with 5% FBS and non-essential amino acids). Bacteria was further diluted in infection media to have enough bacteria for a multiplicity of infection (MOI) of 5 in 100 μl per well of 96 well plate, or 1 ml per well of 24 well plate. Media was aspirated from cells and diluted *S. flexneri* was added to each well. Plate was centrifuged for 10 min, 800×g, at room temperature. The end of the centrifugation was considered the start of infection, and plates were then put in 37°C incubator with 5% CO_2_. After 30 minutes, cells in 96 well plate were washed once with 200 μl warm HBSS. For microscopy, cells in 24 well plate were washed twice with 1ml warm HBSS. After washing, infection medium containing 50 μg/ml gentamicin was added to wells to kill extracellular bacteria. For *S.* Typhimurium infections, single colonies were grown overnight at 37°C shaking, in 1.5 ml LB. Next day 45 μl overnight culture was diluted into 1.5 ml LB and grown for 2 hours 40 minutes. Bacterial culture was pelleted and resuspended in infection media (described above for *S. flexneri* infections). Bacteria was further diluted in infection media to have enough bacteria for a multiplicity of infection (MOI) of 200 in 100 μl per well of 96 well plate. Media was aspirated from cells and diluted *S.* Typhimurium was added to each well. Plate was centrifuged for 10 min, 800×g, at room temperature. The end of the centrifugation was considered the start of infection, and plates were then put in 37°C incubator with 5% CO_2_. After 30 minutes, cells in 96 well plate were washed once with 200 μl warm HBSS. After washing, infection medium containing 50 μg/ml gentamicin was added to wells to kill extracellular bacteria.

### Cell death assays, bacterial luminescence and IL-18 ELISA

Cells were plated in 96 well plate with white sides (Corning 3610). *S. flexneri* or *S.* Typhimurium infections were performed as described above, sytox green (Invitrogen) was added at the same time as gentamicin at a final concentration of 1.5-3 nM in 200 μl per well of 96 well plate. For lysed control wells, 1% triton X-100 was added to media containing sytox green and gentamicin. Sytox green fluorescence and bacterial luminescence (*S. flexneri* only) was measured every 30 minutes starting at 1 hpi. Fluorescence and luminescence were measured using Enspire 2300 multilabel plate reader (Perkin Elmer). At 3 hpi, 100 μl supernatant was removed from each well, 30 μl was used for LDH assay and the rest was frozen at −80°C until running IL-18 ELISA. LDH was measured by CytoTox-ONE homogenous membrane integrity assay (Promega). Briefly, 30 μl supernatant was mixed with 30 μl CytoTox-ONE reagent, incubated for 7 minutes at room temperature, then measured on Enspire 2300 plate reader. IL-18 was measured using IL-18 ELISA matched antibody pair (Thermo Fisher BMS267-2MST) following manufacturer’s instructions.

### Western blot

Cells were plated in a 12 well plate and primed with IFNγ overnight (if applicable). Next day, cells were washed twice with cold PBS, then lysed in 90 μl RIPA buffer (Millipore Sigma) containing protease inhibitors (Millipore Sigma P8340) on ice for 30 minutes. Lysates were clarified through centrifugation at 20,000×g for 10 minutes at 4°C. Clarified lysate was mixed with Laemmli sample buffer (BioRad) containing beta-mercaptoethanol and boiled for 5 minutes. Samples were run on 4-20% mini-PROTEAN TGX Stain-free gel (BioRad) with all blue protein standards (BioRad) as a ladder. Running buffer contained 25 mM Tris base, 190 mM glycine, 0.1% SDS. Gel was transferred to PVDF using BioRad Trans-blot turbo. Following transfer, membrane was dried then incubated with primary antibody in Tris-buffered saline containing 0.1% tween 20 (TBST) and 5% non-fat milk. Blots were incubated in primary antibody either 1 hour at room temperature, or overnight at 4°C. Following primary antibody, blots were washed three times in TBST, then incubated with secondary antibody in 5% milk in TBST for 45-60 minutes. Blots were then washed five times in TBST then incubated with Clarity ECL substrate (BioRad) or SuperSignal West Femto ECL substrate (Thermo Fisher). Blots were imaged using an Azure 500 imaging system. Primary antibodies were used at the following concentrations: GBP1 (Abcam ab131255, 1:5000), GBP2 (Santa Cruz sc271568, 1:200), GBP4 (Proteintech 17746-1-AP, 1:10,000), GBP5 (Cell Signaling 67798, 1:5,000), GAPDH (Abcam ab9485, 1:10,000), mCherry (Abcam ab183628, 1:10,000). Secondary antibodies: goat anti-rabbit HRP (BioRad 1706515 or Invitrogen 65-6120, 1:5,000), anti-mouse HRP (Santa Cruz sc-525409, 1:5,000).

### Immunofluorescence microscopy

Cells were plated on glass coverslips in 24 well plates and primed overnight with IFNγ. The next day cells were infected with *Shigella* at an MOI of 5. At the indicated time points after infection, media was aspirated from wells and cells were fixed in 4% paraformaldehyde in PBS for 15 minutes at room temperature. Cells were washed three times with PBS, then permeabilized with 0.1% triton X-100 in PBS for 15 minutes. Cells were blocked for 30 minutes in PBS containing 5% BSA and 2.2% glycine. Anti-GBP1 (Abcam ab131255, 1:150 dilution) was added in blocking buffer for 1 hour at room temperature. Cells were washed three times with PBS containing 0.05% triton X-100, then incubated with donkey anti-rabbit IgG Alexa Fluor 568 (Thermo Fisher A10042, 1:1000 dilution) and Hoechst 33258 (1μg/mL) for 45 minutes in blocking buffer. Cells were then washed three times with PBS containing 0.05% triton X-100 and mounted on glass slides. For experiments without any antibody staining, following fixation, cells were washed three times in PBS then incubated with 1 μg/mL Hoechst for 20 minutes in PBS, washed twice with PBS and mounted on glass slides. Coverslips were mounted with mounting media containing 9 parts Mowiol solution (100 mM Tris-HCl, pH 8.5, 25% glycerol, 125 μg/mL Mowiol 4-88) and 1 part PPD solution (0.1 mg/mL 1,4-Phenylenediamine dihydrochloride in water). Mounting media was allowed to harden overnight at room temperature. Images were acquired using a Zeiss Axio Observer Z1 microscope using Zeiss Plan-Apochromat 63×/1.4 oil objective, or a DeltaVision Elite Deconvolution microscope with UPLSAPO 100×/1.40 oil objective. Images were processed with DeltaVision software for deconvolution, or with Fiji.

### Time-lapse microscopy of infected cells

700,000 A459 GBP1-KO pInducer-mCherry, -GBP1, or GBP1^3R^ cells were plated on glass bottom 10 mm microwell dishes and treated with 2 µg/ml aTc and 100 U/ml IFNγ for 20-22 h to stimulate mCherry fusion protein expression. Cells were infected with GFP-expressing *S. flexneri* Δ*ipaH9.8*Δ*ospC3* as described above with the following alterations. After incubating cells at 37 °C and 5% CO_2_ following centrifugation for 10 min at 700 x g at room temperature, cells were washed three times with 2 ml HBSS. After washing, 2 ml infection media supplemented with 150 nM sytox blue (Invitrogen) was added to each dish to stain cells undergoing pyroptosis. Time-lapse images of infected cells were acquired every 3 min with a Zeiss 880 AiryScan Fast Inverted Confocal on AxioObserver Z1 microscope using a Zeiss Plan-Apochromat 63×/1.4 oil objective, with stage incubator set to 37°C and 5% CO_2_ buffering. All images were processed with Fiji.

### LPS electroporation

Electroporation was done using the Neon Transfection System (Thermo Fisher). Cells were plated in a 6 well plate and were either left untreated or were treated with 100 U/ml IFNγ overnight. The next day, cells were trypsinized, washed twice with PBS and the cell number was determined. Cells were pelleted and diluted in Neon buffer R to 5,000 cells/µl. *E. coli* O55:B5 LPS (Invivogen tlrl-pb5lps) was diluted in Neon R buffer to 500 μg/ml, 100 μg/ml and 50 μg/ml. Cell and LPS dilutions were gently mixed and 500,000 cells were electroporated with either 2.5 μg, 0.5 μg or 0.25 μg LPS in a 100 µl Neon pipette tip using electrolytic buffer E2 and 2 pulses of 1005 V for 35 ms. After electroporation prewarmed infection media supplemented with 3 µg/ml propidium iodide was quickly added to cells and 50,000 cells were plated per well in black tissue culture treated 96-well plates in triplicates. Non-electroporated cells left untreated or treated with 1% triton served as live and dead controls. Fluorescence was measured using an Enspire 2300 multilabel plate reader (Perkin Elmer) at 1, 2, and 4 hours post electroporation. After measuring fluorescence, 100 μl supernatant was removed from each well and was frozen at −80°C until running IL-18 ELISA. IL-18 was measured using IL-18 ELISA matched antibody pair (Thermo Fisher BMS267-2MST) following manufacturer’s instructions.

### LPS transfection

Transfection protocol was modified from Santos et al. (25) A549 cells were seeded in a 96 well plate, 2.5×10^4^ cells per well. The next day, LPS transfection mixture was prepared-for each well 75 μl optimem (31985–062), 1 μl lipofectamine 2000 (Invitrogen), and either 1 μg or 0.1 μg *E. coli* O55:B5 LPS (Invivogen tlrl-pb5lps). LPS and lipofectamine mixture was incubated for 20 minutes before adding to cells. During incubation, media in 96 well plate was aspirated and 75 μl optimem containing 0.5 μM sytox green (Invitrogen) was added to each well. Lysed control wells contained 75 μl optimem, 0.5 μM sytox green, and 1% triton X-100. 75 μl lipofectamine and LPS mixture was added on top of media containing sytox green.

### Plasmids

All GBP expression plasmids were in the lentiviral pInducer20 backbone (58) containing a C-terminal mCherry (GBP4) or N-terminal mCherry (all other constructs). Full plasmid sequences are provided as a supplemental file. For experiments using these plasmids, expression was induced by adding 1 μg/ml anhydrotetracycline (Takara) overnight in cell culture media. When applicable, anhydrotetracycline was added at the same time as IFNγ. For *S. flexneri* luminescence experiments, strains were transformed with ilux pGEX(-), which was a gift from Stefan Hell (Addgene plasmid #107879; http://n2t.net/addgene:107879; RRID:Addgene_107879) *S. flexneri* was transformed with pGFPmut2 (59) for microscopy experiments.

### Lentivirus production and cell line complementation

293T cells were plated in a 6 well plate with 1×10^6^ cells per well in 2ml. The next day cells were transfected using TransIT 293 transfection reagent (Mirus), following manufacturer’s instructions. Each well was transfected with 1 μg pInducer plasmid, 750 ng pSPAX2 (Addgene), 250 ng VSVG. At 24 hours post transfection, media was removed and 3 ml fresh media added. Supernatant containing virus was collected at 48 and 72 hours post infection, and filtered through 0.45 μm nylon filters (corning). Virus was frozen at −80°C until use.

For transduction, A549 cells were trypsinized and resuspended in media containing 10 μg/mL polybrene (Millipore Sigma) to a concentration of 3.33×10^4^ cells/ml. In 6 well plate, 250 μl lentivirus supernatant (described above) and 1.5 ml diluted cells were added to each well. Cells were incubated for 48-72 hours, then 2 mg/ml geneticin was added for approximately 10 days to select for cells containing the pInducer plasmid.

### Expression, purification, and prenylation of recombinant protein

Recombinant GBP1, GBP5, and FTase were expressed and purified as described previously(34). N-terminally His_6_-tagged GBP1 and GBP2 were expressed in *E. coli* strain BL21 CodonPlus (DE3) RIL from bacterial vector pQE-80L. N-terminally His_10_-tagged GBP5 was expressed in *E. coli* strain Rosetta (DE3) pLysS from bacterial vector pQE-80L. N-terminal His_6_-tagged farnesyltransferase (FTase) and geranylgeranyltransferase (GGTase) were expressed in *E. coli* strains Rosetta (DE3) pLysS and BL21 CodonPlus (DE3) RIL from pRSF-Duet1 vector. Bacteria were cultivated in terrific broth media (GBPs) or terrific broth media supplemented with 60 µM ZnCl_2_ (Ftase, GGTase) and grown at 37 °C and 90 rpm to an OD_600_ of 0.4–0.8. After decreasing the temperature to 20 °C, protein expression was induced with 100 µM IPTG. For FTase and GGTase expression, an additional 0.5 mM ZnCl_2_ was added to the culture. Bacteria were harvested after 16–18 h at 3000 x g for 15 min at 4°C (Sorvall LYNX 6000 centrifuge, F9-6×1000 LEX rotor, Thermo Fisher Scientific).

Buffer compositions for the purification of recombinant GBPs and FTase or GGTase differed in the use of 50 mM HEPES, pH 7.8 for FTase and GGTase instead of 50 mM Tris-HCl, pH 7.9 for GBPs. Harvested bacteria were resuspended in buffer A (50 mM Tris-HCl, pH 7.9, 500 mM NaCl, 5 mM MgCl_2_) supplemented with 1 mM phenylmethylsulfonyl fluoride to inhibit proteases and sonicated on ice at 30% amplitude pulsing at 1 sec on/ 1 sec off for a total of 10 min (Ultrasonic homogenizer Sonoplus HD 2200, Bandelin) with the temperature of the resuspension kept below 8 °C. Cell debris was removed from lysate containing soluble protein by centrifugation at 35,000 x g and 4 °C for 45 min (Sorvall LYNX 6000 centrifuge, F21-8×50y rotor, Thermo Fisher Scientific). Proteins were further purified by immobilized metal affinity chromatography (IMAC) followed by size-exclusion chromatography (SEC). All chromatography columns were connected to ÄKTA Purifier or Prime systems (GE Healthcare Life Sciences). After loading of soluble proteins, the IMAC column (30 ml HisPur Cobalt Resin, 30 ml) was sequentially washed with 5-10 column volumes (CVs) buffer A and 3-4 CVs buffer B_10_ (50 mM Tris-HCl, pH 7.9, 150 mM NaCl, 5 mM MgCl_2_, 10 mM imidazole). His-tagged protein was eluted with 2 CVs buffer B_150_ (50 mM Tris-HCl, pH 7.9, 150 mM NaCl, 5 mM MgCl_2_, 150 mM imidazole). GBP containing fractions from IMAC were pooled and precipitated by slowly adding 3 M (NH4)_2_SO_4_. (NH4)_2_SO_4_ protein precipitates were dissolved in buffer C (50 mM Tris-HCl, pH 7.9, 150 mM NaCl, 5 mM MgCl_2_) and loaded on a with buffer C equilibrated SEC column (Superdex 200 26/ 60, 320 ml) to remove (NH_4_)_2_SO_4_ and aggregated protein from monomeric protein. FTase and GGTase containing fractions from IMAC were pooled, concentrated via ultra-filtration using Vivaspin 20 centrifugal columns (10 kDa cut-off, Sartorius), and loaded on the SEC column (Superdex 200 26/ 60, 320 ml) to isolate monomeric protein. Fractions containing monomeric GBPs, FTase, and GGTase were pooled, concentrated via ultra-filtration using 10 kDa cut-off Vivaspin 20 centrifugal columns, frozen in liquid nitrogen, and stored at −80°C.

GBP1 was farnesylated *in vitro* as described previously(36). Monomeric GBP1, GBP2, and GBP5 were incubated for 16 h at 4°C in glass vials with farnesyl pyrophosphate (FPP) or geranylgeranyl pyrophosphate (GGPP) and FTase or GGTase in buffer D (50 mM Tris-HCl, pH 7.9, 5 mM MgCl_2_, 150 mM NaCl, 10 µM ZnCl_2_). For farnesylation, 60 µM GBP1 was supplemented with 150 µM FPP and 1.25 µM FTase in a total volume of 4 ml. For geranylgeranylation, 60 µM GBP2 or GBP5 were supplemented with 150 µM GGPP and 5 µM GGTase in a total volume of 4 ml. Reaction mixtures were supplemented with (NH_4_)_2_SO_4_ (3 M stock solution) to a final concentration of 1.25 M and loaded on a hydrophobic interaction chromatography (HIC) column (Butyl FF 16/10, 20 ml), equilibrated with buffer E (50 mM Tris-HCl, pH 7.9, 5 mM MgCl_2_, 1.2 M (NH_4_)_2_SO_4_). For GBP1, after loading, the HIC column was sequentially washed with 2 CVs buffer E and with 2 CVs of buffer E with its initial (NH_4_)_2_SO_4_ concentration decreased to 60%. Farnesylated GBP1 was separated from non-farnesylated GBP1 by decreasing the (NH_4_)_2_SO_4_ concentration further in a continuous gradient over 3 CVs from 60% to 45% (NH_4_)_2_SO_4_ (elution of farnesylated GBP1) followed by a continuous gradient over 3.75 CVs from 45% to 25% (NH_4_)_2_SO_4_ (elution of non-farnesylated GBP1). For GBP2 and GBP5, after loading, the HIC column was sequentially washed with 2 CVs buffer E and with 2 CVs of buffer E with its initial (NH_4_)_2_SO_4_ concentration decreased to 80%. Geranylgeranylated GBP2 and GBP5 were eluted from the HIC by decreasing the (NH_4_)_2_SO_4_ concentration from 80% to 0% in a continuous gradient over 20 CVs. Fractions with prenylated GBPs were pooled, concentrated via ultra-filtration using 10 kDa cut-off Vivaspin 20 centrifugal columns and further purified by SEC to isolate monomeric protein. Following SEC, fractions with monomeric prenylated GBPs were pooled, concentrated via ultra-filtration, frozen in liquid nitrogen, and stored at −80°C. Concentrations of proteins were calculated according to Lambert–Beer law, using absorption at 280 nm in buffer F (6 M guanidine hydrochloride, 20 mM potassium phosphate, pH 6.5) and respective molar absorption coefficients (GBP1 43,240 M^-1^cm^-1^, GBP2 52,050 M^-1^cm^-1^, GBP5 46,005 M^-1^cm^-1^, FTase 158,235 M^-1^cm^-1^, GGTase 138,170 M^-1^cm^-1^).

### Labeling of recombinant proteins with fluorescent dyes

After exchanging buffer C to buffer G (50 mM Tris-HCl, pH 7.4, 150 mM NaCl, and 5 mM MgCl_2_), recombinant GBPs were incubated with Alexa Fluor 488 C_5_ maleimide dye or Alexa Fluor 647 C_2_ maleimide dye (Invitrogen) in a ratio of 1:1 or 1:2 on ice for 10–20 min. Labeling reactions were stopped by changing buffer G to buffer C supplemented with 2 mM DTT via ultra-filtration using 10 kDa cut-off Vivaspin Turbo 4 centrifugal columns. Concentrations of proteins and labeling efficiencies were calculated according to Lambert-Beer law, using absorptions at 280 nm, 491 nm, and 651 nm in buffer C, respective molar absorption coefficients (GBP1 45,840 M^-1^cm^-1^, GBP2 53,860M^-1^cm^-1^, GBP5 45,380 M^-1^cm^-1^, Alexa Fluor 488 71,000 M^-1^cm^-1^, Alexa Fluor 647 268,000 M^-1^cm^-1^), and correction factors for fluorescent dyes (Alexa Fluor 488 0.11, Alexa-Fluor647 0.03). Labeling efficiencies for Alexa Fluor 488-labeled proteins ranged from 44% to 120%. Labeling efficiencies for Alexa Fluor 647-labeled proteins ranged from 13% to 49%.

### Binding of protein to bacteria

*S. flexneri* expressing RFP was streaked on an agar plate, then single colonies were picked and grown overnight at 37 °C shaking, in 5 ml TSB. The next day 175 μl overnight culture was diluted into 5 ml TSB and grown for 1 hour 20 minutes. 2.5 ml bacterial culture was pelleted, washed with 1 ml PBS, and resuspended in 1 ml 4% formaldehyde in PBS, pH 7.4 for 20 min to fix. Formaldehyde-fixed bacteria were washed twice with PBS and resuspended in PBS supplemented with 0.03% NaN3. Final concentrations for *in vitro* binding experiments were 10^5^-3 x 10^6^ bacteria/ml, 5 µM GBP, and 2 mM GTP. Bacteria were diluted in buffer C, supplemented with 50 µM BSA, and the dilution was applied to the cover slide of a glass bottom 10 mm microwell dish. Following centrifugation for 1 min at 3,000 x g bacteria were incubated for 5 min at 25 °C on the temperature-controlled microscope stage. Alexa Fluor-labeled GBPs were diluted in buffer C supplemented with 50 µM BSA, mixed with GTP, and the mixture was added to bacteria at t = 0 min. The samples were gently mixed, and images were collected every 1.5 min. After recording time-lapse images for 60 min different field of views were imaged for quantification. Imaging was performed on a Zeiss 880 Airyscan Fast Inverted Confocal on Axio Observer Z1 microscopes using Zeiss Plan-Apochromat 63×/ 1.4 oil objectives. All images were processed with Fiji.

### LPS aggregation

Final concentrations for LPS aggregation microscopy experiments were 5 µM recombinant GBP, 50 µg/ml LPS, and 2 mM GTP. Alexa Fluor 568-labeled *E. coli* O55:B5 LPS (Invitrogen) was diluted in buffer C, supplemented with 50 µM BSA, and the dilution was applied to the cover slide of a glass bottom 10 mm microwell dish. Following centrifugation for 1 min at 3,000 x g bacteria were incubated for 5 min at 25 °C on the temperature-controlled microscope stage. GBPs were diluted in buffer C supplemented with 50 µM BSA, mixed with GTP, and the mixture was added to LPS at t = 0 min. The samples were gently mixed, and after 20 min, different field of views were imaged for quantification. Imaging was performed on a Zeiss 880 Airyscan Fast Inverted Confocal on Axio Observer Z1 microscopes using Zeiss Plan-Apochromat 63×/ 1.4 oil objectives. All images were processed with Fiji and LPS areas and numbers were quantified with the integrated analyze particle tool.

### Dynamic light scattering

Dynamic light scattering (DLS) experiments of recombinant GBPs and LPS was performed with Delsa Max Pro instrument (Beckman Coulter). GBPs were diluted in buffer C or buffer C (apo) supplemented with 300 µM AlF_X_, 10 mM NaF, and 250 µM GDP (GDP·AlF_X_) to a final concentration of 5 µM in the presence or absence of 0.05 mg/ml *E. coli* O55:B5 LPS. Samples were incubated at room temperature (RT) for 1 hour prior starting DLS measurements in a temperature-controlled cuvette set to RT. The particle size was measured over three measurements, each consisting of ten runs. The number-weighted radius (R_n_) of the particles was determined with the manufacturer’s software. A series of controls were also measured to establish the underlying particle sizes of the filtered aqueous buffers and the protein solutions. These were buffer C, buffer C supplemented with GDP·AlF_X_, and 5 µM bovine serum albumin (BSA) or 0.05 mg/ml LPS in the respective buffers.

### Absorbance-based light scattering

Absorbance-based light scattering experiments were performed with a Specord200 UV/Vis spectrophotometer (Analytik Jena) as described previously(35). GBP1, GBP2, and GBP5 were diluted in buffer C supplemented with 50 µM BSA to a final concentration of 5 µM. Protein dilutions were incubated in a temperature-controlled cuvette at 25°C for 5 min. GTP was added to protein dilutions at a final concentration of 2 mM to start GBP self-assembly. Polymerization of GBPs was followed as absorbance signal at 350 nm over time. To determine formation of mixed polymers, absorbance of 5 µM GBP1 in the presence of 5 µM GBP1, GBP2, or GBP5 following GTP addition was monitored (final GBP concentration 10 µM, polymerization of 5 µM GBP1 is shown as control).

### Native polyacrylamide gel electrophoresis

Native GBP-LPS complexes were analyzed with native polyacrylamide gel electrophoresis (NPAGE). Recombinant GBP1, GBP2, GBP5, BSA (control), and *E. coli* O55:B5 LPS were diluted in buffer C or buffer C supplemented with 300 µM AlF_X_ and 10 mM NaF. Protein and LPS dilutions were mixed and GDP was added to induce complex formation. Final concentrations were 5 µM protein, 250 µM GDP, 1 mg/ml, 0.1 mg/l, or 0.01 mg/ml LPS (Fig. 7B), and 2 mg/ml to 0.008 mg/ml (1:1 dilutions, Fig. 7C and Fig. S6C). After incubation at RT for 30 min, samples were mixed 1:1 with 2 x native sample buffer (BioRad) and loaded on a 4-20% precast protein gel (BioRad) for electrophoresis in Tris/Glycine running buffer (BioRad) at 80 V for 15 min followed by 180 V for 1 h. After electrophoresis, gels were fixed overnight in 60% methanol and 10% acetic acid. The next day, gels were rehydrated in 3% acetic acid and successively stained using Pro-Q Emerald 300 Lipopolysaccharide Gel Stain Kit (Invitrogen) and Coomassie Brilliant Blue R-250. LPS staining and Coomassie staining were visualized with an Azure 500 Biosystem using excitation at 365 nm and emission at 595 nm or the UV320 Coomassie detection program (utilizing the orange tray), respectively. All gel images were processed with Fiji. Gel bands were analyzed with the built-in Fiji tool.

## Acknowledgments

We thank Dr. Edward Miao for sharing *Salmonella* strains with us and Dr. Gerrit Praefcke for sharing the pRSF-Duet1-FTase plasmid. This work was supported by National Institutes of Health grants AI139425 (to J.C.), AI064285 (C.F.L.) and AI28360 (C.F.L.); by Deutsche Forschungsgemeinschaft (DFG, German Research Foundation) Research Fellowship 427472513 (to M.K.) and by research grant HE2679/6-1 (to C.H). J.C. holds an Investigator in the Pathogenesis of Infectious Disease Award from the Burroughs Wellcome Fund. This work was performed in part at the Duke University Light Microscopy Core Facility (LMCF) and Duke Functional Genomics. We thank LMCF members Drs. Benjamin Carlsen, Yasheng Gao, and Lisa Cameron for providing training and Dr. So Young Kim for generating knockout cell lines.

---

**Movie 1. GBP2 and GBP5 fail to encapsulate bacteria on their own.** Recombinant Alexa Fluor 488-labeled GBP1, GBP2, or GBP5 were supplemented with GTP and added to formaldehyde-fixed RFP-expressing *S. flexneri*.

**Movie 2. Both GBP2 and GBP5 associate with bacteria GBP1-dependently but only GBP2 incorporates in the microcapsule.** Recombinant Alexa Fluor 488-labeled GBP1, GBP2, or GBP5 mixed with Alexa-647-labeled GBP1 were supplemented with GTP and added to formaldehyde-fixed RFP-expressing *S. flexneri*.

**Movie 3. Pyroptosis of *S. flexneri* does not require GBP1 binding to bacteria.** GBP1^KO^ cells primed with 100 U/ml IFNγ overnight expressing mCherry (control), mCherry-GBP1, or mCherry-GBP1^3R^ (magenta) infected with GFP-expressing *S. flexneri* Δ*ospC3* Δ*ipaH9.8* (yellow). Dying cells are shown in blue (sytox blue).

**Figure S1.**
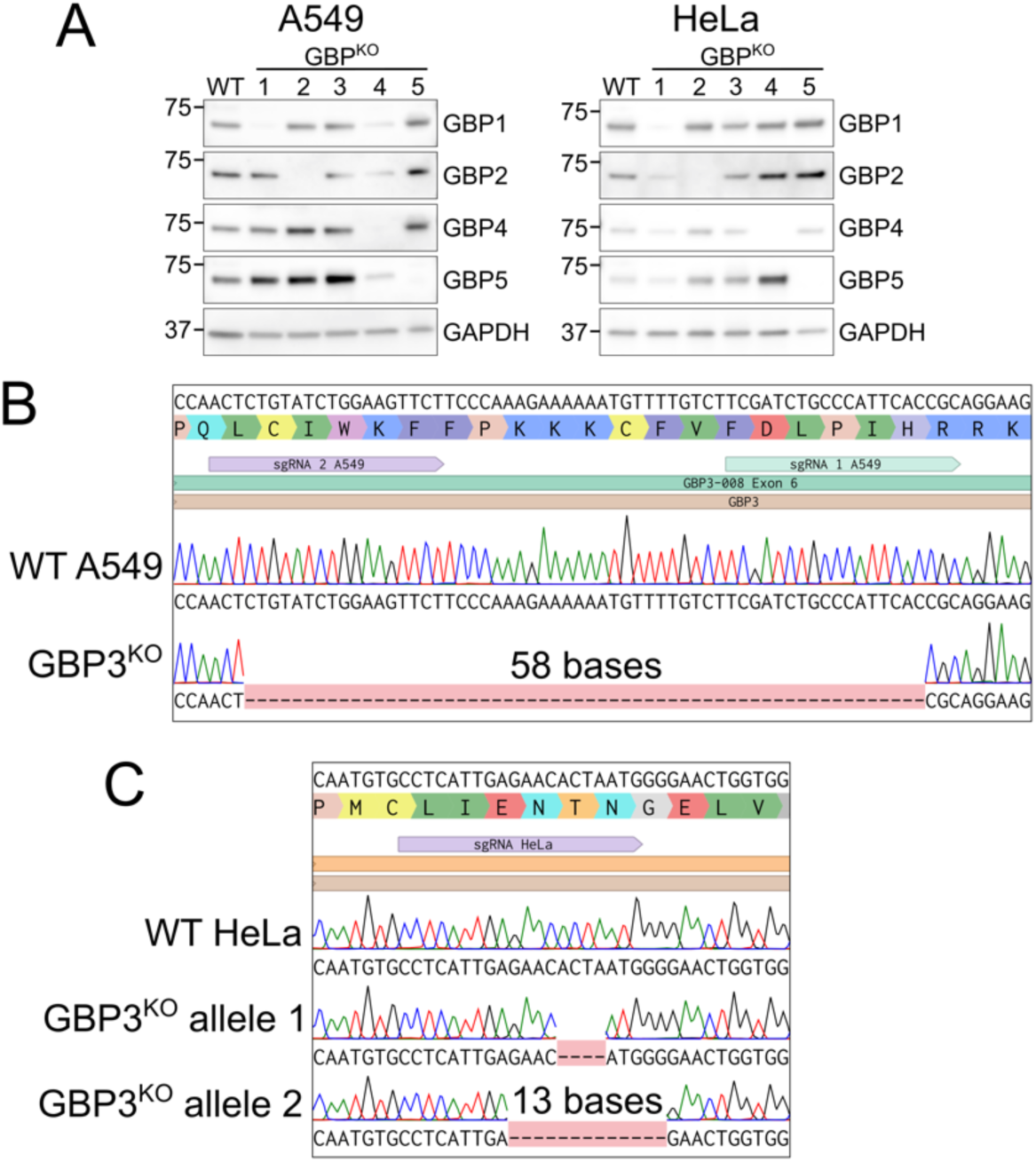
GBP1-5 were individually knocked out in A549 and HeLa cells. A549 and HeLa wildtype or CRISPR knockout clones were plated and primed with 100 U/ml IFNγ overnight. Cells were then lysed and GBP or GAPDH expression was analyzed by western blot. (B) Genomic DNA was extracted from wildtype and GBP3^KO^ A549 cells, PCR was used to amplify the edited region of the GBP3 gene, then PCR product was sequenced using sanger sequencing. Alignments were made using Benchling. (C) Genomic DNA was extracted from wildtype and GBP3^KO^ HeLa cells, PCR was used to amplify the edited region of the GBP3 gene, then PCR cloning was used to separate different alleles. Colony PCR and sanger sequencing were used to determine the sequence of each allele. Alignments were made using Benchling.

**Figure S2.**
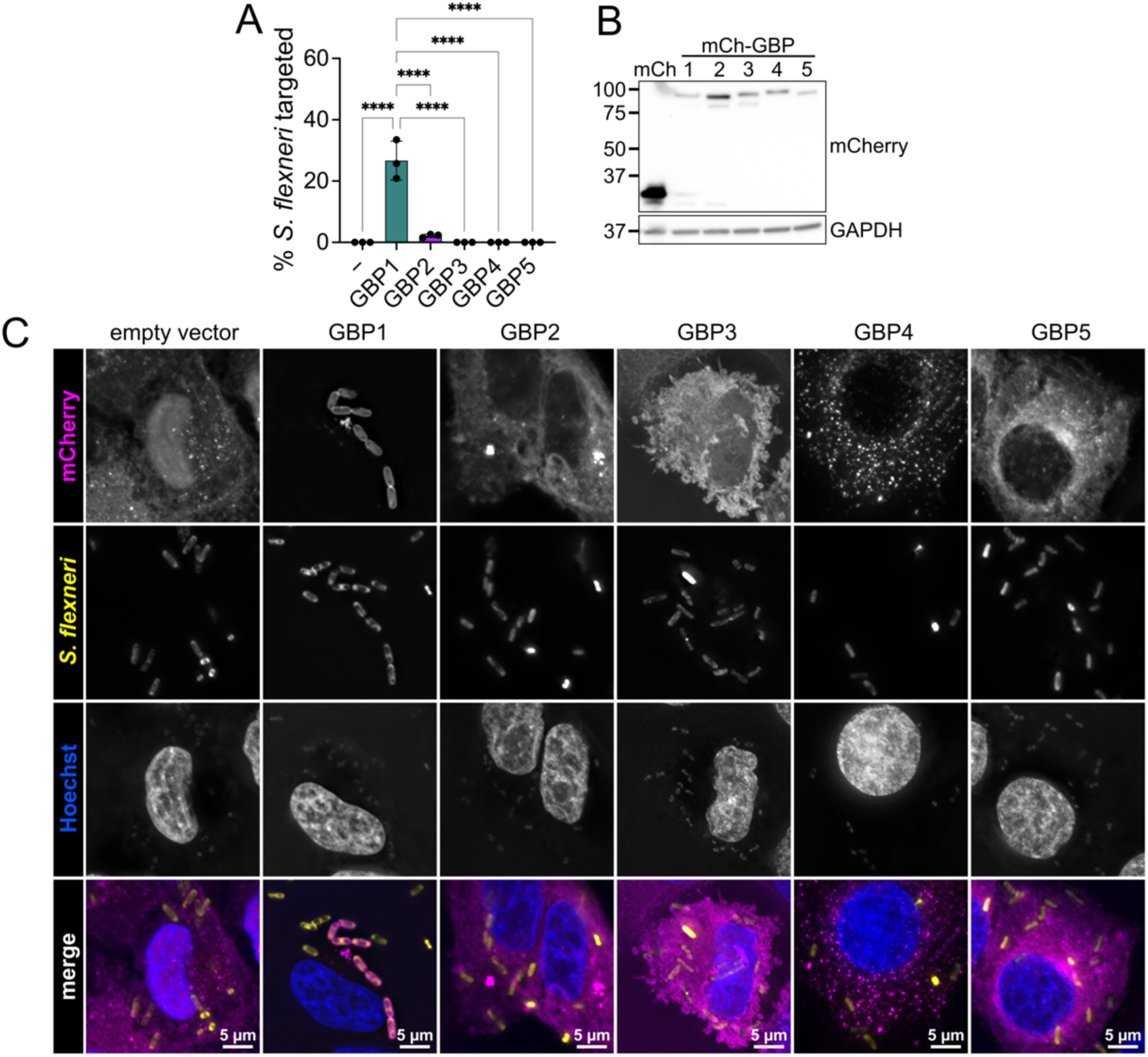
GBP1, but not GBP2-5, can bind bacteria in cells. (A) A549 GBP1^KO^ cells overexpressing mCherry or mCherry-GBPs were primed with 100 U/ml IFNγ overnight, then infected with *S. flexneri* Δ*ipaH9.8* expressing GFP. Cells were fixed at 2 hpi, images were taken at 63x by widefield microscopy, then total number of *S. flexneri* as well as mCherry positive *S. flexneri* were counted using ImageJ. *S. flexneri* with mCherry signal around at least 50% of the bacterial membrane were counted as targeted. (B) Expression levels of each mCherry constructs were tested using western blot. (C) A549 GBP1^KO^ cells overexpressing mCherry or mCherry-GBPs were primed with 100 U/ml IFNγ overnight, then infected with *S. flexneri* Δ*ipaH9.8* expressing GFP. Cells were fixed at 2 hpi and images were taken at 100x magnification by widefield microscopy. Images were deconvolved and z-projections are shown. (A) Graph shows averages from three independent experiments and is represented by mean ± SD. One-way ANOVA with Tukey’s multiple comparisons test was used. All significant comparisons are shown. **** = P < 0.0001. (B and C) Images are representative of three independent experiments.

**Figure S3.**
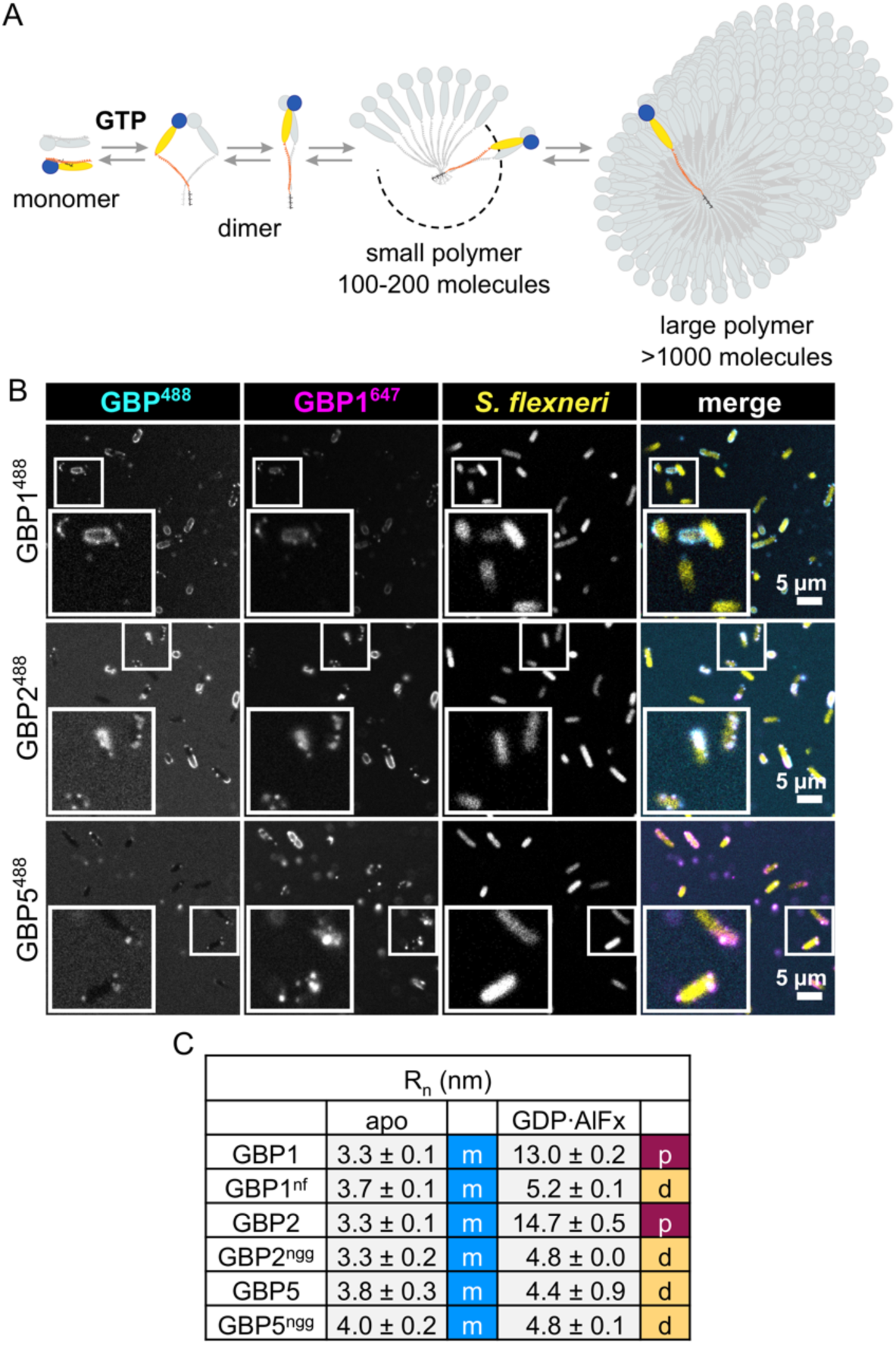
Mixed GBP1-GBP2 and GBP1-GBP5 polymers attach to the bacterial surface. (A) Mechanism of GBP1 polymerization. GBP1 exit its closed monomeric state, in which the farnesyl moiety is buried inside a hydrophobic pocket, and forms dimers in a GTP hydrolysis-dependent manner. GBP1 dimers can assemble into small polymers consisting of 100-200 molecules and continue their growth to form large polymers holding over 1000 molecules. (B) Confocal microscopy time frames of recombinant Alexa Fluor 488-labeled GBP1, GBP2, or GBP5 mixed with Alexa-647-labeled GBP1 supplemented with GTP at 5 min after addition to formaldehyde-fixed RFP-expressing *S. flexneri*. (C) Number-weighted mean radius (R_n_) of nucleotide-free (apo) or GDP·AlF_X_-bound nonisoprenylated (non-farnesylated = nf, non-geranylgeranylated = ngg) and isoprenylated GBPs determined in DLS experiments (values for data shown in Fig. 4G). monomer – m; dimer – d; polymer – p. (B) Representative images of three independent experiments. (C) Mean ± SD from three independent experiments.

**Figure S4.**
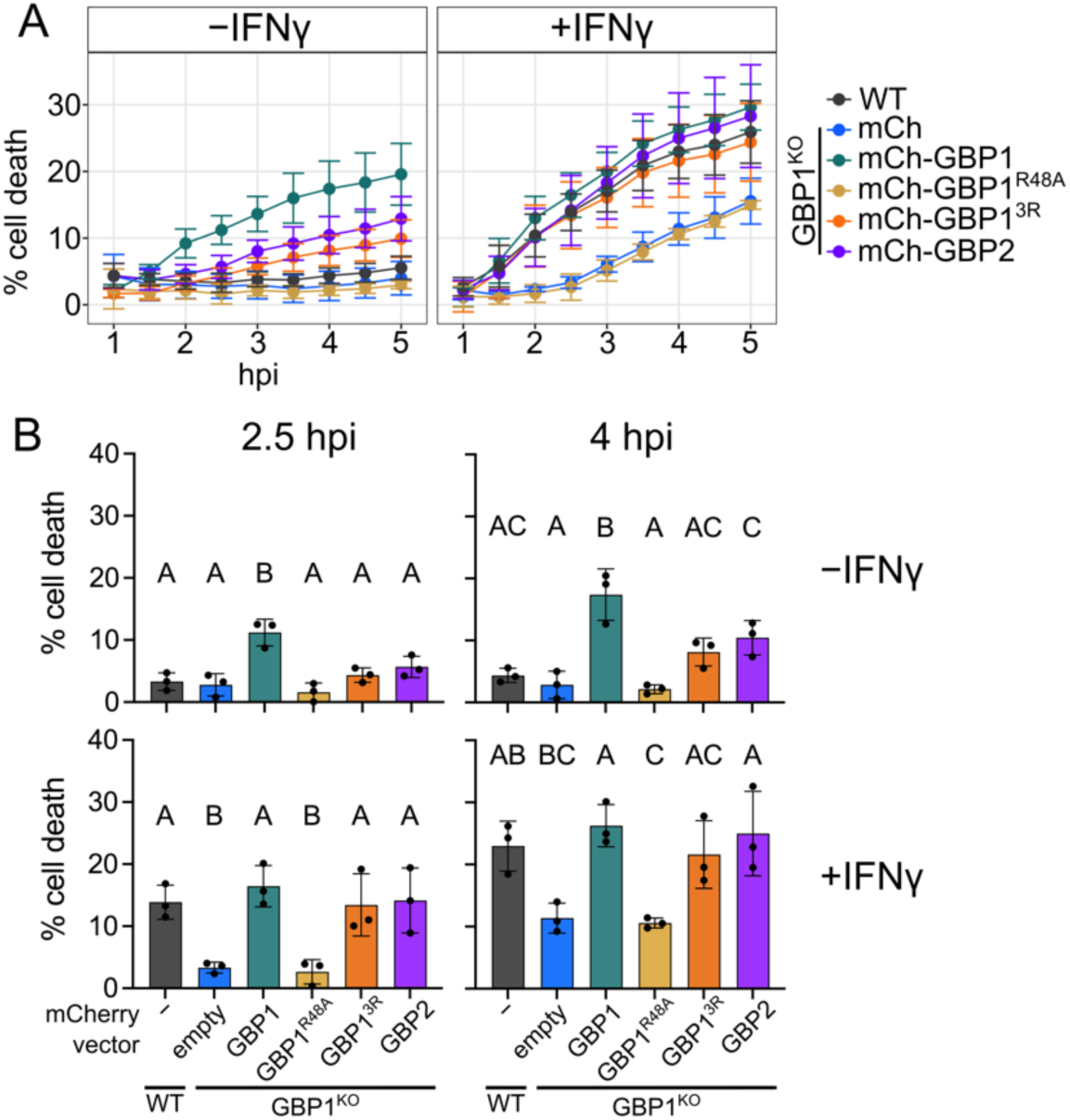
GBP1, GBP1^3R^, and GBP2 can rescue pyroptosis in GBP1^KO^ cells during *S.* Typhimurium infection. Wildtype A549 or GBP1^KO^ A549 cells overexpressing mCherry or mCherry-GBPs were plated and cells were unprimed or primed with 100 U/ml IFNγ overnight, then infected with *S.* Typhimurium. Cell death was monitored over time using sytox green fluorescence (A). For statistical analysis, the sytox green signal at 2.5 hours and 4 hours post infection was used (B). All graphs show averages from three independent experiments and are represented by mean ± SD. (B) One-way ANOVA with Tukey’s multiple comparisons test was used. Statistical comparisons are shown by letters, with bars sharing no matching letters being significantly different. * = P < 0.05, ** = P < 0.01, *** = P < 0.001, **** = P < 0.0001.

**Figure S5.**
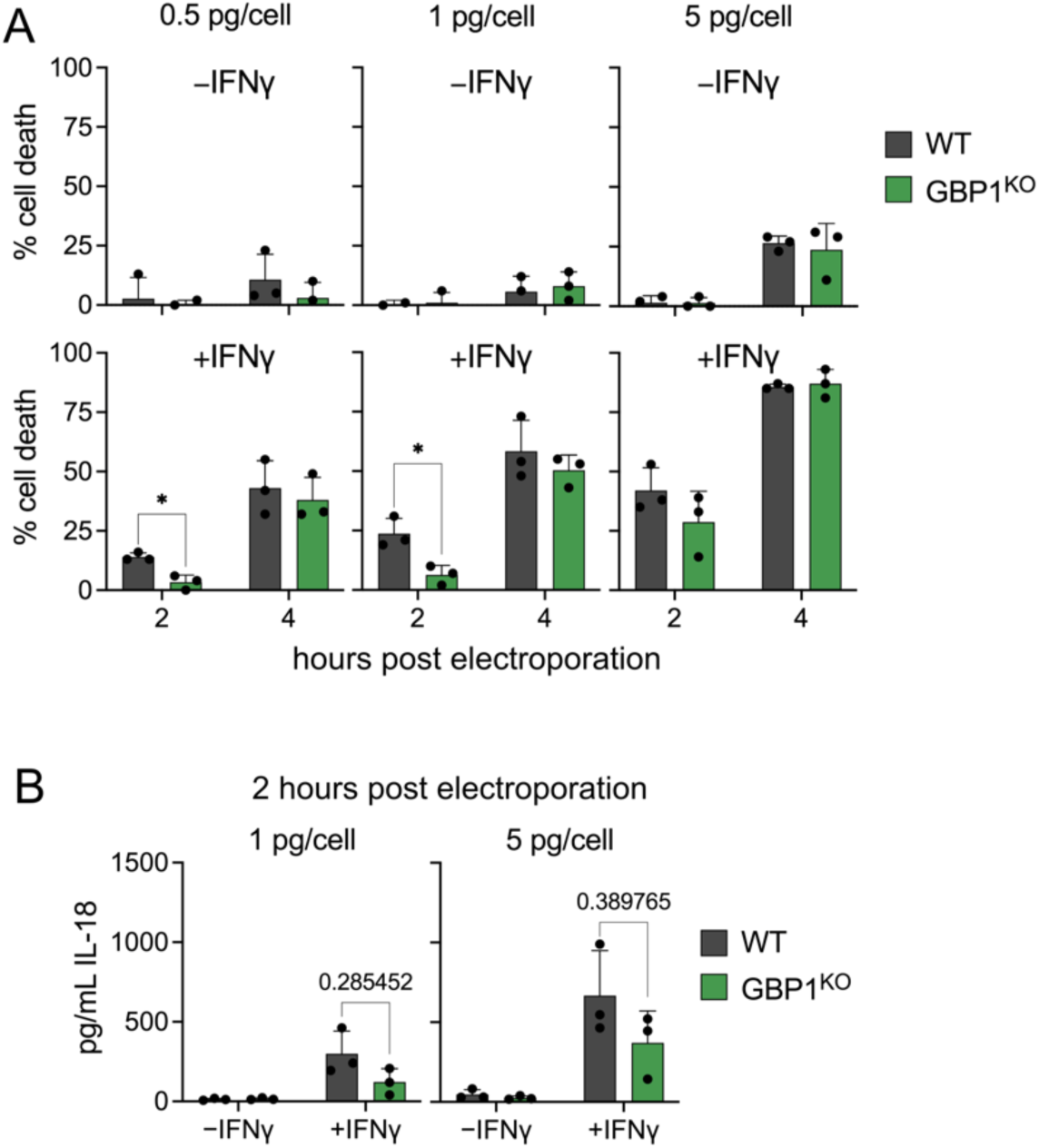
GBP1 is not required for pyroptosis in response to electroporated LPS. (A, B) Wildtype and GBP1^KO^ A549 cells unprimed or primed with 100 U/ml IFNγ overnight were electroporated with 0.5 pg/cell, 1 pg/cell, or 5 pg/cell *E. coli* O55:B5 LPS. Cell death was measured using sytox green fluorescence at 2 and 4 hours post electroporation (A). (B) IL-18 secretion was measured in supernatants taken at 2 hours post electroporation. All graphs show averages from three independent experiments and are represented by mean ± SD. Statistical significance was determined using multiple unpaired T tests with Welch correction, with multiple comparisons corrected with Holm-Šídák method. All significant comparisons are shown. Exact P values are shown in (B) for IFNγ primed cells. * = P < 0.05.

**Figure S6.**
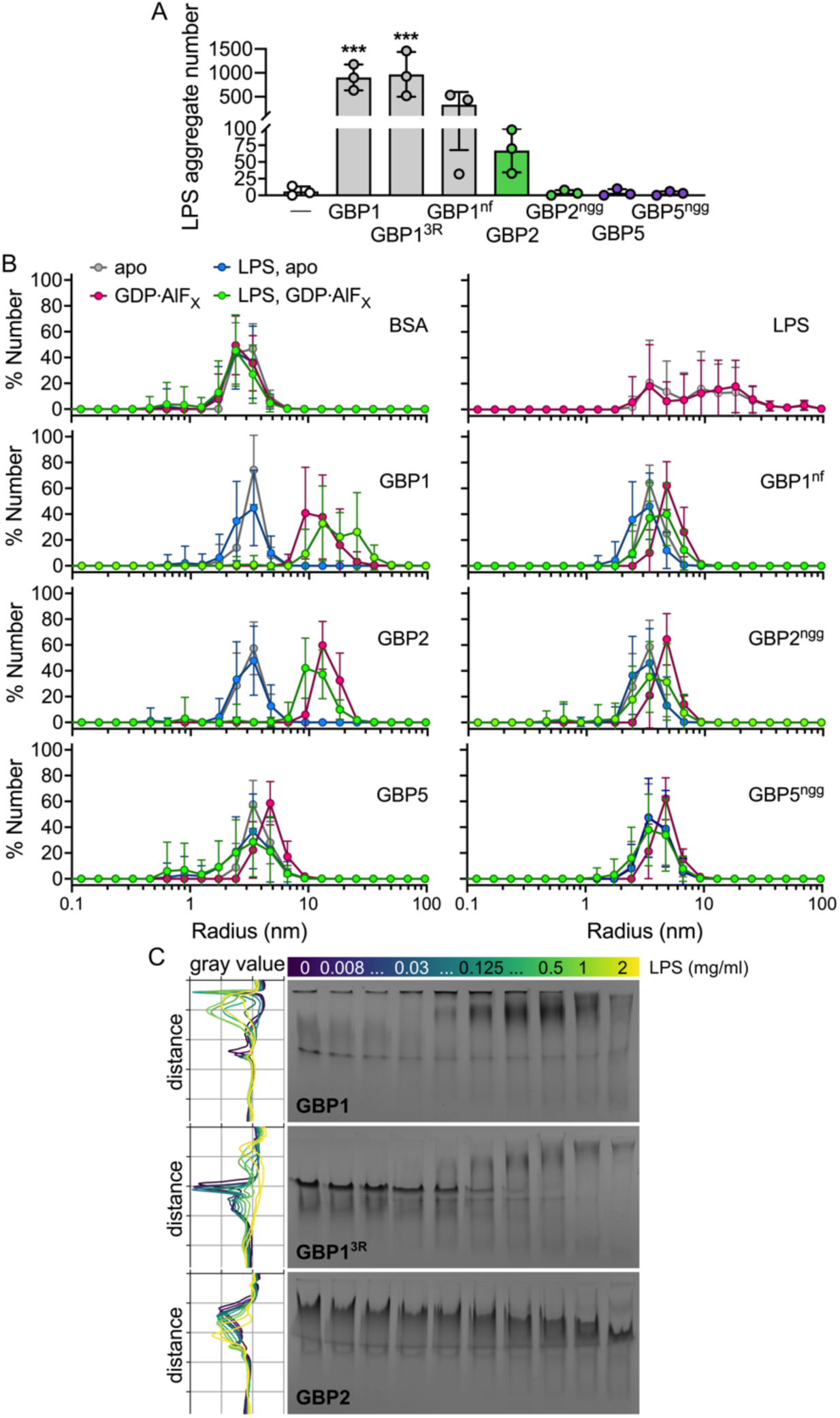
GBP2 binds directly to LPS. (A) Recombinant isoprenylated and nonisoprenylated GBPs were supplemented with GTP and added to Alexa Fluor 568-labeled *E. coli* O55:B5 LPS. LPS particles were analyzed from different fields of views taken after 20 min with Fiji, and the number of LPS aggregates was plotted. (B) Number-weighted mean radius (R_n_) of nucleotide-free (apo) or GDP·AlF_X_-bound nonisoprenylated (non-farnesylated - nf, non-geranylgeranylated - ngg) and isoprenylated GBPs in the presence and absence of LPS were determined in DLS experiments. (C) NPAGE of GBP1, GBP1^3R^, and GBP2 titrated with LPS (final concentrations 2 mg/ml to 0.008 mg/ml) supplemented with GDP·AlF_X_. Grey values for each gel lane were plotted with Fiji (GBP1 gel from Fig. 7C). All graph shows averages from three independent experiments and are represented by mean ± SD. (A) One-way ANOVA with Dunnett’s multiple comparisons test comparing to control (no GBP addition) was used. All significant comparisons are shown. *** = P < 0.001. (C) Representative NPAGEs from three independent experiments are shown.

Plasmid sequences shown below for GBP overexpression constructs. mCherry shown in red, GBPs in blue, point mutations in green.

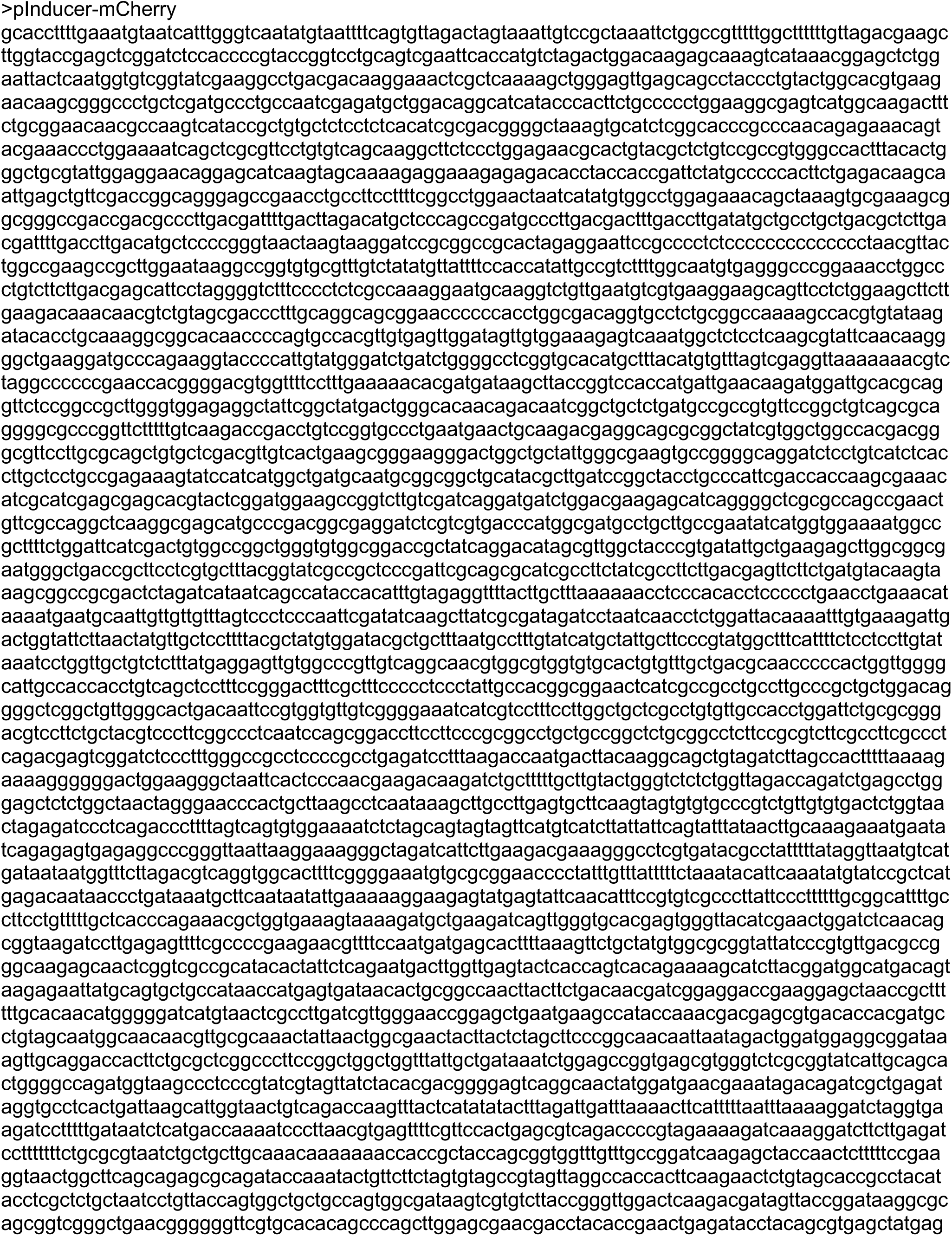
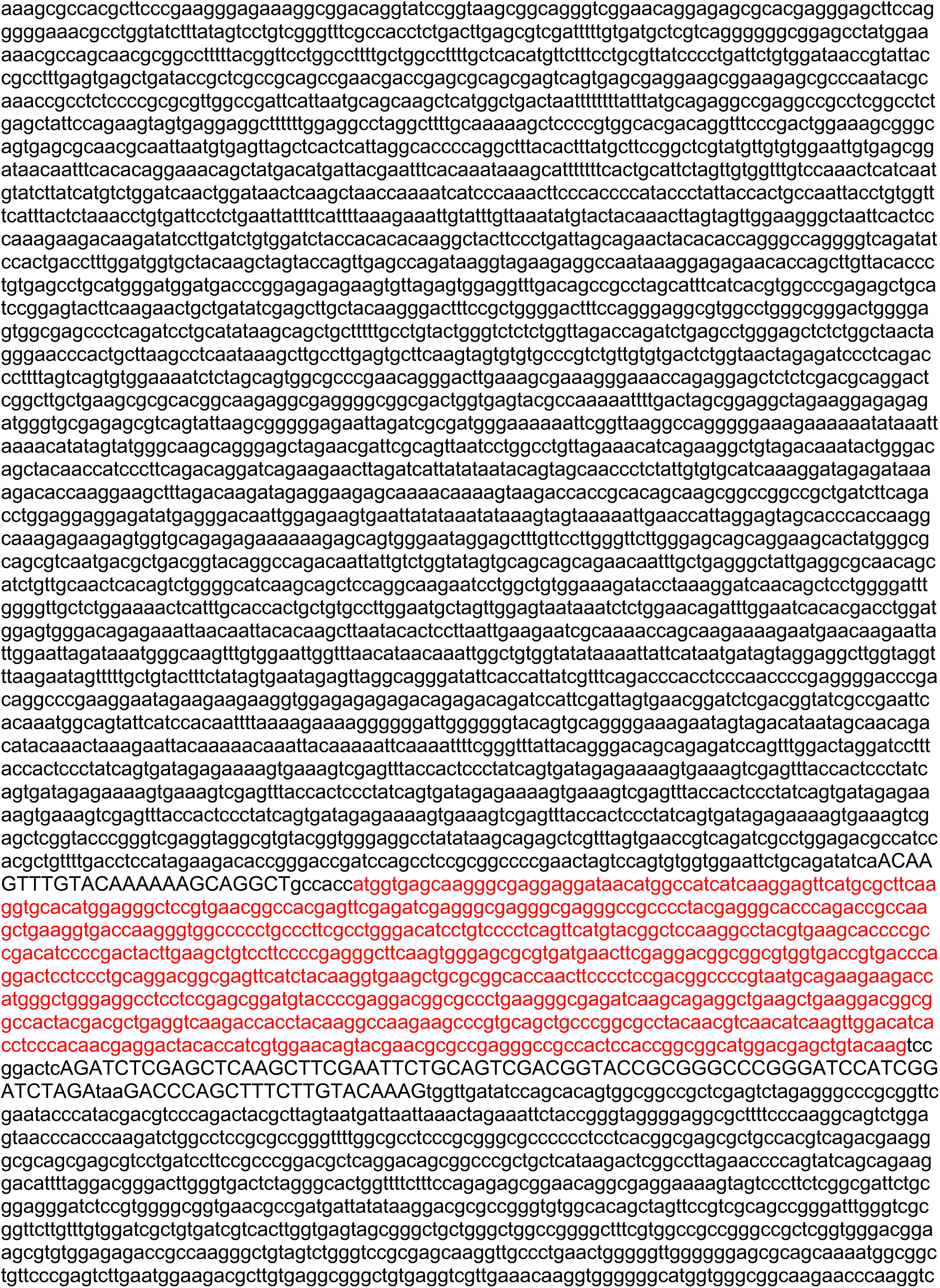
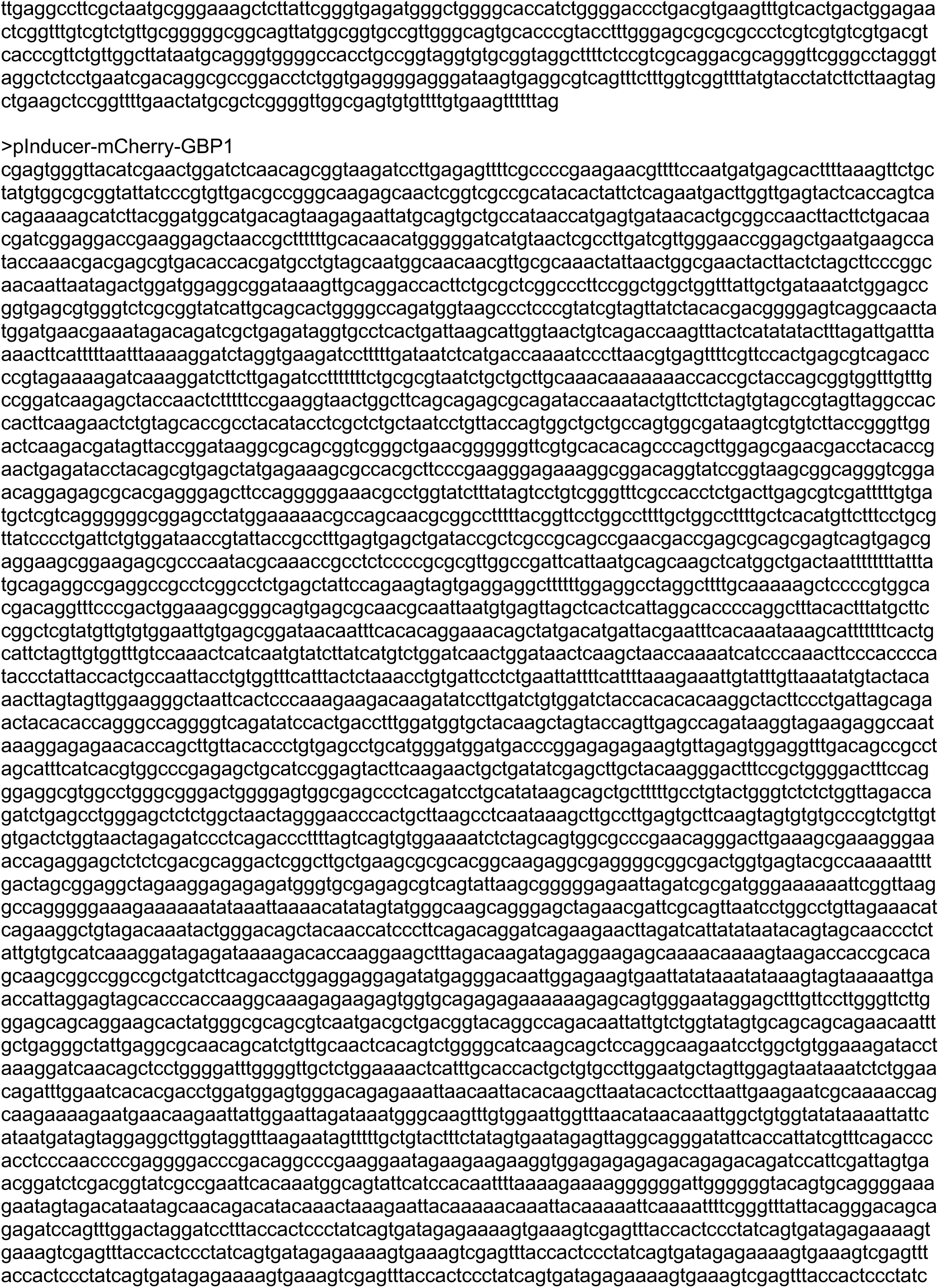
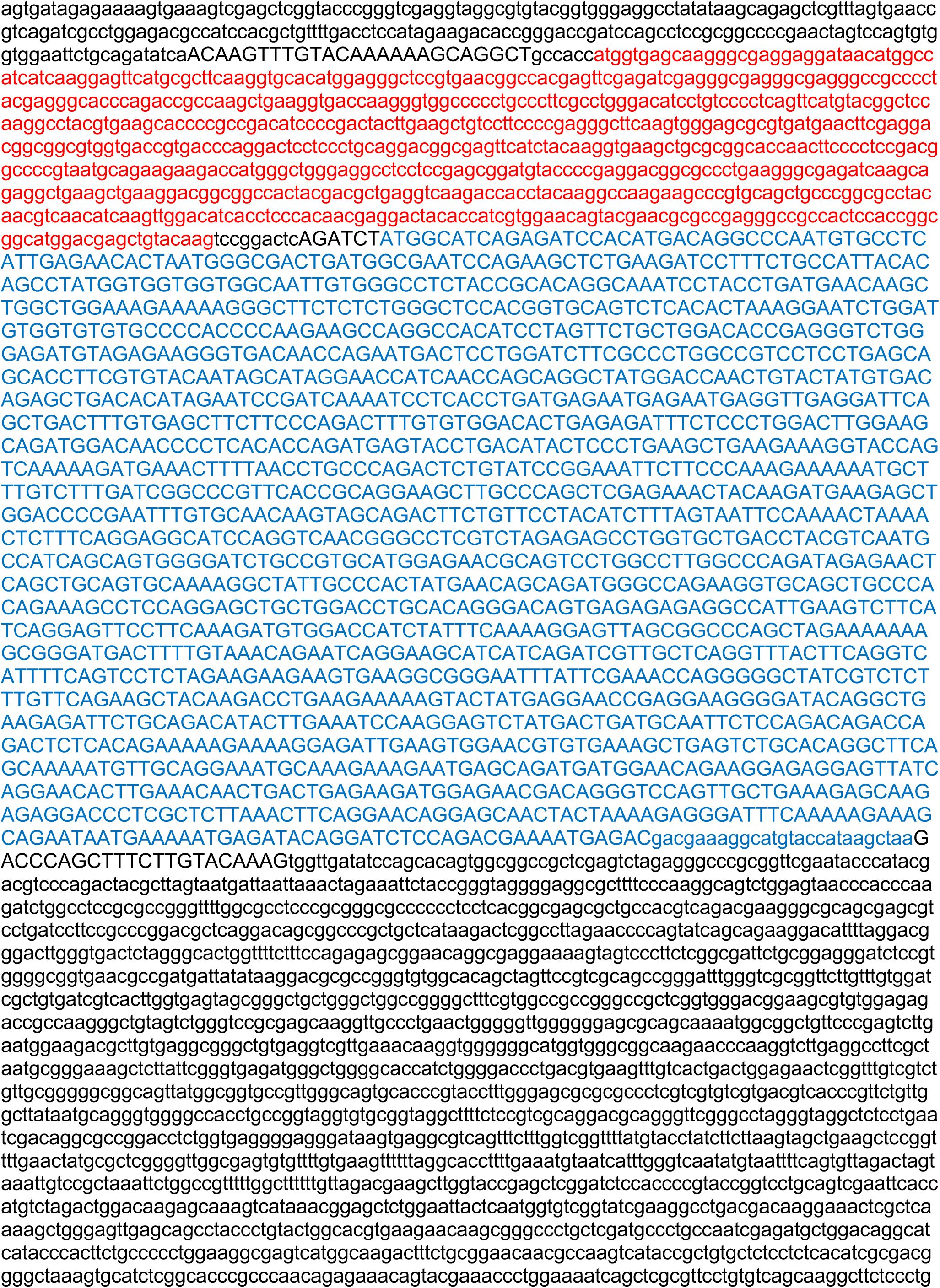
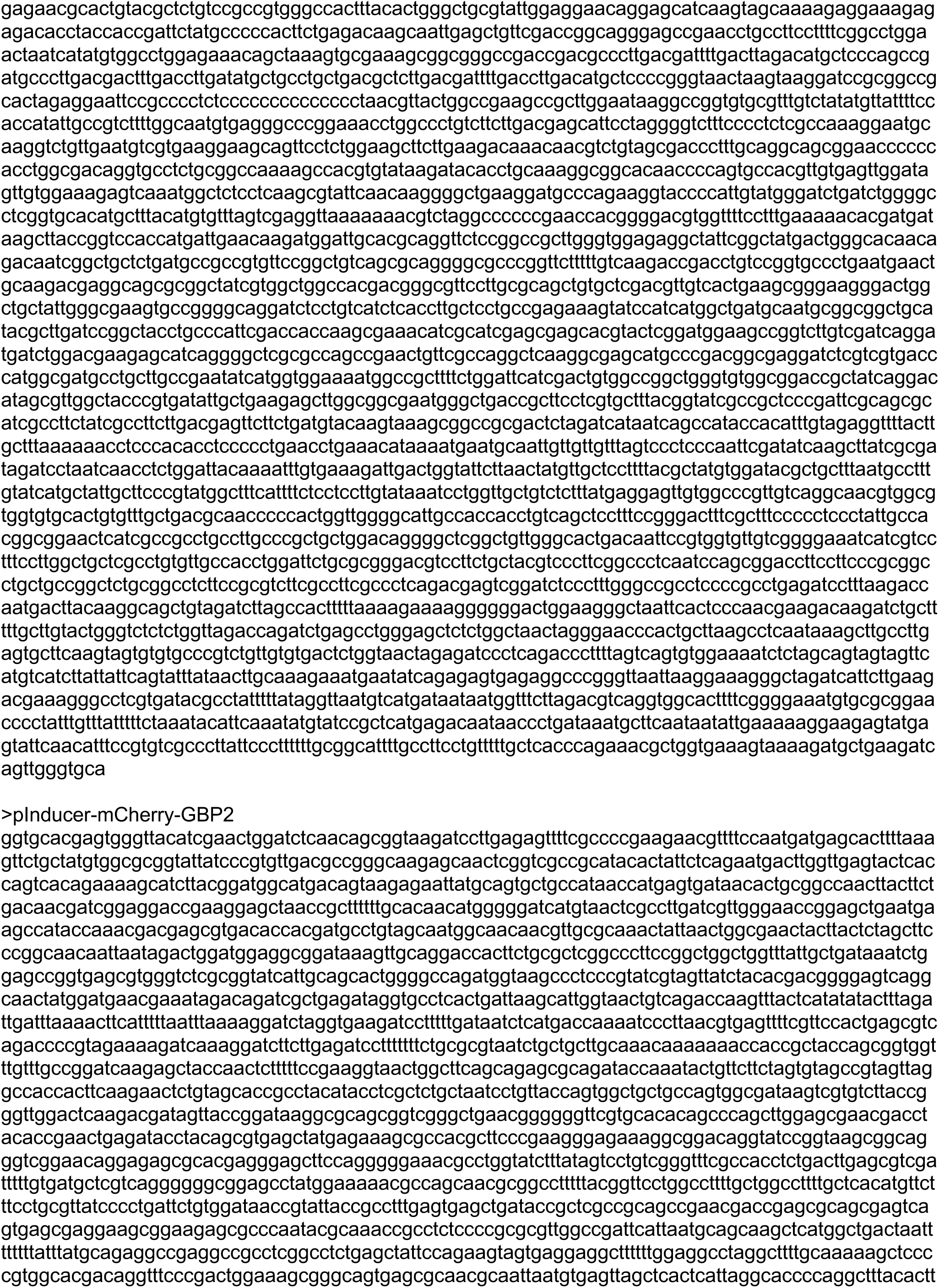
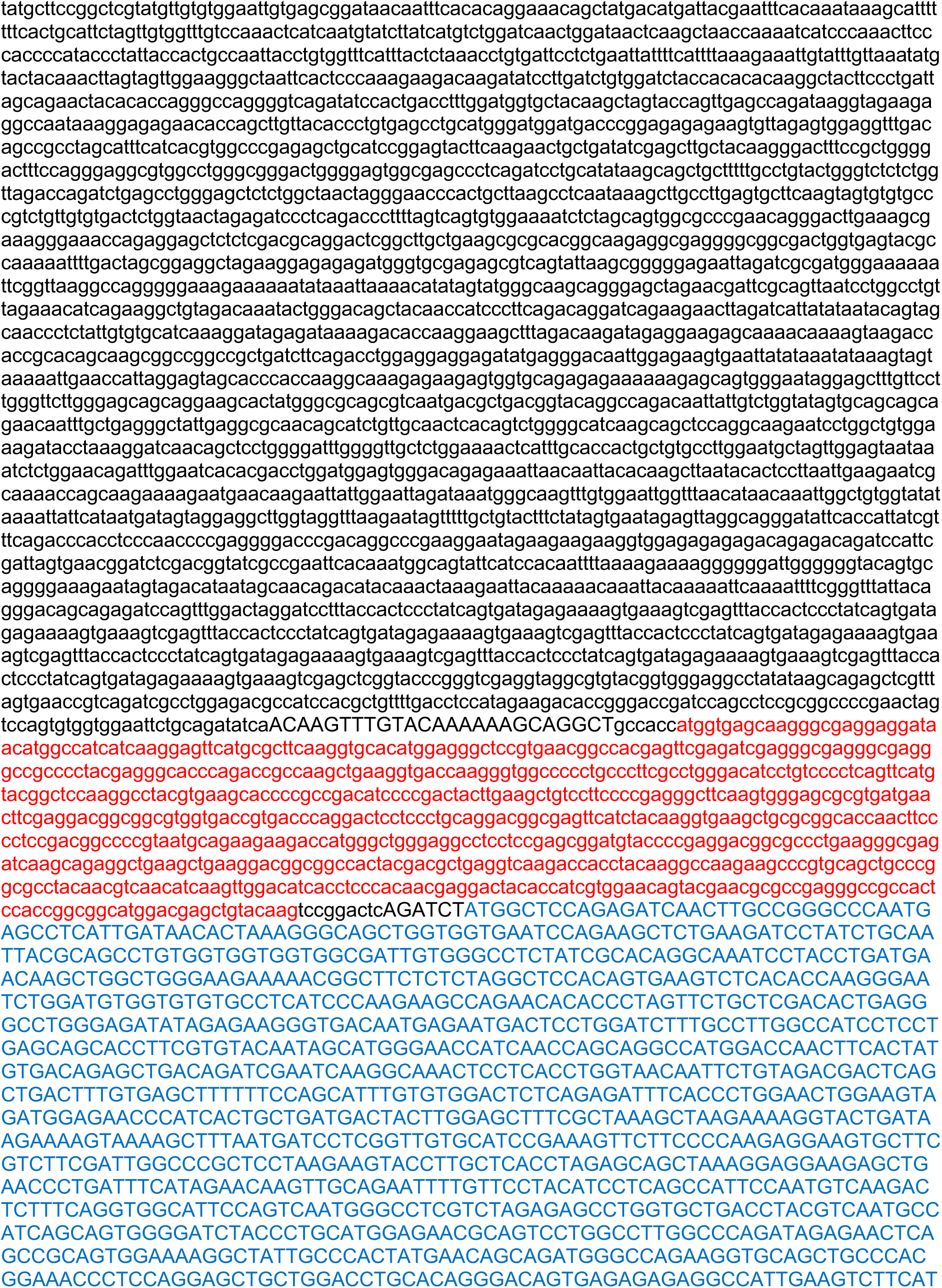
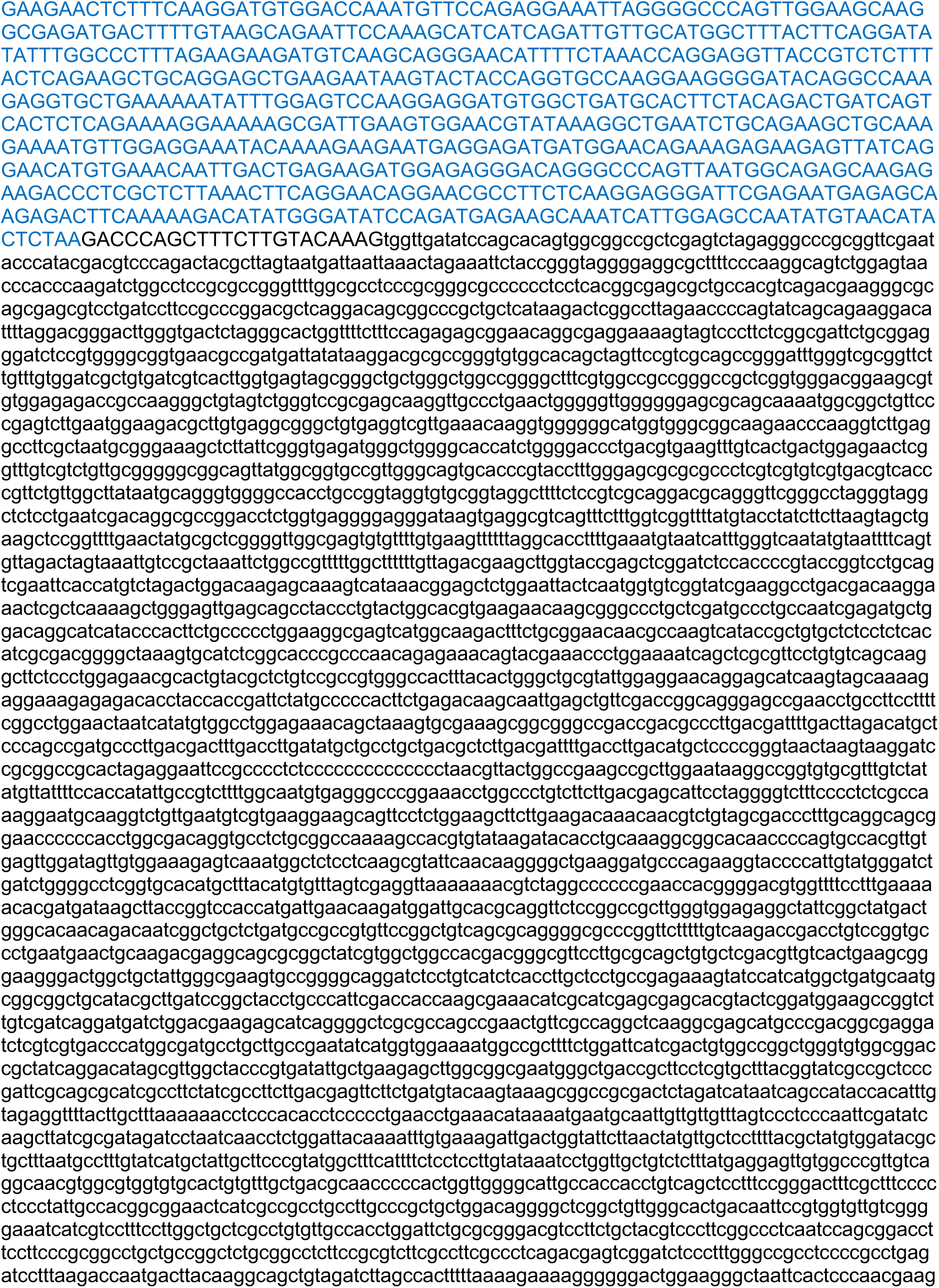
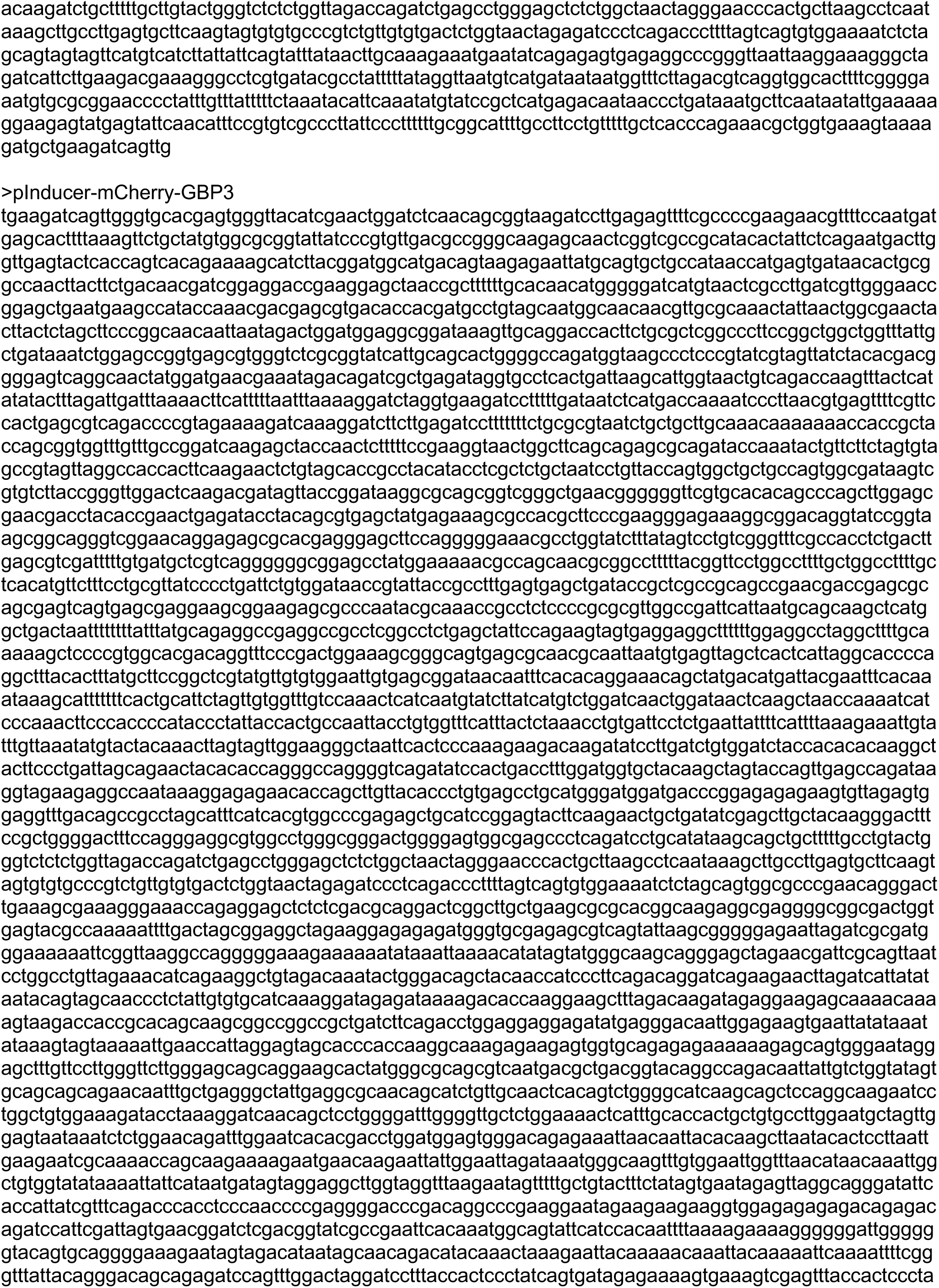
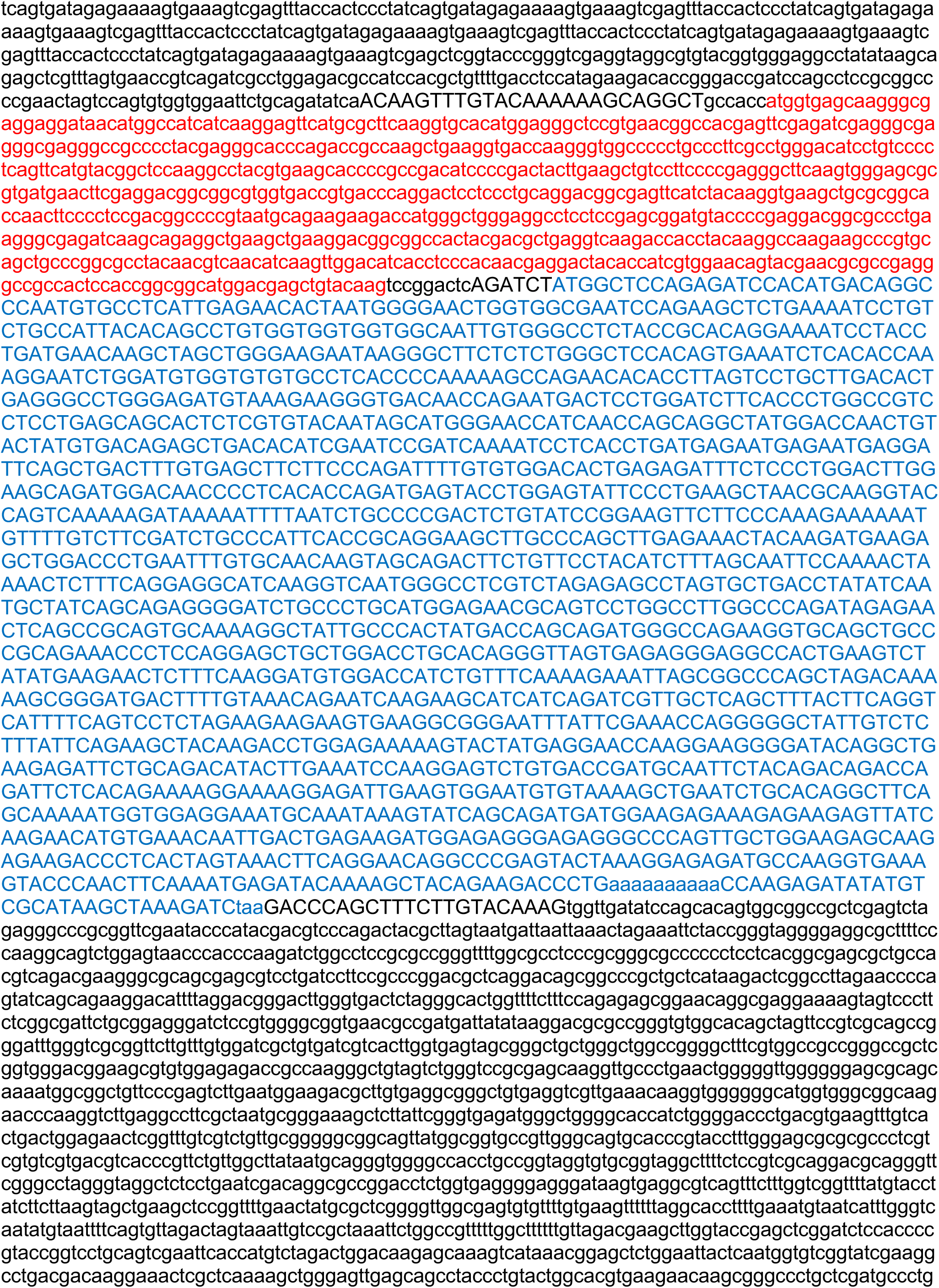
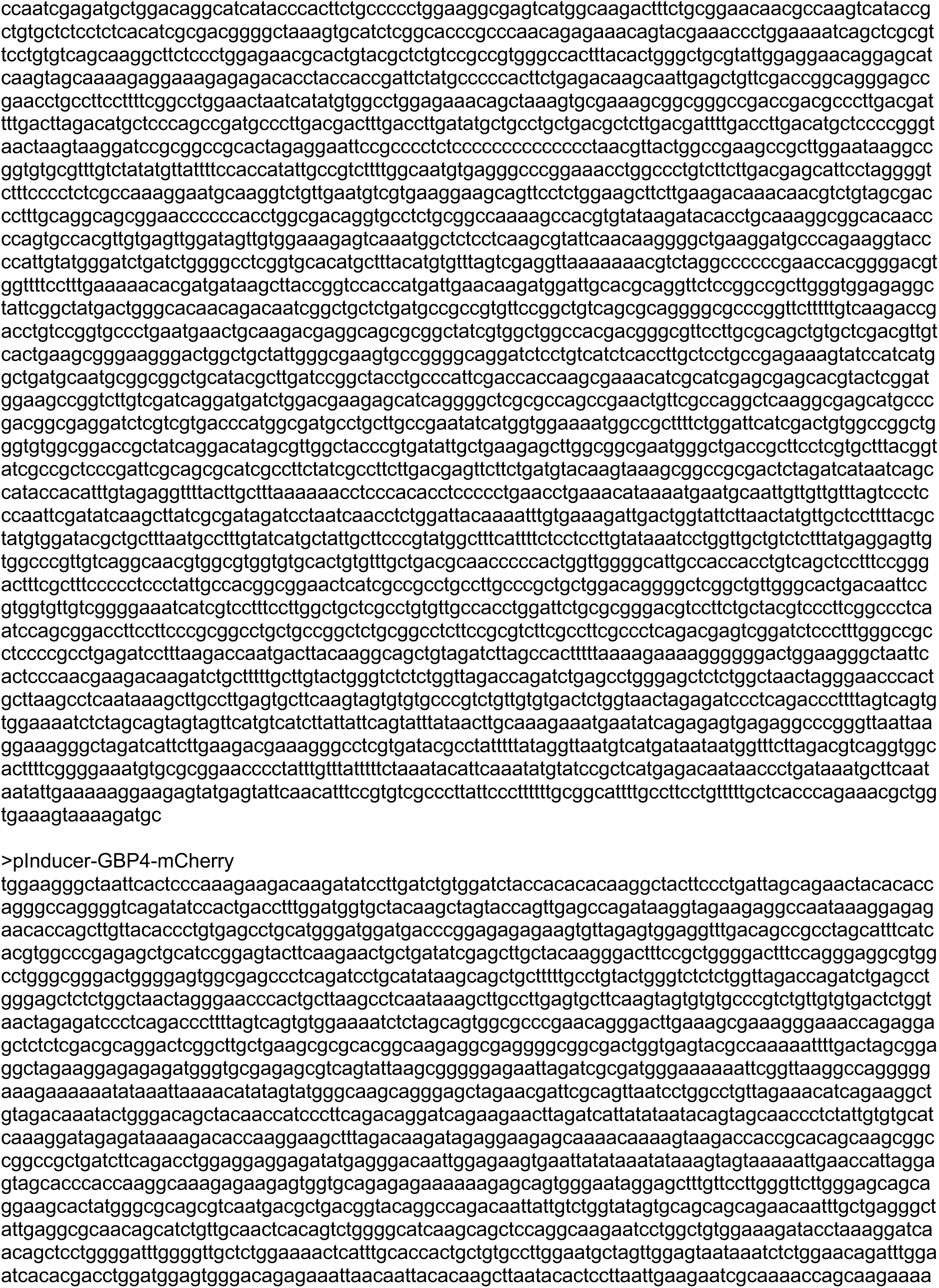
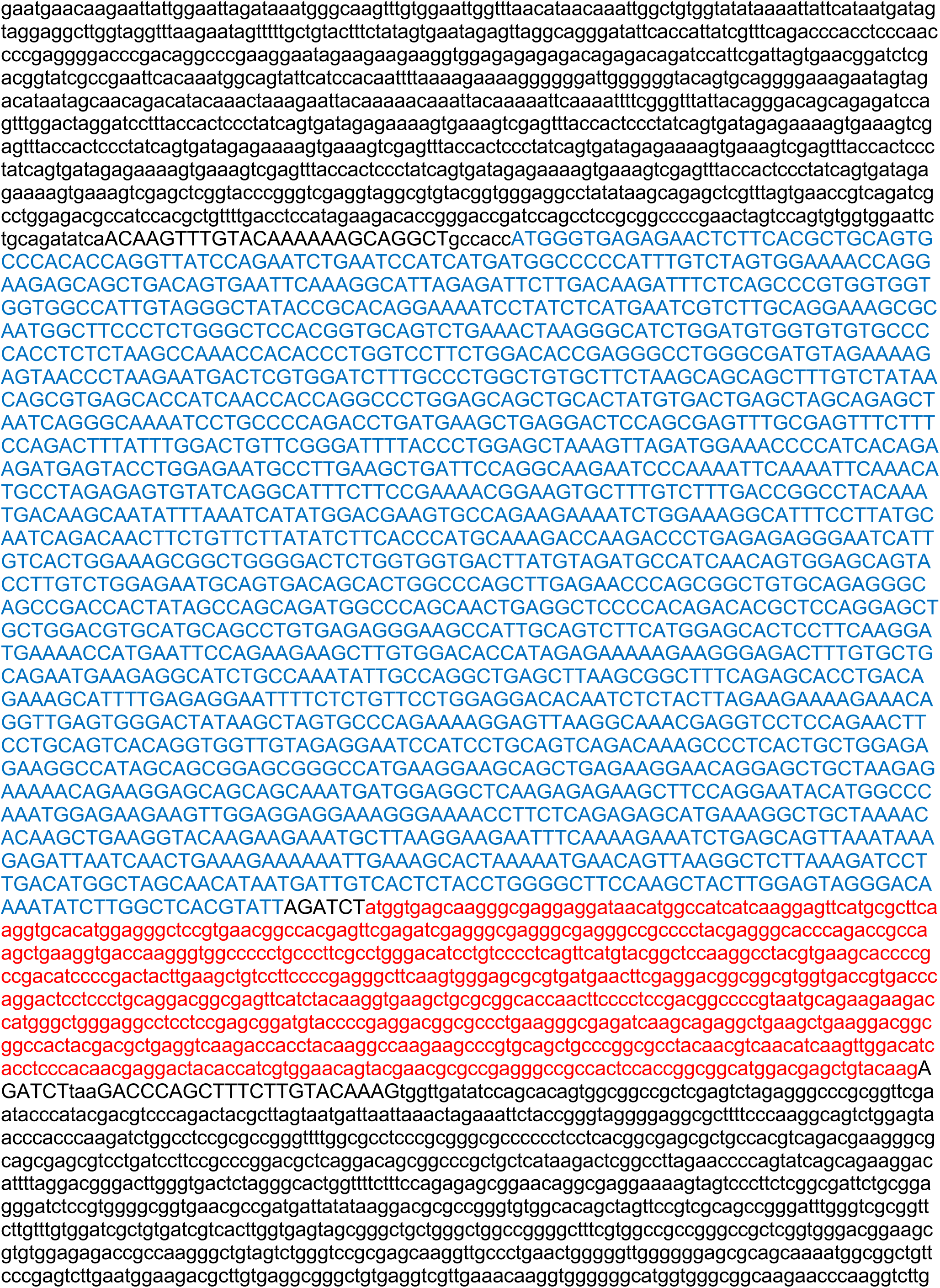
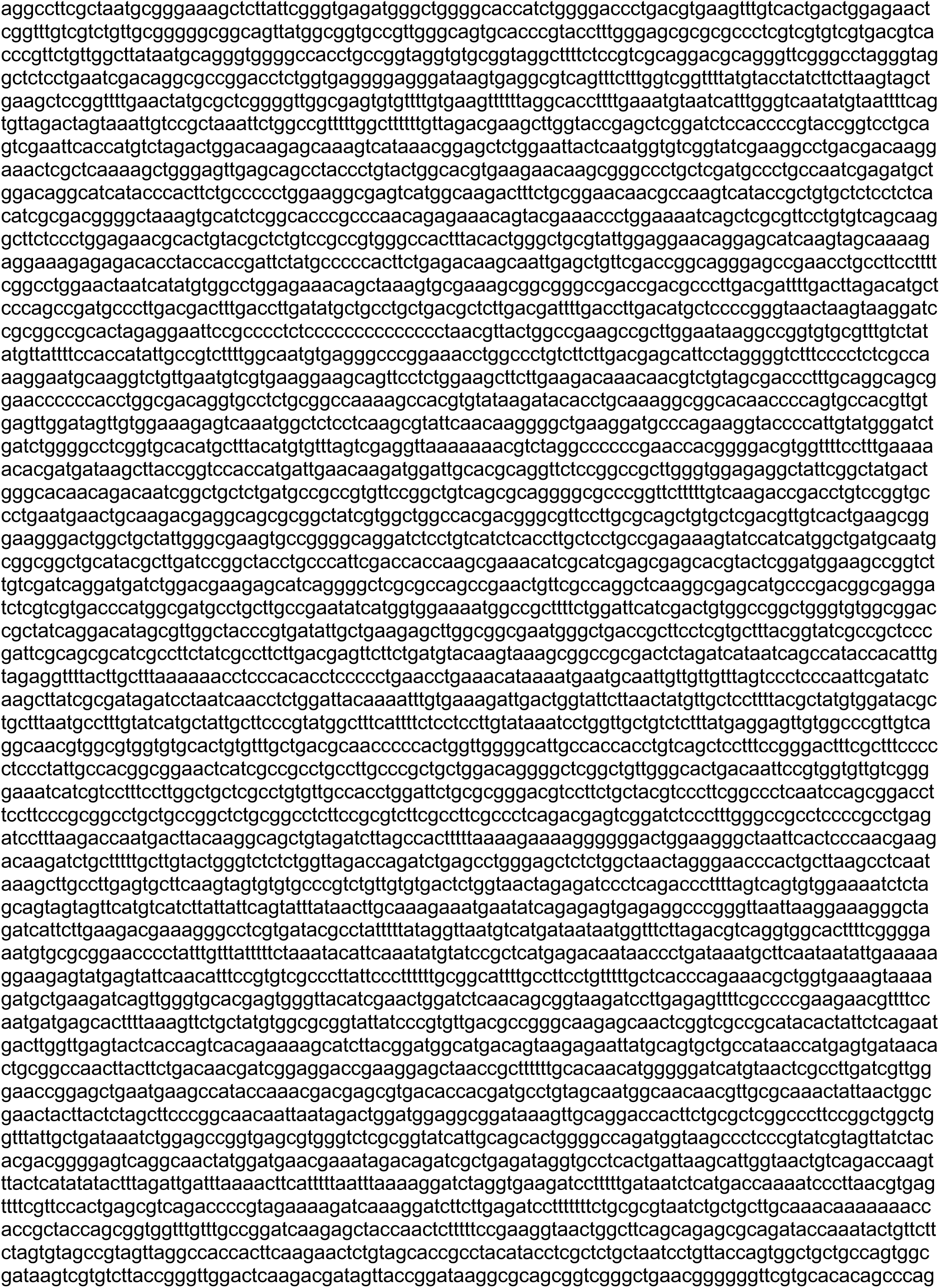
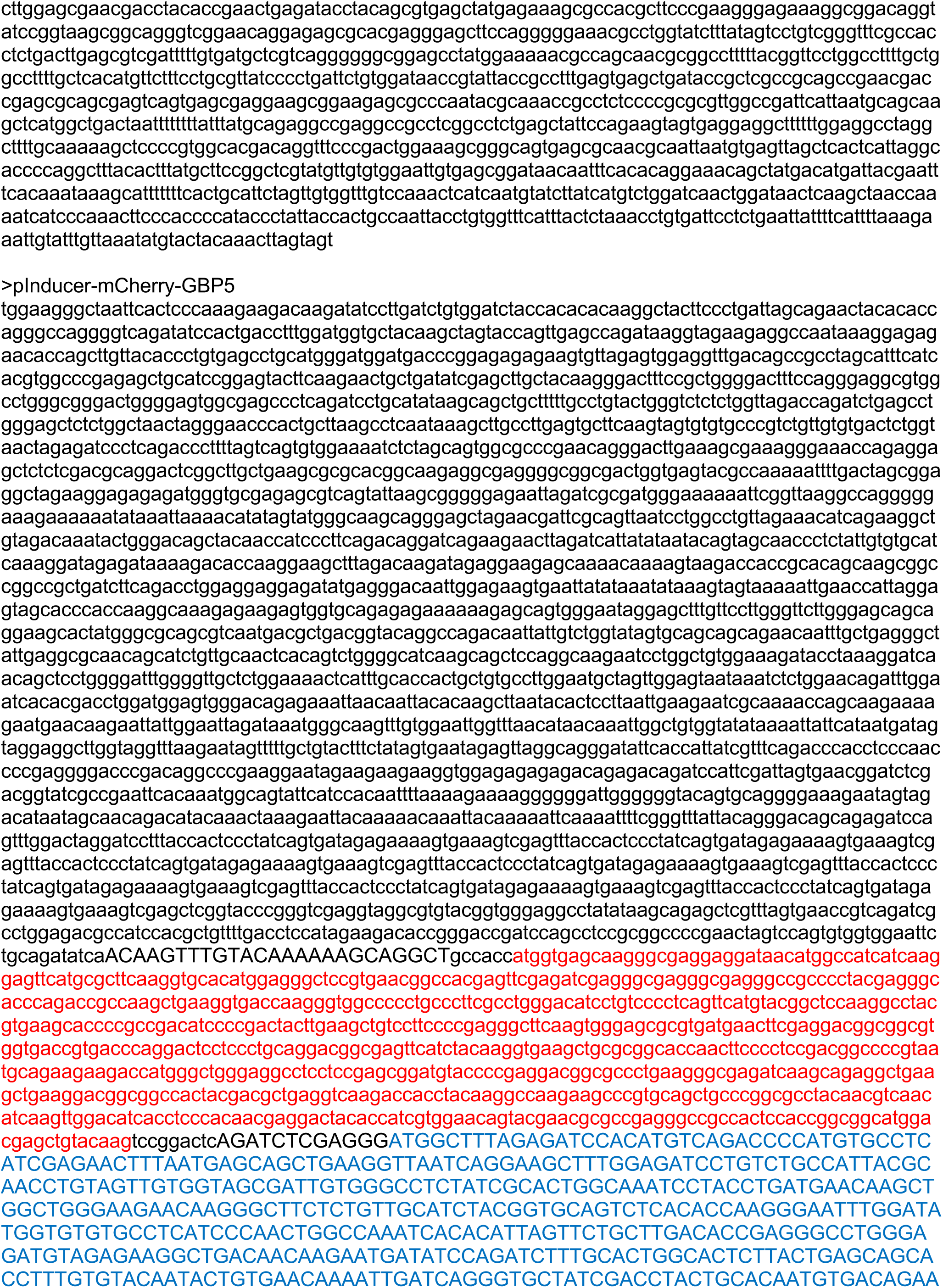
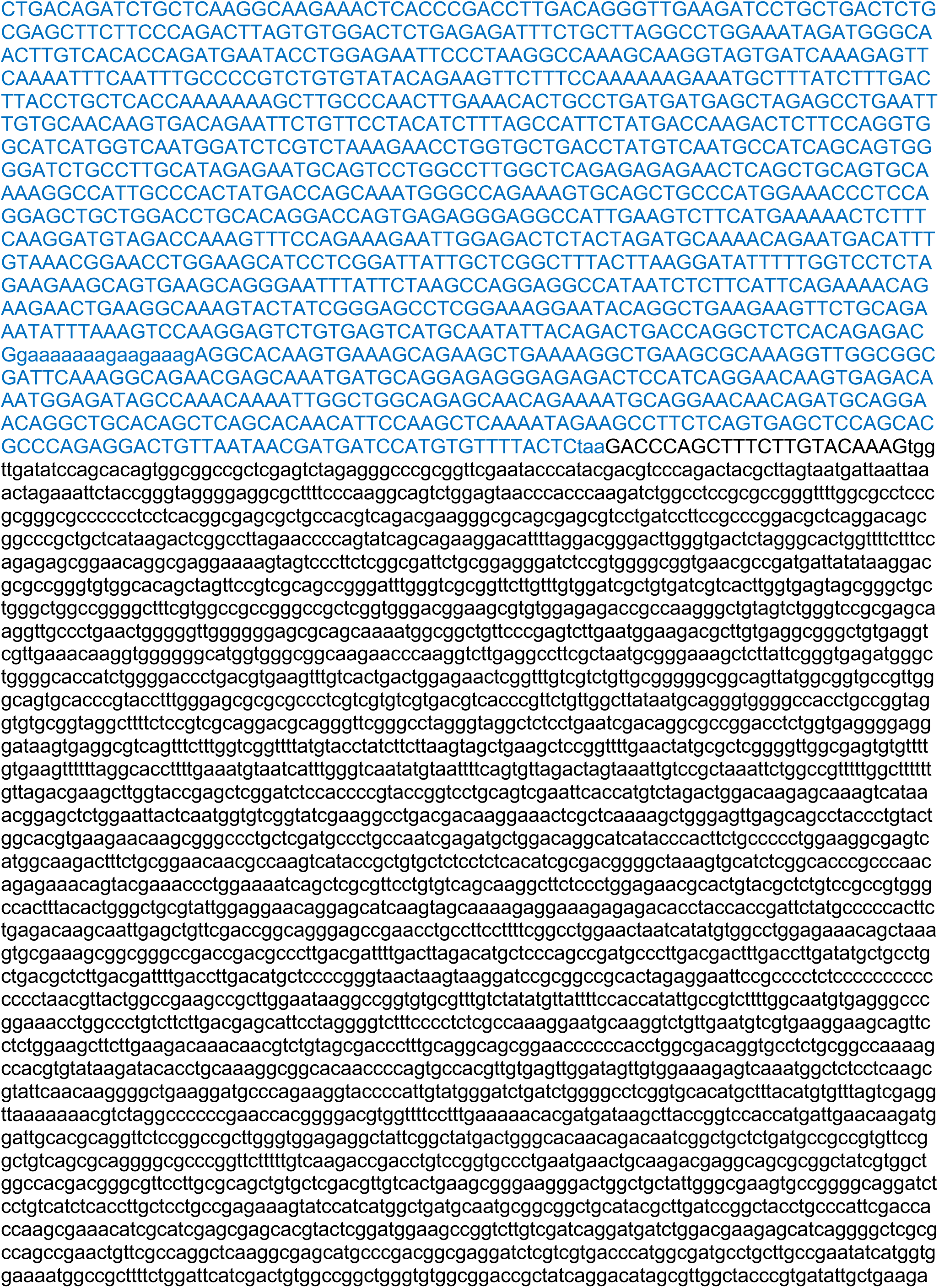
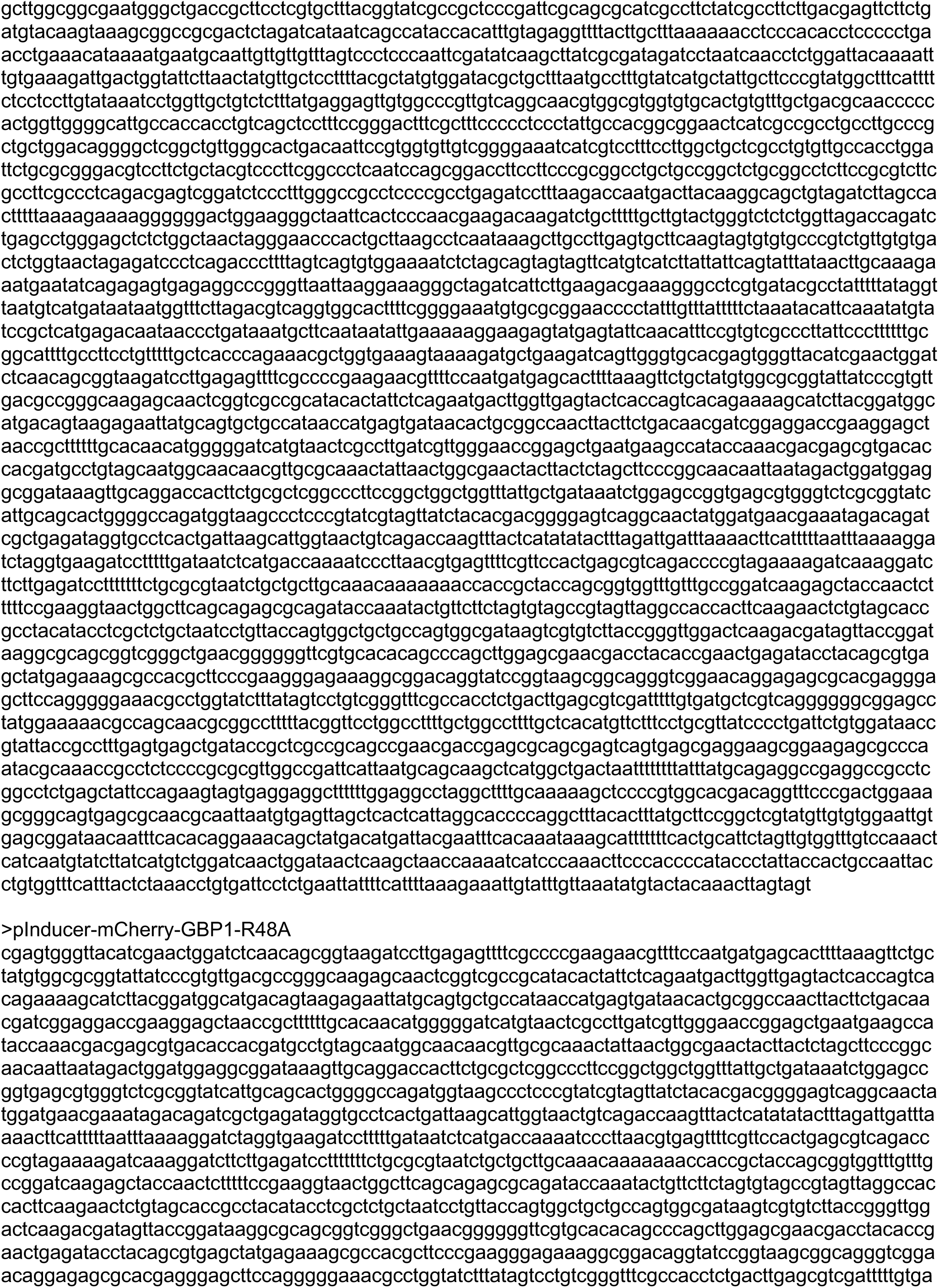
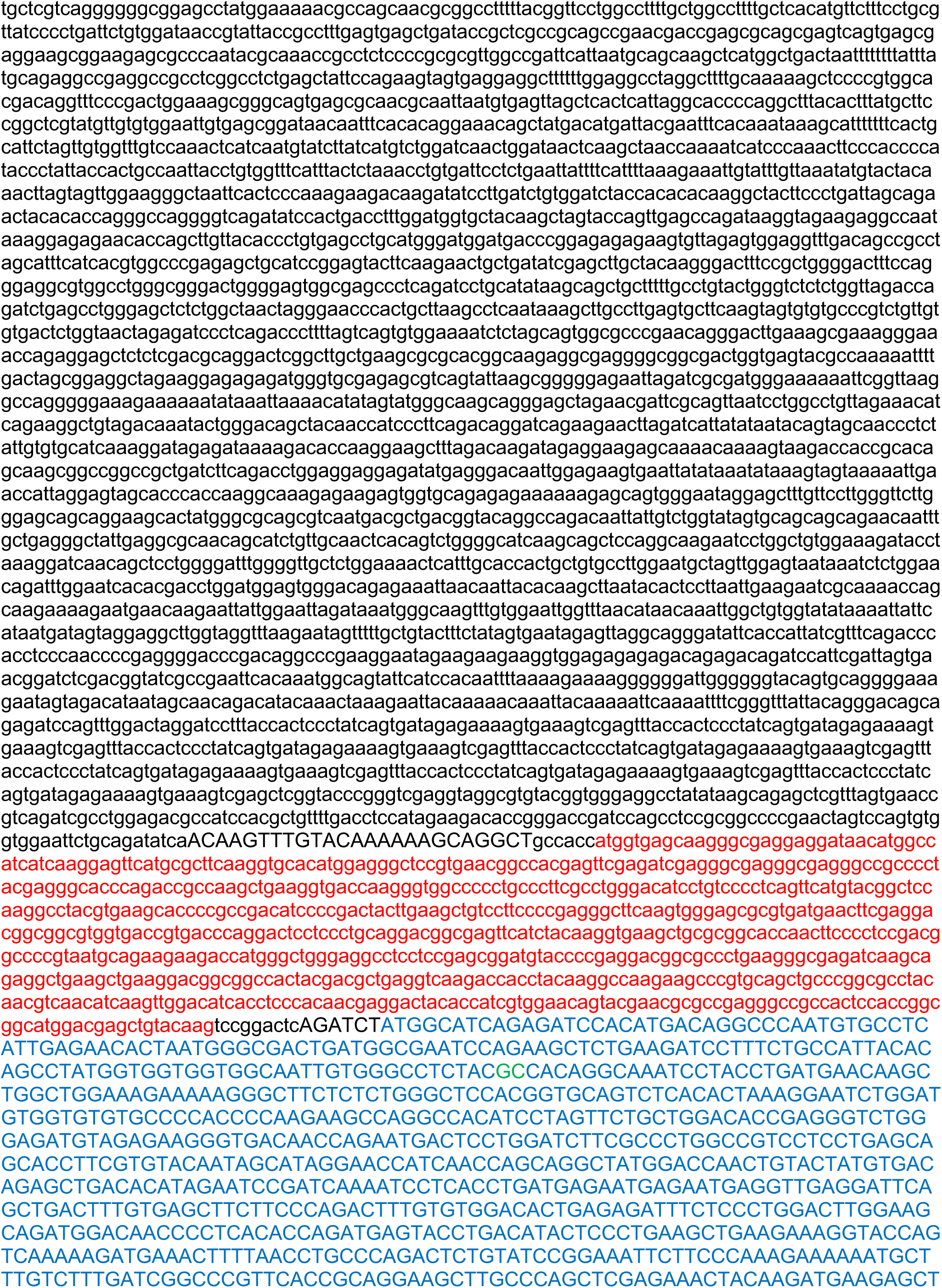
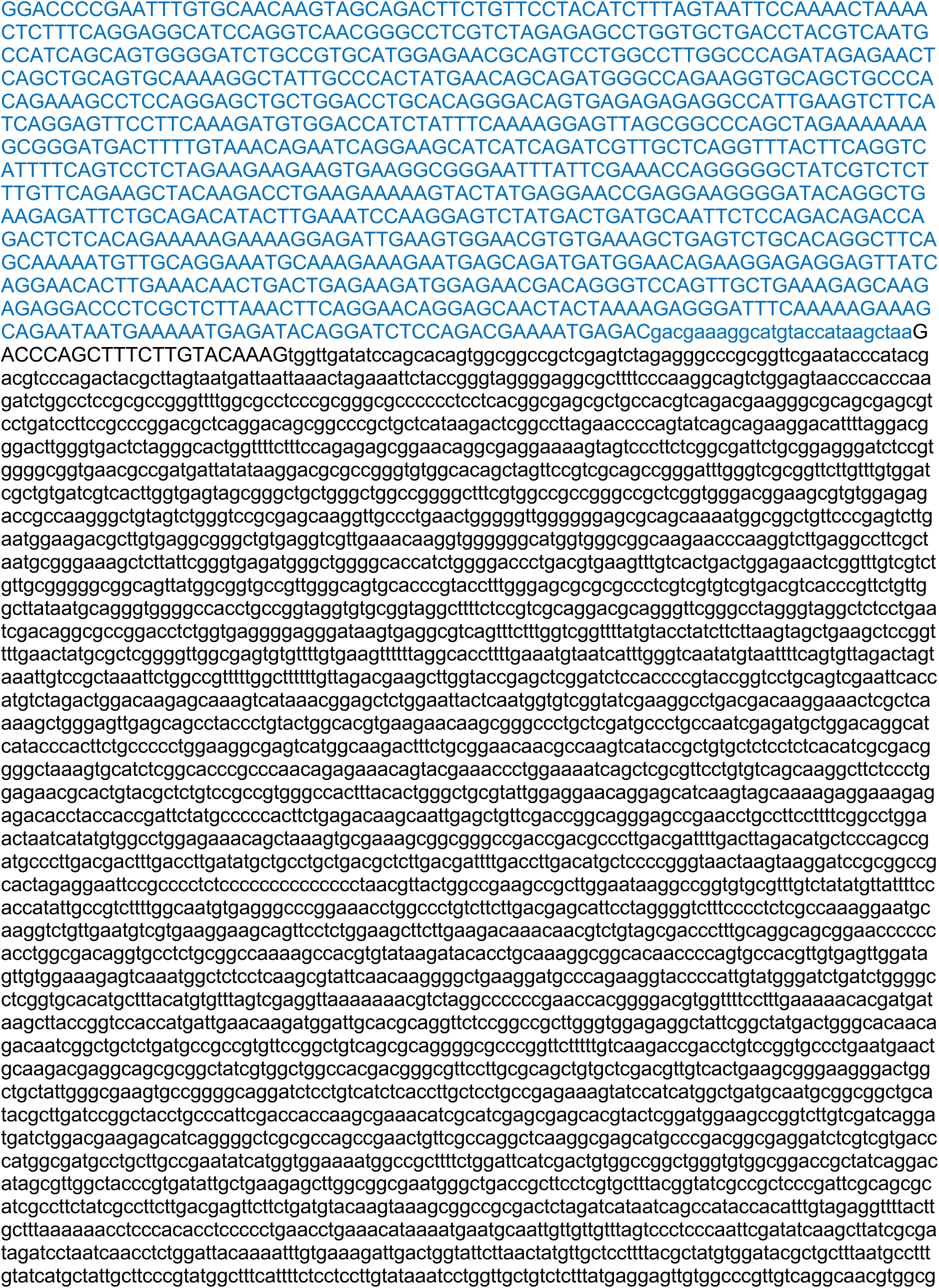
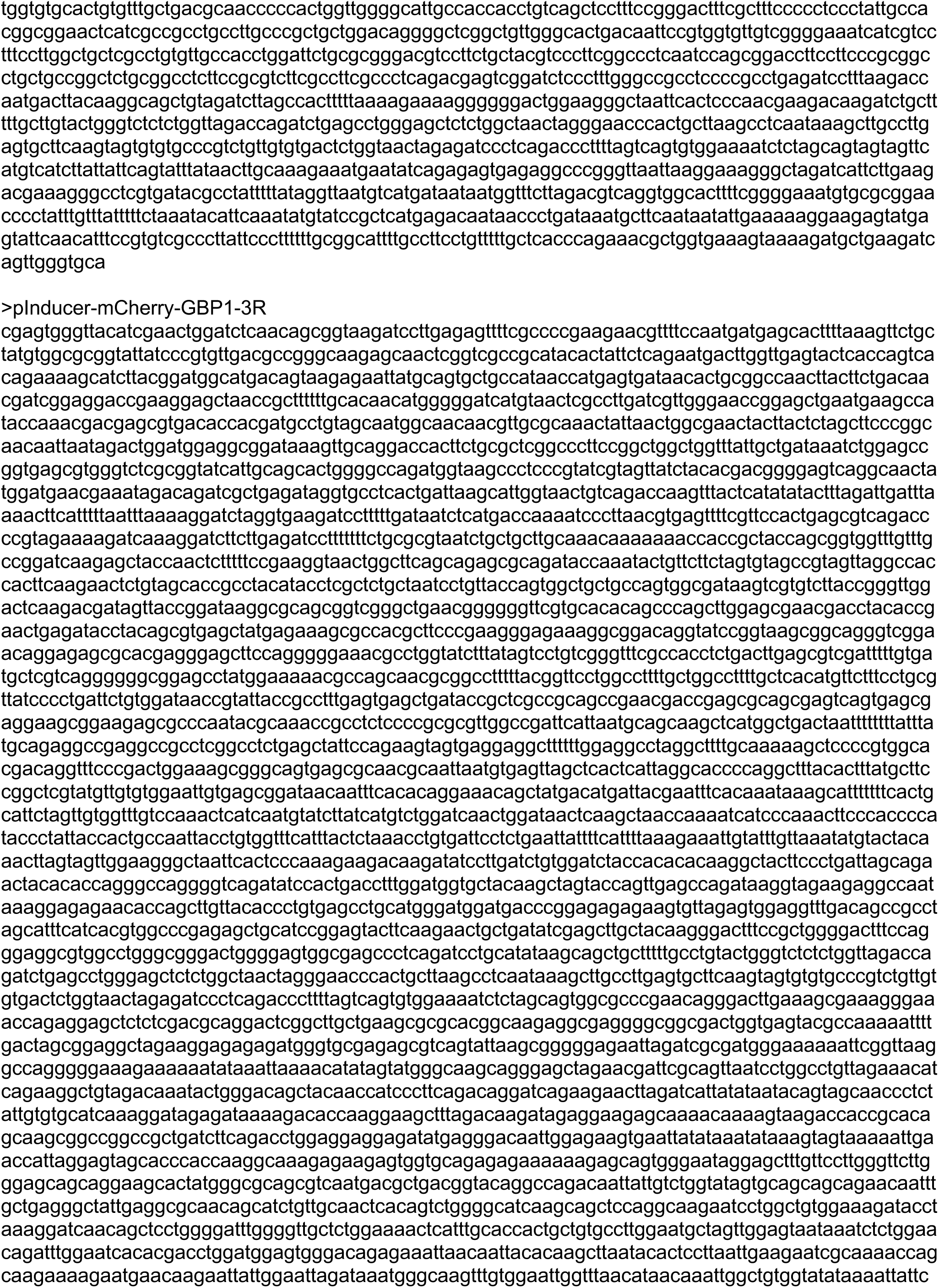
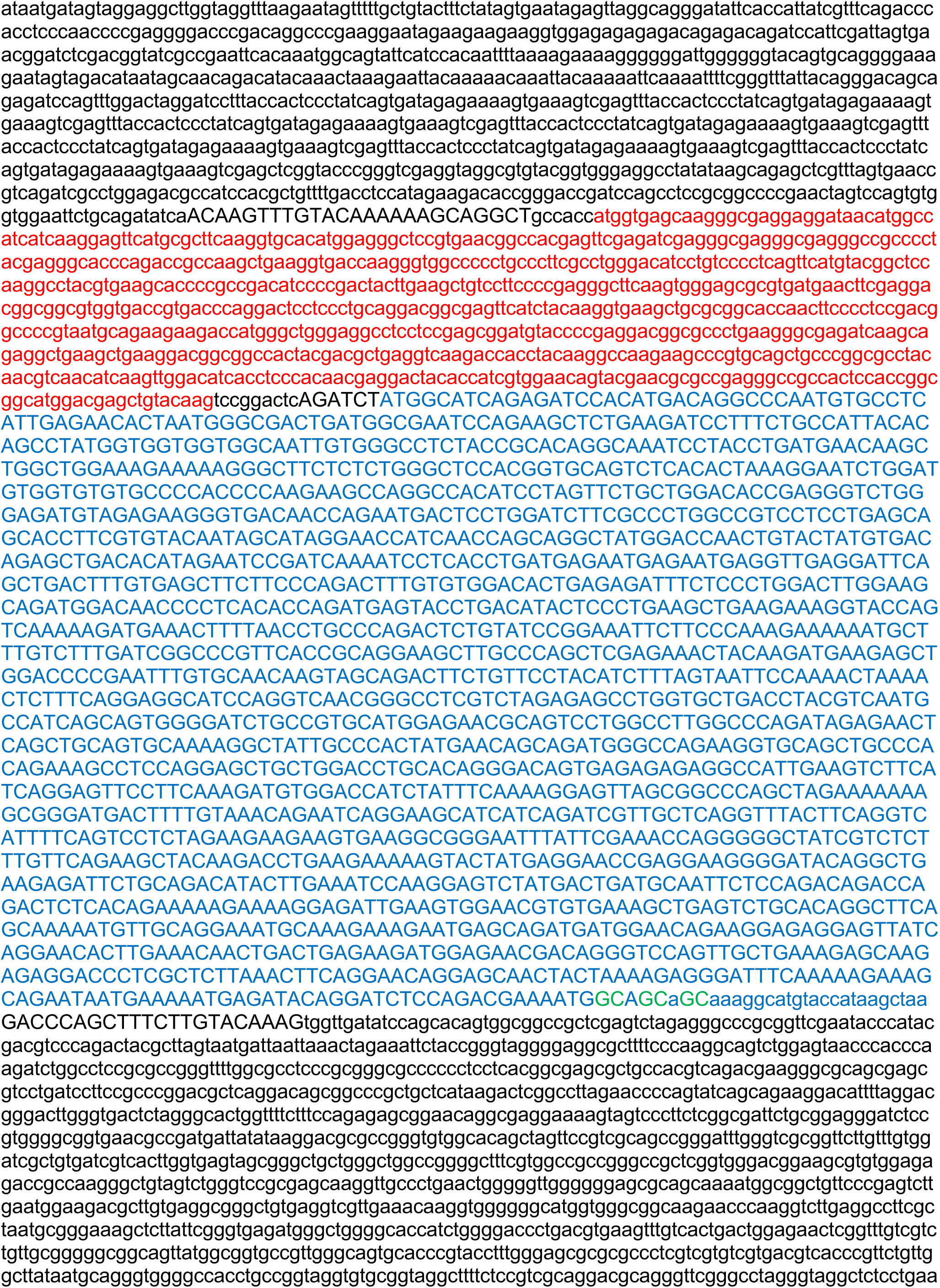
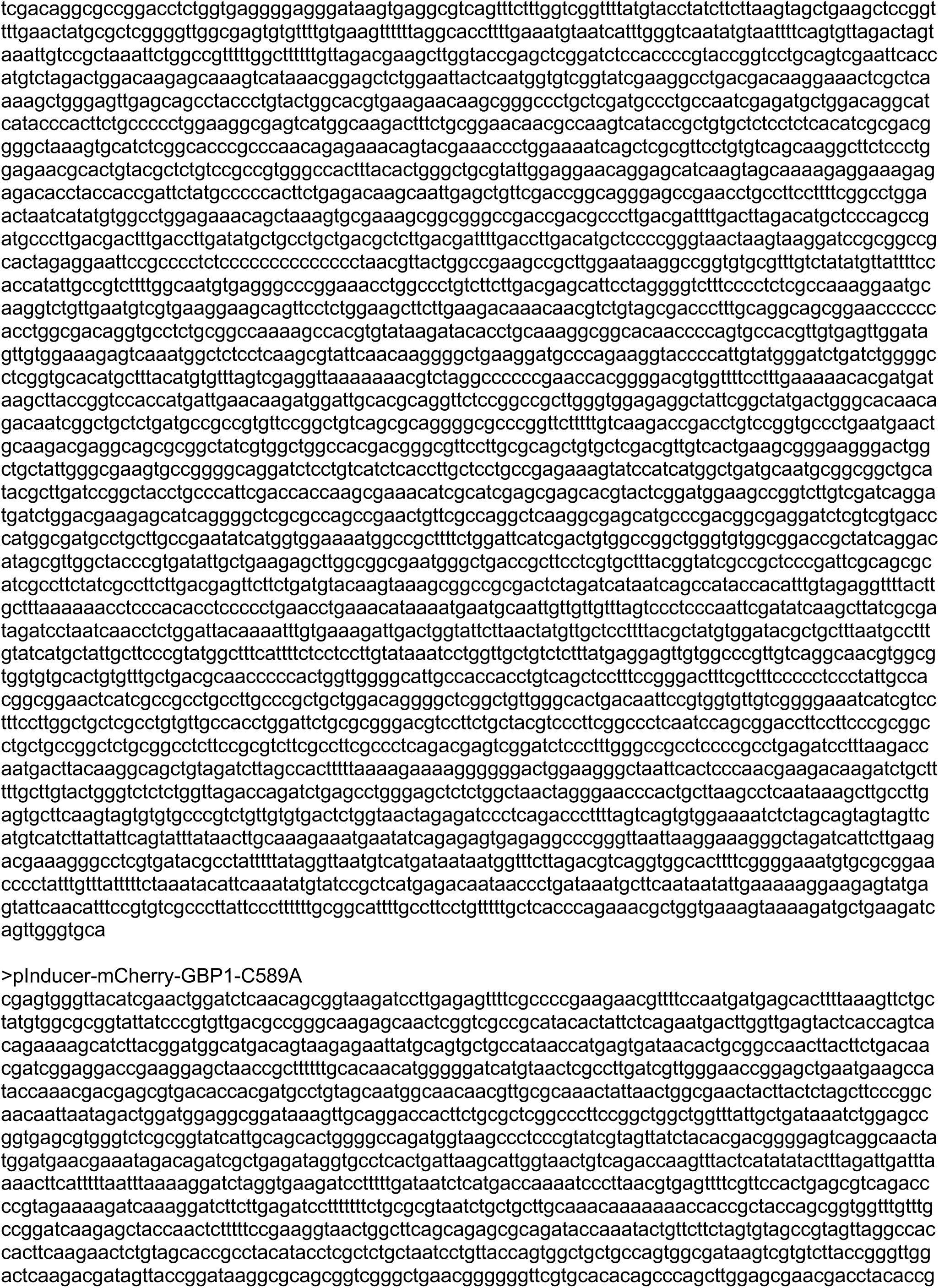
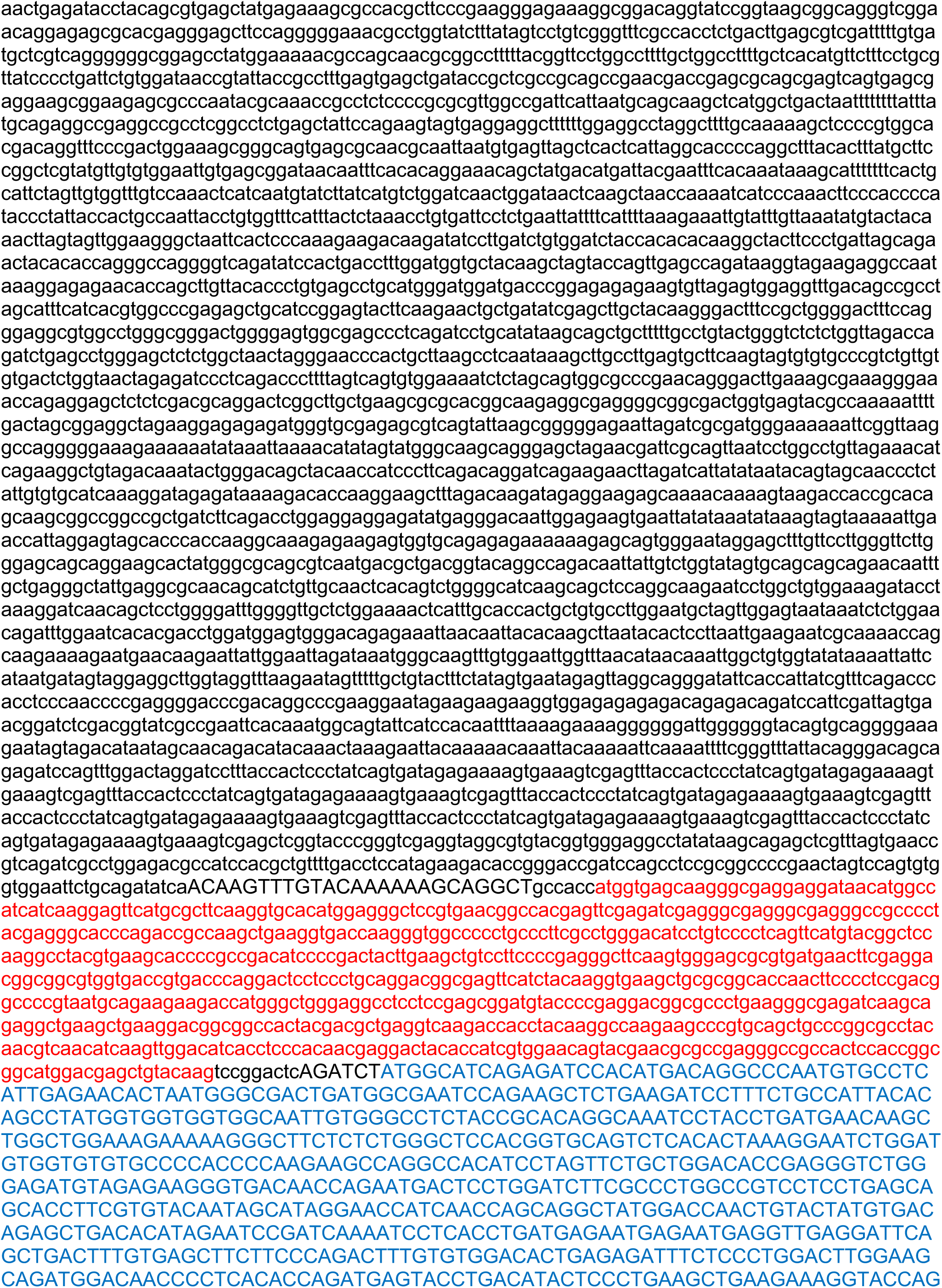
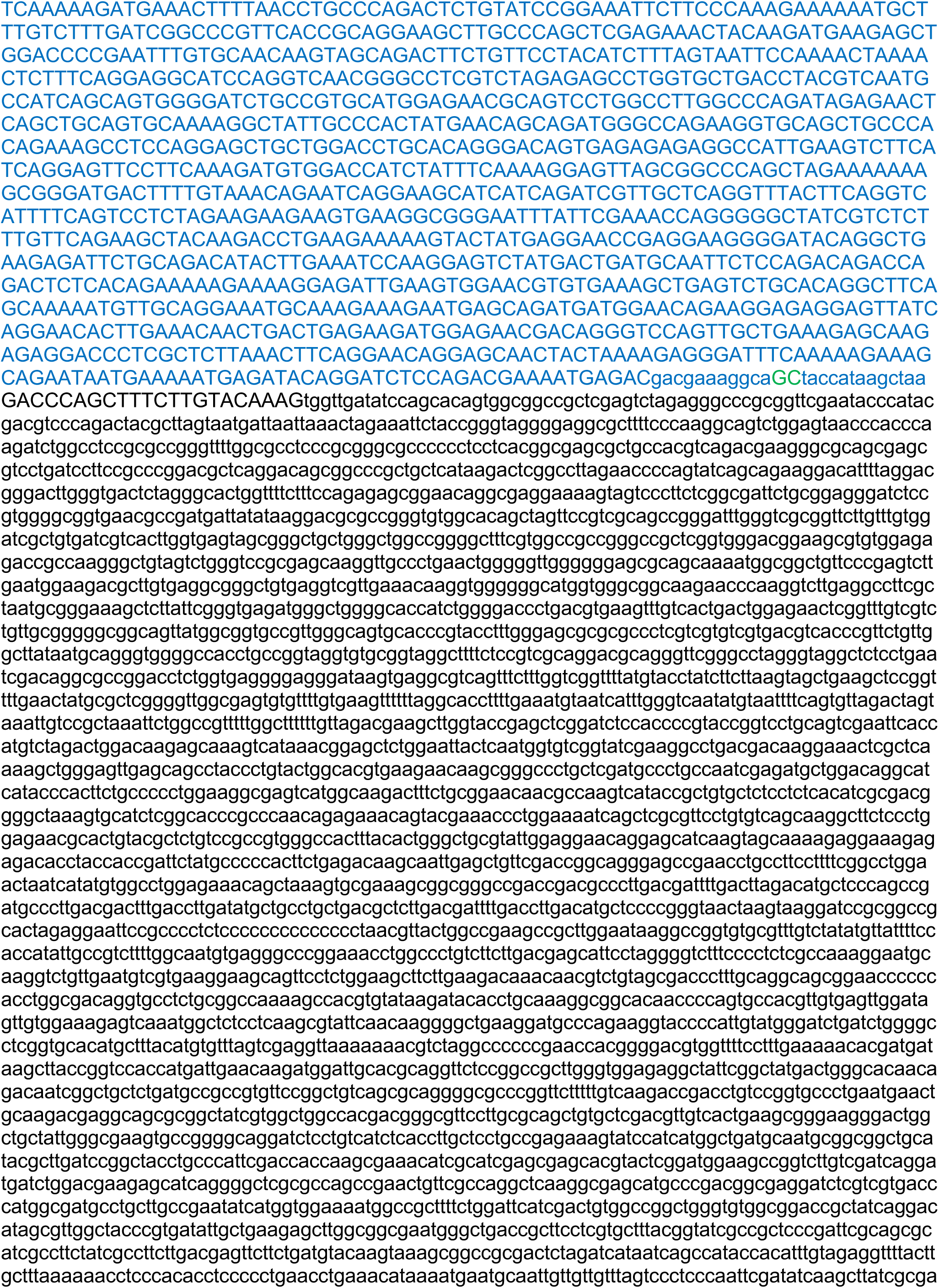
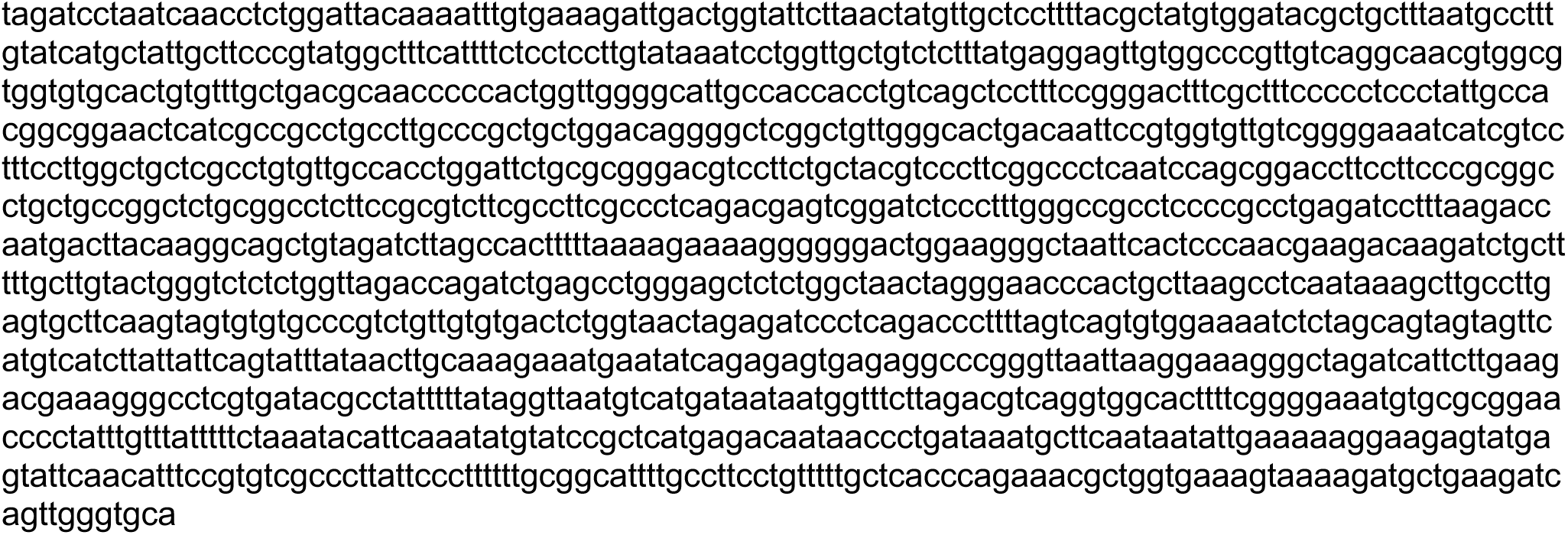

